# BRIDGE-AD reveals Alzheimer’s disease effectors through interpretable large-scale omics integration

**DOI:** 10.64898/2026.09.14.750802

**Authors:** Greta Baltusyte, Jonas Cerneckis, Harry Convey, Guoqiang Sun, Zyrille Chloe E. Abela, Miguel Ramirez, Daniel Wang, Guihua Sun, Tao Zhou, David Spring, Kourosh Saeb-Parsy, Yanhong Shi, Namshik Han

**Affiliations:** Milner Therapeutics Institute, University of Cambridge, Cambridge, UK; Department of Surgery, University of Cambridge, and Cambridge NIHR Biomedical Research Centre, Cambridge, UK; Cambridge Stem Cell Institute, University of Cambridge, Cambridge, UK; Cambridge Centre for AI in Medicine, University of Cambridge, Cambridge, UK; Department of Neurodegenerative Diseases, Beckman Research Institute of City of Hope, Duarte, CA, USA; Irell and Manella Graduate School of Biological Sciences, Duarte, CA, USA; Yusuf Hamied Department of Chemistry, University of Cambridge, Lensfield Road, Cambridge CB2 1EW, UK; Department of Biological Sciences, California State University Long Beach, Long Beach, CA, USA; Department of Quantum Information, Institute for Convergence Research and Education in Advanced Technology and Engineering, Yonsei University, Seoul, Republic of Korea; Department of Nano Biomedical Engineering (NanoBME), Advanced Science Institute, Yonsei University, Seoul, Republic of Korea; Center for Nanomedicine, Institute for Basic Science (IBS), Seoul, Republic of Korea; Department of Artificial Intelligence, Yonsei University, Seoul, Republic of Korea; College of Pharmacy, Yonsei University, Seoul, Republic of Korea

## Abstract

The growing landscape of Alzheimer’s disease (AD) datasets creates opportunities to integrate heterogeneous evidence and systematically discover disease effectors. We present BRIDGE-AD, an interpretable network medicine framework that transforms multimodal data into a unified, disease-specific gene representation for AD effector prioritisation. We integrated more than 30 datasets and curated resources spanning omics, functional, genetic and prior disease knowledge layers. BRIDGE-AD outperformed recently published pretrained and modality-specific gene embeddings in recovering AD-associated genes and produced a genome-wide resource of candidate AD effectors. Established and newly prioritised effectors formed 19 functional clusters, revealing a global molecular landscape of AD biology. BRIDGE-AD supported an SPP1-centred cross-compartment hypothesis and nominated SCARB2, a poorly characterised candidate, for functional validation. SCARB2 rewired lysosomal, lipid-handling and autophagic programmes in microglia, whereas disrupted SCARB2 glycosylation in AD implicated altered SCARB2 processing and function. The accompanying website, https://explore-bridgead.com, enables users to trace the curated evidence and generate mechanistic hypotheses.

## Introduction

Understanding the molecular basis of disease increasingly depends on integrating diverse forms of biological data, as different molecular profiling approaches offer complementary views of biological systems and can reveal emergent behaviours when considered together. However, differences in scale, format, and measurement type make their integration into unified models of disease biology challenging.

Gene representations provide a computational means of capturing functional relationships among genes and placing them within a broader biological context. These include embeddings derived from biomedical literature (BioConceptVec^1^ and GenePT^2^), protein-protein interaction topology (Mashup^3^), single-cell transcriptomes (scGPT^4^ and Geneformer^5^), bulk transcriptomes (FRoGS-ARCHS4^6^), and protein sequence data (protT5^7^). Although valuable, such representations are typically derived from a single data modality and are not designed to integrate the full range of available molecular, cellular, and functional evidence. Here, we address this gap by developing Biological Representation for Integrated Discovery of Gene Effectors in AD (BRIDGE-AD), an interpretable network medicine framework that transforms standardised multimodal evidence into a unified gene representation for genome-wide effector prioritisation. The underlying approach is applicable to any disease for which such evidence is available.

We apply the framework to Alzheimer’s disease, a progressive neurodegenerative disorder whose pathological and molecular complexity has motivated large-scale profiling initiatives such as AMP-AD^8^, AGORA^9^, and NIAGADS^10^. Nonetheless, these efforts have not yet translated into effective treatments, in part because the evidence remains fragmented. We therefore assembled more than 30 datasets and curated resources into a multimodal library spanning (i) bulk molecular profiles^11–18^, including transcriptomic, proteomic, metabolomic, and post-translational datasets capturing case-control differences, healthy aging, and AD progression; (ii) cell-resolved expression and regulatory profiles from single-cell (sc)– and single-nucleus (sn)-RNAseq and ATAC-seq datasets^19–22^; (iii) functional evidence from CRISPR screens^23–27^ and interaction proteomics^28–30^; and (iv) genetic^31–34^, pharmacological^35^, and prior disease knowledge^36,37^ resources. Each dataset was analysed to derive harmonised gene signatures, including differentially expressed genes, age– and biomarker-associated trajectory programmes, disease-enriched transcription factor regulons, functional screen hits, and genetically or pharmacologically implicated AD genes. These signatures were subsequently integrated into an AD-specific, network medicine-based gene embedding, where each gene in the human protein interactome was represented by interpretable features capturing its network proximity to the curated AD signatures. Using experimentally supported AD effector genes curated from the literature as a high confidence positive class, we trained machine learning models to distinguish these genes from the background and thereby identify previously unrecognised candidates. Across benchmark comparisons, the resulting AD-specific representation outperformed recently published alternatives in recovering known disease-associated genes.

Finally, we used BRIDGE-AD to translate gene prioritisation into biological hypotheses. High-confidence effectors recapitulated major themes in AD biology, including inflammation and lipid metabolism, while also highlighting poorly characterised genes. To demonstrate how this integrated evidence framework can support mechanistic interpretation, we developed a detailed, cross-tissue hypothesis linking SPP1 to brain-boundary immune activity, integrin-linked phagocytic programmes, and changes that occur during aging. We further investigated SCARB2, a highly prioritised but poorly characterised AD effector, and found that it induced a coordinated lysosomal, lipid-handling and autophagic state in microglia. In parallel, we identified extensive remodelling of SCARB2 glycosylation in AD, with potential consequences for SCARB2 maturation, endoplasmic reticulum retention, protein fate, and coupling to glucocerebrosidase. Collectively, this work establishes a structured framework for integrating heterogeneous datasets into an interpretable model of disease biology. To make BRIDGE-AD accessible as a discovery engine, we provide an accompanying platform at https://explore-bridgead.com that links prioritised effectors to the broad evidence space underlying their nomination. By allowing users to trace each prediction across these molecular, cellular, temporal and tissue contexts, BRIDGE-AD Explorer supports mechanistic hypothesis generation that extends beyond any single dataset or disease compartment.

## Results

### Curation of Class 1 effectors, assembly of the gene signature library, and construction of the AD-specific gene embedding

To define positive training labels for BRIDGE-AD, we curated a Class 1 reference set of 153 experimentally supported AD effector genes from the literature (**Figure 1A; Supplementary Table 1**). Class 1 genes were defined as those whose experimental manipulation using *in vitro* or *in vivo* models altered AD pathology, irrespective of whether the reported effect was protective or detrimental. We chose this approach to focus on genes with direct functional evidence in AD model systems, while treating association-based resources such as GWAS-derived candidates as complementary evidence rather than standalone labels. The Class 1 set was curated through a combination of manual review and programmatic literature screening (see **Methods**). Class 1 genes were assigned directionality (protective or detrimental) and structured evidence categories (*in vitro* and/or *in vivo* perturbations, neuropathology and/or behaviour outcomes, genetic support, and therapeutic tractability) based on the data used to curate them (**Figure 1A** & **Supplementary Table 1**). Canonical AD effectors, including ADAM10, APOE, APP, BACE1, MAPT, PSEN1, PSEN2, and TREM2, were also included in the Class 1 list but were not assigned to any evidence categories, as they were not sourced from specific publications.

**Figure 1.**
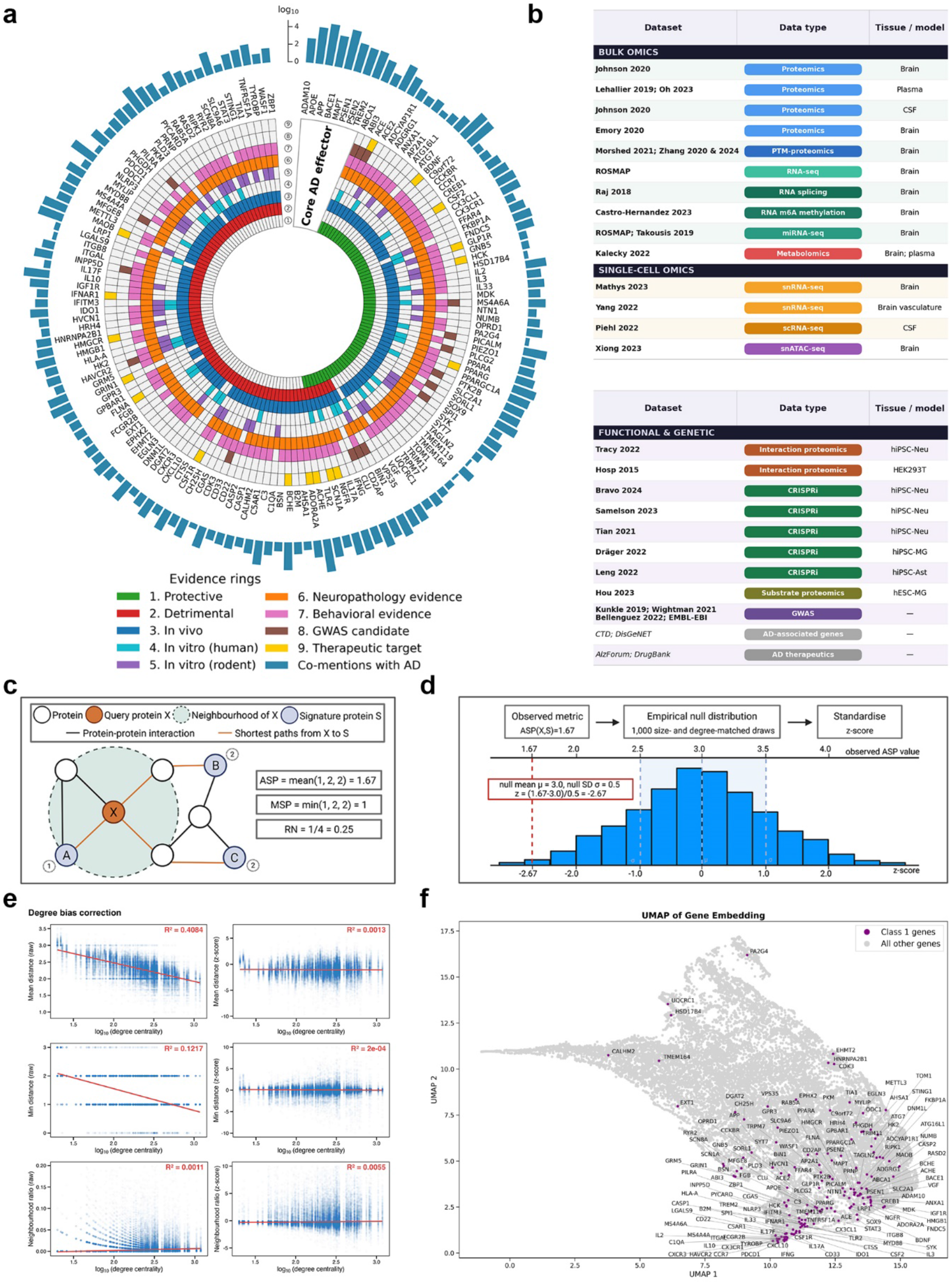
Class 1 effector curation, multimodal AD signature generation, and network-based feature construction for BRIDGE-AD. **(A)** An evidence map of Class 1 effectors used to create a positive label set for BRIDGE-AD. 153 Class 1 effectors were curated from literature and assigned disease directionality (protective or detrimental) as well as supporting evidence shown in coloured rings. Canonical AD effectors were also included without reference to specific publications or evidence types. The literature co-mention histogram reports log_10_-scaled PubMed abstract counts in which each Class 1 effector is mentioned alongside “Alzheimer’s disease” or “dementia” keywords. **(B)** Overview of datasets and resources analysed to construct the AD signature library. AD datasets include bulk multi-omics, single-cell and single-nucleus atlases, genetic studies, and functional knowledge resources. **(C)** Network proximity features computed for each STRING PPI network node relative to each AD signature: average shortest-path distance (ASP), minimum shortest-path distance (MSP), and ratio of immediate neighbours (RN) that belong to the signature. **(D)** Schematic of degree-matched proximity normalisation. For each protein-signature proximity metric, the observed value was compared with a null distribution generated from size– and degree-matched random gene sets. The resulting z-score indicates whether the observed proximity is greater or smaller than expected by chance. **(E)** Diagnostic assessment of degree bias normalisation. Plots show the relationship between Class 1 gene degree centrality and raw or z-score-normalised proximity metrics, with lower R² values indicating reduced degree dependence after normalisation. **(F)** UMAP visualisation of the resulting gene-feature space, showing the distribution of Class 1 AD effectors.

Overall, 81 Class 1 effectors were annotated as detrimental-only, 56 as protective-only, and 8 as having both protective and detrimental effects, reflecting the literature’s stronger emphasis on pathogenic drivers while still capturing protective candidates. Most Class 1 effectors were supported by neuropathological readouts (137/153; 89.5%) and behavioural or cognitive outcomes (112/153; 73.2%), indicating their functional relevance to AD pathogenesis. We also quantified PubMed abstract co-mentions between each Class 1 effector and AD (see **Methods**), confirming that biomedical literature is skewed towards a small subset of central AD genes, particularly the canonical AD effector group (**Figure 1A**, outer histogram). To further characterise the Class 1 list, we assigned dominant pathways and primary cell types to each of the effectors based on the studies used for curation. We found that Class 1 effectors represented common themes in AD, including proteostasis and endolysosomal trafficking (51/153; 33.3%), neuroimmune interactions and inflammation (49/153; 32.0%), and amyloid processing (43/153; 28.1%) (**Figure S1**). Primary cell type annotation indicated that neuron– (70/153; 45.8%) and microglia-associated (68/153; 44.4%) effectors were the most abundant, consistent with classical neuronal pathology in AD and a more recent focus on microglial inflammatory and phagocytic programmes (**Supplementary Table 2**). Finally, Gene Ontology (GO) biological process enrichment analysis revealed that most enriched terms contained both protective and detrimental AD effectors (**Extended Data Figure 1**), suggesting that a gene’s effect on AD pathogenesis is determined not by pathway membership alone, but by direction and context of pathway perturbation.

Next, to assemble the AD signature library, we analysed a diverse collection of omics datasets and curated AD resources (**Figures 1B** and **S2**). We first derived case-control signatures from bulk omics datasets spanning brain^15,38^ and CSF^11^ proteomics, brain post-translational modification proteomics^12–14^, RNA-seq^39–41^, and miRNA-seq^42,43^ (**Figure S3A**; for complete signatures, see **Data Availability**). For differentially expressed miRNAs, we generated target-gene signatures by mapping predicted targets using miRDB^44^ and TargetScan^45^ databases. We also analysed AD-associated metabolomic changes in brain and plasma tissues and used these differences to infer altered enzyme activities^46^ (**Figure S3B–C**). Since the CSF proteome dataset^11^ used for diagnostic case-control analysis also included quantitative AD biomarker measurements, such as pTau and Aβ_42_, we further modelled protein-level trajectories along disease severity and clustered significantly altered trajectories into gene signatures (**Figure S4A**). In addition, given the strong link between aging and AD, we analysed healthy brain^11^ and plasma^17,18^ proteomes to identify age-associated protein abundance trajectories (**Figure S4B–C**).

To derive cell type-specific signatures, we analysed four single-cell/nucleus atlases of AD gene expression and chromatin accessibility. From brain parenchyma snRNA-seq datasets^19^, brain vasculature snRNA-seq^22^, and CSF scRNA-seq^20^, we generated healthy cell type marker signatures and cell type-specific AD versus control differential expression signatures (**Figure S5**). The CSF dataset also included young and aged healthy donors, allowing us to define age-associated transcriptional changes across cell types. Meanwhile, we used the brain parenchyma snRNA-seq dataset to infer AD-associated cell type-specific interactome rewiring using Single-Cell Imputation and NETwork construction (SCINET, see **Supplementary Methods**)^47^. From the matched brain snRNA-seq/snATAC-seq dataset^21^, we further derived cell type-specific markers of gene accessibility and transcription factor activity in healthy cells, as well as AD-associated changes in these regulatory features (**Figure S6**). Finally, we incorporated signatures from previously published AD-relevant CRISPRi screens (**Figure S7**)^23–27^, interaction proteomics studies^28–30^, GWAS results^31,32,34,48^, and curated drug-disease and gene-disease associations^36,37,49,50^ (**Figure S2**). In total, the AD signature library comprised 317 standardised entries spanning single-cell and single-nucleus omics (215 signatures), bulk multi-omics (65 signatures), and functional and genetic knowledge sources (37 signatures) (**Figure S8A; Supplementary Table 3**), capturing diverse profiling technologies, cell types, disease states, regulatory programmes and AD-relevant biological contexts (see **Supplementary Discussion**).

To represent these 317 signatures within a common biological scaffold, we mapped them onto the human protein-protein interaction (PPI) network obtained from STRING^51^ (confidence threshold ≥ 400; 19,486 proteins). For each network protein, we calculated three proximity features to every signature: the minimum shortest-path distance (MSP), the average shortest-path distance (ASP), and the proportion of immediate neighbours (ratio of neighbours, RN) belonging to the signature, resulting in 951 graph-derived features per protein (317 signatures x 3 proximity metrics; **Figure 1C**). Because PPI networks are biased toward well-studied, highly connected proteins^52^, we normalised each observed proximity value against a null distribution generated from size– and degree-matched random protein sets (**Figure 1D**). This yielded z-normalised features with substantially reduced dependence on degree centrality (**Figure 1E**). Principal component analysis of the resulting gene-level feature matrix showed that the three feature blocks captured partially overlapping but distinct sources of network structure (**Figure S8B & Extended Data Figure 2**). Finally, UMAP visualisation revealed a non-random distribution of Class 1 AD effectors within the derived representation, supporting the ability of the graph-derived feature space to capture AD-relevant biological signal (**Figure 1F**).

### BRIDGE-AD yields a genome-wide resource of prioritised AD effectors and outperforms published gene representations

Having defined the Class 1 positive labels and constructed the graph-derived feature space, we trained supervised machine learning (ML) models to prioritise AD effectors across all genes represented in the interactome (**Figure 2A**). The 153 Class 1 genes and the remaining 19,333 background genes (Class 0 genes) were split into training and test sets using an 80:20 split. To address class imbalance, training Class 0 genes were partitioned into balanced subsets matched to the number of Class 1 genes, generating 126 balanced datasets. On each balanced dataset, we trained gradient boosting (GB), logistic regression (LR), random forest (RF), and support vector machine (SVM) classifiers, along with a soft voting ensemble of the four. Using this framework, we first assessed the effect of correlation-based feature pruning on cross-validation performance by training classifiers on the full 951-feature matrix and on pruned matrices generated across a range of absolute Spearman correlation thresholds (**Figure S9A**). At each threshold, feature pairs with correlations above the cut-off were grouped into redundancy clusters, with one representative feature retained from each cluster. Feature pruning did not improve performance relative to the unfiltered baseline, with more aggressive filtering making the models more conservative by increasing precision at the expense of recall. This pattern suggests that the framework benefits from a broad set of complementary features, rather than relying on a small subset of dominant predictors. We therefore adopted the most conservative threshold (|ρ| ≥ 0.95), which reduced feature redundancy while preserving baseline performance. Across all feature matrices, the ensemble classifier consistently achieved the strongest performance and was selected for downstream modelling using the 893-feature matrix generated at the |ρ| ≥ 0.95 pruning threshold (**Table 1**).

**Figure 2.**
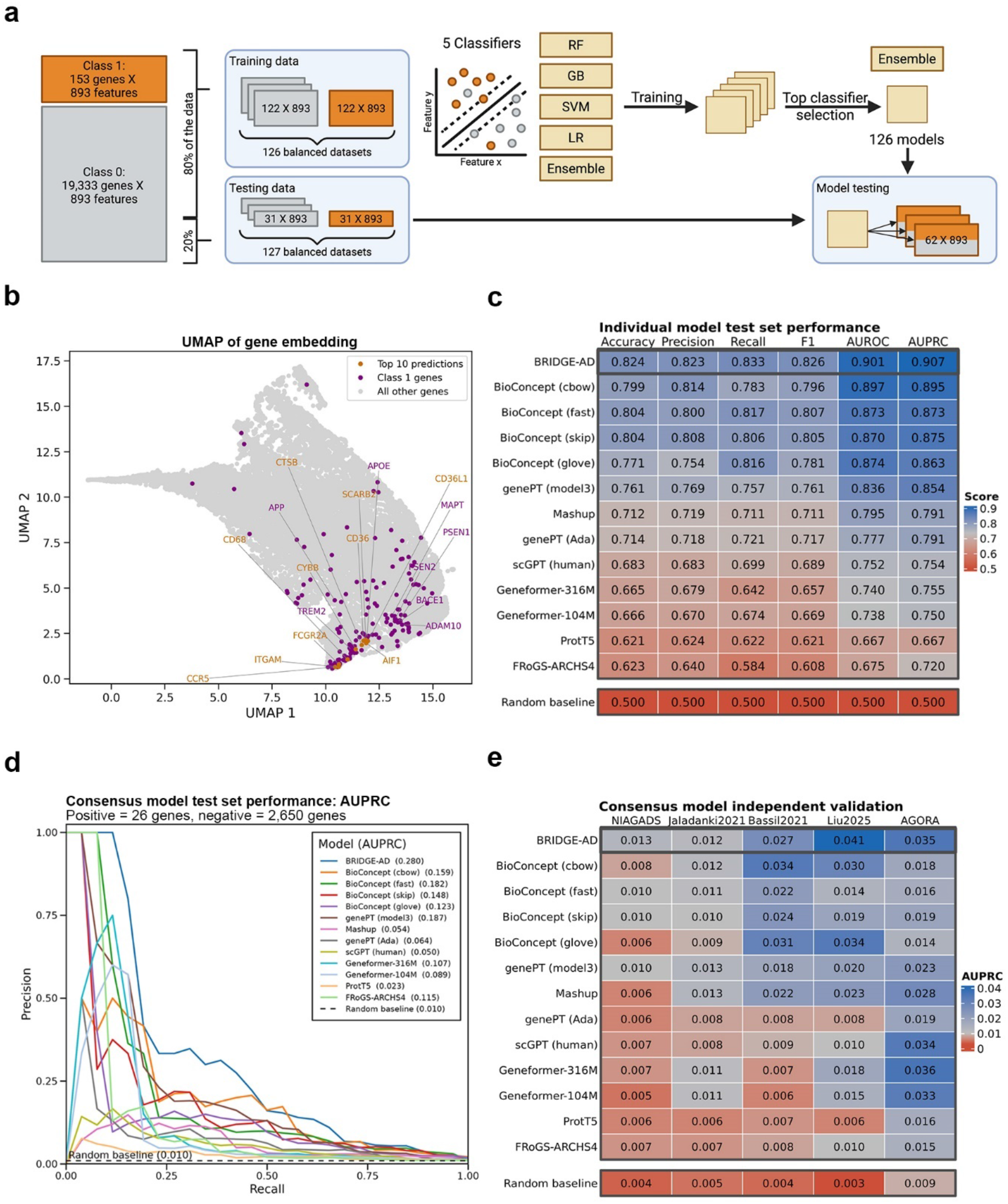
Implementation, performance, and benchmarking of the machine learning pipeline. **(A)** Schematic of the machine learning (ML) pipeline. A positive Class 1 set of 153 genes and the remaining 19,333 Class 0 genes were each represented by 893 network-derived features after correlation filtering at |ρ| ≥ 0.95. Class 1 and Class 0 genes were split into training (80%) and held-out test (20%) sets. To ensure balanced representation, Class 0 genes were randomly sampled without replacement to match the number of Class 1 genes. Four ML classifiers and an ensemble model were trained on each balanced dataset and evaluated by 5-fold cross-validation, after which the top-performing classifier was selected and assessed on the held-out test set. **(B)** UMAP embedding showing top 10 prioritised Class 0 genes in the context of Class 1 effectors. Labels highlight canonical AD effectors (purple) and top 10 predictions (yellow). **(C)** Test set performance of BRIDGE-AD and alternative gene embeddings on balanced test datasets. Each of the models trained on a balanced training dataset was evaluated across all balanced test datasets, with the heatmap summarising average test set accuracy, precision, recall, F1 score, AUROC and AUPRC. **(D)** Test set performance of BRIDGE-AD and alternative gene embeddings on the full imbalanced held-out test set. Consensus predictions were generated for BRIDGE-AD and each alternative embedding model by averaging the probability scores assigned to each held-out gene (Class 1 = 26 genes, Class 0 = 2,650 genes) across the trained models. Model performance was then assessed across the full test split using AUPRC. BRIDGE-AD achieved an AUPRC of 0.280, exceeding the random baseline AUPRC of 0.010 and outperforming alternative gene embeddings. **(E)** Independent validation of BRIDGE-AD and alternative gene embeddings using external AD effector gene sets. The heatmap shows AUPRC values measuring recovery of external AD-relevant positive label sets, with random baseline AUPRC shown for comparison.

**Table 1.**
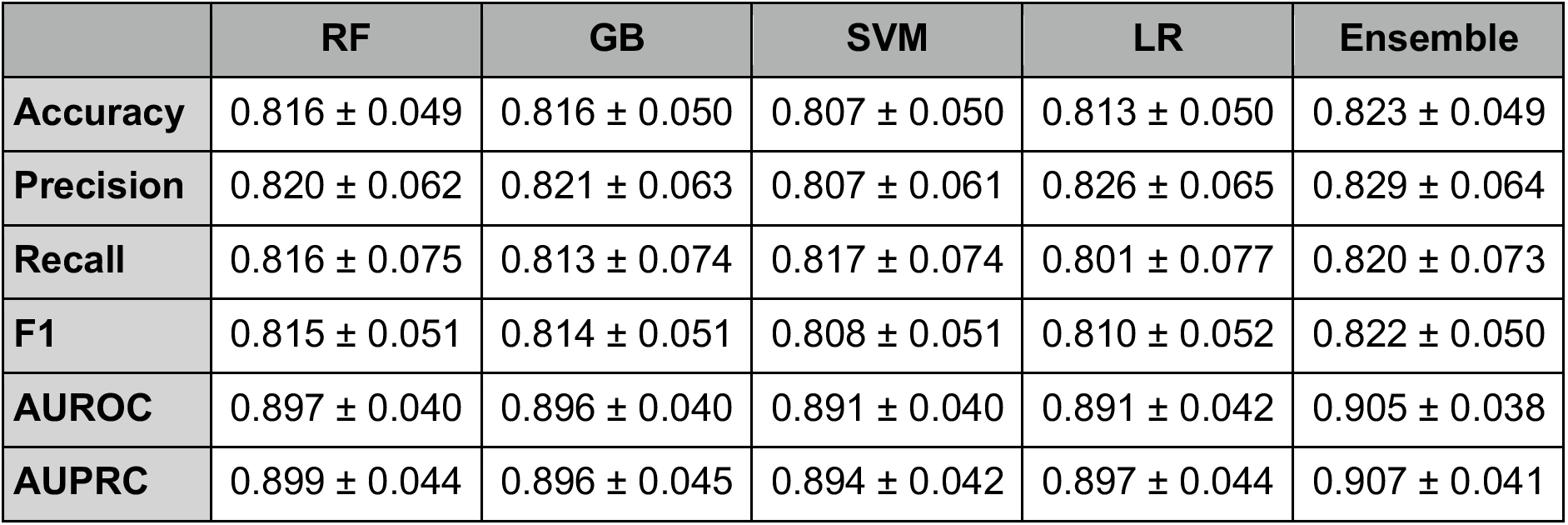
Five-fold cross-validation performance of the five classifiers on training data (|ρ| ≥ 0.95 feature pruning), reported as mean ± SD. AUROC, area under the receiver operating characteristic curve; AUPRC, area under the precision-recall curve.

We next evaluated the selected ensemble classifier on the held-out test split, comprising 31 Class 1 genes and 3,961 Class 0 genes. To account for class imbalance, we followed the training strategy as described above and generated balanced test datasets by sampling Class 0 genes to match the number of held-out Class 1 genes. Each of the 126 ensemble models was evaluated across the balanced test sets, achieving average accuracy of 0.851, precision of 0.840, recall of 0.872, F1 score of 0.855, AUROC of 0.909 and AUPRC of 0.915. Because the balanced sets do not reflect the marked scarcity of known disease effectors, we additionally assessed performance on the full imbalanced hold-out set. For this analysis, we generated consensus predictions by averaging the probability scores assigned to each held-out gene across the 126 ensemble models. The resulting consensus model achieved an AUPRC of 0.209 on the 3,992-gene hold-out set, substantially above the random baseline of 0.008 (**Figure S9B**). We subsequently examined model feature importance using Shapley Explanatory (SHAP) values^53^. We found that the top-ranked features included proximity to AD effectors from the Comparative Toxicogenomics Database, neuronal cell death suppressor hits from CRISPRi screens, GWAS candidates from the NHGRI-EBI GWAS Catalog, as well as snATAC-seq-derived regulatory signatures (**Figure S9D**). This feature profile suggests that BRIDGE-AD predictions are supported by complementary signals spanning curated disease knowledge, functional perturbation evidence, human genetic association, and cell type-resolved regulatory programmes. Finally, using the held-out test genes, we evaluated the precision–recall trade-off across Class 1 score thresholds to identify a high-confidence prioritisation cut-off that balanced the recovery of known AD effectors against the risk of nominating false-positive candidates (**Figure S9C**). Precision and recall curves intersected at a threshold of 0.889, which we used to define a high-confidence tier of prioritised AD effectors (see **Figure 3**). Together, these analyses yielded a genome-wide resource of prioritised AD effectors (**Supplementary Table 4**); UMAP visualisation of the gene-feature space further showed that known effectors and the top newly prioritised candidates clustered closely together (**Figure 2B**).

**Figure 3.**
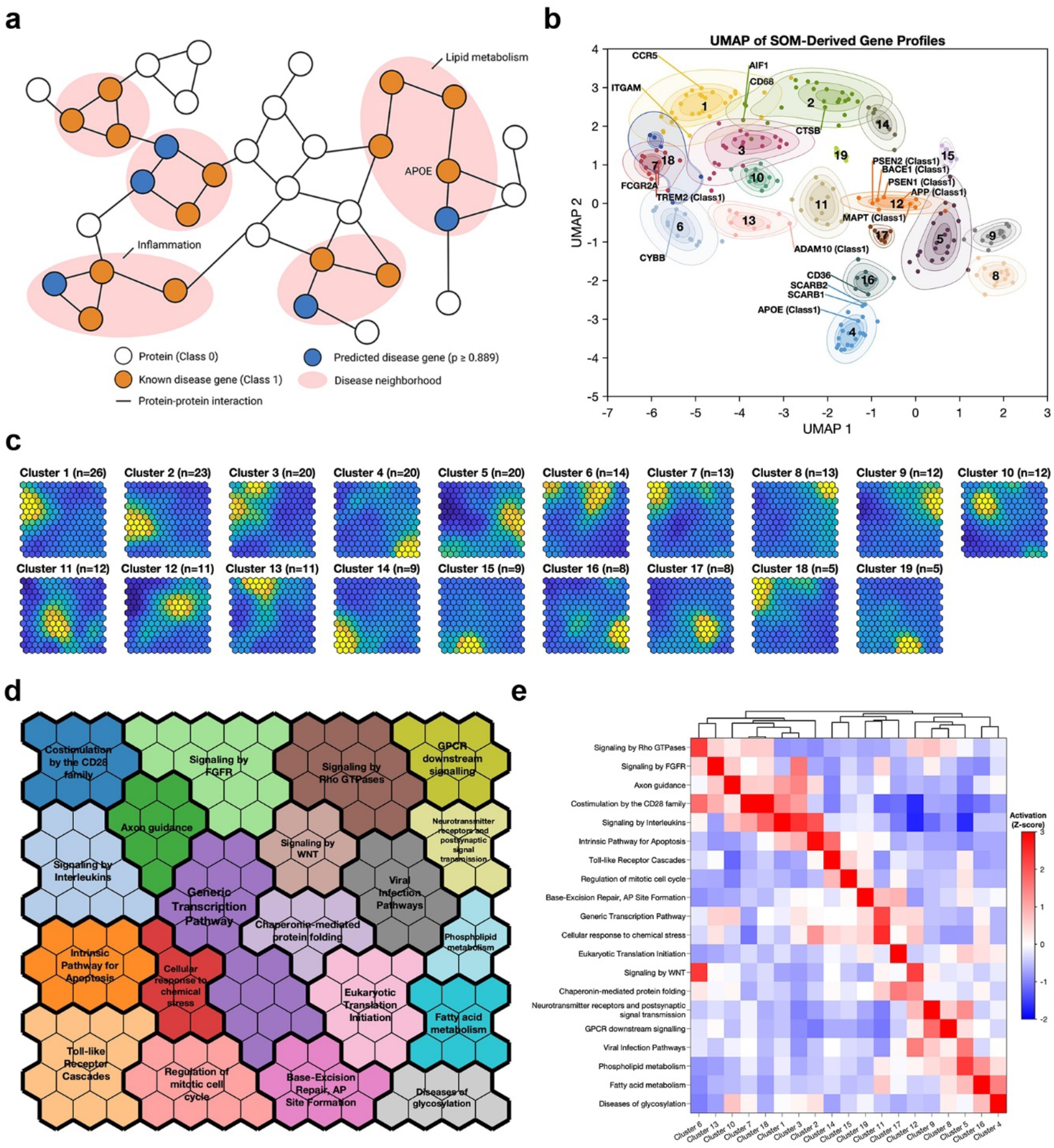
Functional clustering of the extended AD effector set by shared pathway activity. **(A)** Schematic of disease neighbourhoods in the interactome. Known disease genes (Class 1, orange) and predicted disease genes (blue) can colocalise within regions of the interactome enriched for functionally related genes, such as lipid metabolism and inflammation. **(B)** UMAP of self-organising map (SOM)-derived gene profiles, coloured by cluster assignment. Shaded contours indicate kernel density estimates per cluster (5^th^, 25^th^, 50^th^, and 75^th^ density quantiles). Labels highlight canonical AD effectors and top 10 predictions. **(C)** Average SOM activation profiles per gene cluster. Genes were grouped into 19 clusters based on the biological pathways that they share. Each heatmap represents one cluster, and each hexagon represents a distinct group of co-clustered pathways, occupying the same position across all heatmaps. The colour of each hexagon reflects how strongly the gene cluster is associated with the pathways at that location. **(D)** SOM grid partitioned into broad pathways. This map is the spatial key for the activation profiles shown in panel C: regions of high activation in each gene cluster correspond to the pathways named here. **(E)** Pathway activity across gene clusters. Each column represents a gene cluster and each row a broad pathway, with colour indicating how strongly each cluster is associated with that pathway.

To evaluate BRIDGE-AD in comparison with existing gene representations, we benchmarked it against twelve recently published gene embeddings derived from biomedical literature (BioConcept^1^, genePT^2^), PPI data (Mashup^3^), single-cell transcriptomics (scGPT^4^, Geneformer^5^), bulk transcriptomics (FRoGS-ARCHS4^6^), and protein sequence data (protT5^7^). Restricting the analysis to the 13,386 genes (Class 1 = 131, Class 0 = 13,255) shared across all embeddings, we used a common training and test split and evaluated all representations within the same classification framework. BRIDGE-AD consistently outperformed all competing embeddings, both under 5-fold cross-validation on the training data (**Figure S10**) and on the unseen test split (**Figure 2C**). As before, we also assessed performance on the full imbalanced hold-out set by averaging each gene’s probability scores into a consensus prediction and comparing AUPRC; BRIDGE-AD again ranked first, achieving the highest value (0.280; 2,676-gene hold-out set, baseline 0.010) of any approach (**Figure 2D**). To probe model behaviour further, we tested how well each embedding recovered five independent AD effector gene sets derived from AGORA^9^, NIAGADS^10^, Jaladanki et al. (2021)^54^, Bassil et al. (2021)^55^, and Liu et al. (2025)^56^. For each embedding, models were trained across all 13,386 shared genes using the original Class 1 labels as the positive class. To ensure unbiased evaluation, each Class 0 gene was scored by averaging predictions only from the balanced models in which that gene had not been included during training. Recovery of each independent gene set was quantified using AUPRC, with genes in the respective set treated as positives and all other Class 0 genes as negatives. BRIDGE-AD outperformed all twelve competing embeddings, indicating superior prioritisation of independently nominated AD genes (**Figure 2E**).

### Organisation of prioritised AD effectors into functional modules

The precision-recall threshold of 0.889 nominated 98 Class 0 genes as predicted Class 1 effectors (**Figure S9C, Supplementary Table 5**). Together with the 153 known Class 1 genes, these candidates defined an expanded set of 251 AD effectors. To examine the functional organisation of these genes, we applied self-organising map (SOM) analysis to the pathway enrichment profiles of their local PPI neighbourhoods (**Figure 3A**). We first extracted the neighbourhood of each effector and assessed Reactome^57^ pathway enrichment, retaining 877 pathways that were significantly enriched in at least three neighbourhoods. Enrichment precision and recall were then combined into an F1 score for each gene–pathway pair, producing a pathway-by-gene matrix. Training the SOM on this matrix arranged pathways with similar enrichment patterns into adjacent regions of the map and generated one component plane heatmap for each AD effector. In these heatmaps, each hexagon represents unique pathways that have clustered together, with hexagon positions conserved across all 251 maps. We then clustered the SOM-derived gene profiles, selecting the optimal number of clusters by maximising the mean silhouette width (**Figures 3B & S11A,** see **Methods**). This analysis yielded 19 gene clusters, for which we computed average SOM activation profiles (**Figure 3C & Supplementary Table 6**). Each gene cluster showed peak activation in a distinct map region, thereby defining its characteristic functional signature. To annotate these map regions, we separately clustered the SOM nodes (hexagons). Silhouette analysis identified 21 as the optimal number of SOM-node clusters (**Figure S11B**). The resulting clusters were annotated according to their dominant Reactome categories. Two clusters with the same dominant annotation were merged, yielding 20 pathway regions that provided the functional map shown in **Figure 3D**. Together, the SOM gene clusters captured multiple biologically coherent groups, including modules centred on lipid metabolism, cytokine and chemokine signalling, protein processing, and synaptic transmission, among others (**Figure 3E & S11C–F**). These results reveal a global molecular landscape of AD effector biology, in which known and newly prioritised candidates are organised into shared network-neighbourhood programmes.

### Investigating individual genes of interest with BRIDGE-AD Explorer

Whereas the SOM analysis captured broad molecular themes across the expanded AD effector set, we next sought to gain insights into individual prioritised genes. Because BRIDGE-AD is built from curated, dataset-derived signatures projected onto an interpretable network feature space, it is possible to trace back the underlying evidence layers to generate gene-level mechanistic hypotheses. To enable this, we developed the BRIDGE-AD Explorer, an interactive resource for visualising individual gene behaviour across the datasets analysed in this study (**Figure 4A–B**). The Explorer can be used to place top predictions across brain and peripheral cell types (**Figure 4C–E**) and enables formulation of complex cross-tissue hypotheses related to AD pathogenesis that could not be captured by any single omics dataset alone. We have made the Explorer accessible for public use at explore-bridgead.com.

**Figure 4.**
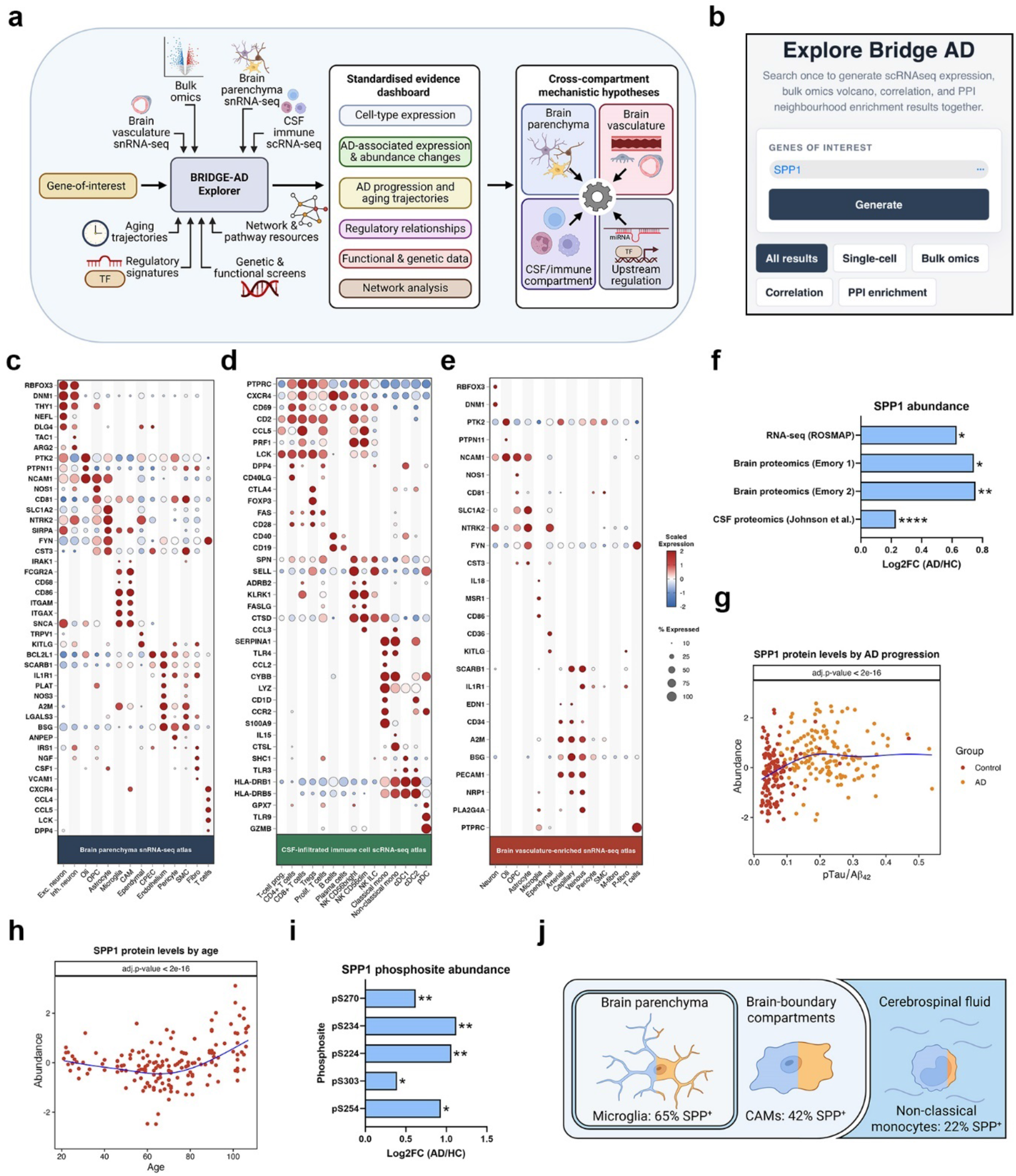
BRIDGE-AD Explorer enables gene-centric exploration across the integrated omics space and nominates SPP1 as a cross-compartment hypothesis candidate. **(A)** Schematic overview of the BRIDGE-AD Explorer. A gene of interest is queried across the integrated omics space to generate a standardised evidence dashboard summarising cell type expression, AD-associated expression and abundance changes, AD progression and aging trajectories, regulatory relationships, functional and genetic data, and network analysis results. These evidence layers can be integrated to generate cross-compartment mechanistic hypotheses spanning brain parenchyma, brain vasculature, CSF/immune compartments, and upstream regulatory mechanisms. **(B)** Snippet of the explore-bridgead.com interface, illustrating how a gene-of-interest query can be used to access BRIDGE-AD Explorer evidence modules, including single-cell, bulk omics, correlation and PPI enrichment results. **(C–E)** Expression of prioritised and cell type-enriched AD effectors across integrated single-cell and single-nucleus RNA-seq atlases, including brain parenchyma snRNA-seq **(C)**, CSF-infiltrated immune cell scRNA-seq **(D)**, and brain vasculature-enriched snRNA-seq **(E)**. Dot size indicates the percentage of cells expressing each gene, and colour indicates scaled mean expression. **(F)** AD-associated changes in *SPP1* RNA and protein abundance across bulk omics datasets, including ROSMAP RNA-seq, brain proteomics datasets, and CSF proteomics. **(G)** Disease-progression trajectory of SPP1 protein abundance in the CSF obtained from Johnson et al. **(H)** Age-associated trajectory of SPP1 protein abundance in plasma from healthy individuals obtained from Lehallier et al. **(I)** AD-associated changes in SPP1 phosphorylation across individual phosphosites. **(J)** Compartment– and cell type-specific *SPP1* expression across brain parenchyma, brain-boundary compartments, and cerebrospinal fluid, highlighting *SPP1* expression in microglia, CNS-associated macrophages (CAMs), and non-classical monocytes. (**F, I**) Asterisks indicate FDR-adjusted significance: *FDR < 0.05, **FDR < 0.01, and ****FDR < 0.0001. FDR, false discovery rate.

To illustrate how the diverse BRIDGE-AD omics space can be used to formulate mechanistic cross-tissue hypotheses, we first inspected the expanded AD effector set of 251 genes. We observed enrichment for an opsonophagocytosis-related programme that included integrin machinery (*ITGAX*, *ITGAM*, *ITGB2*), Fc receptors (FCGRs), TREM2–DAP12 signaling (*SYK*, *TYROBP*, *TREM2*, *INPP5D*, *PTK2B*), complement cascade components (*C1QA*, *C3*, *CLU*), and endolysosomal machinery involved in antigen processing and cargo degradation (*CD63*, *CD68*, *CTSS*, *CTSB*, *CTSD*) (**Figure S12A–B**).

To refine this broad programme into a coherent hypothesis, we searched for upstream extracellular effectors capable of engaging the opsonophagocytic machinery. We first selected genes belonging to the “Integrin cell surface interactions” Reactome pathway (R-HSA-216083) and then restricted this set to secreted factors (Human Protein Atlas Secretome) (**Figure S12C**). This analysis yielded 45 genes that we then ranked based on the BRIDGE-AD probability scores and focused on the top 10 genes: *FGB* (Class 1), *VCAM1*, *VWF*, *VTN*, *FN1*, *SPP1*, *FGA*, *FGG*, *CD44*, and *THBS1*. When queried in the BRIDGE-AD Explorer, *SPP1* showed the broadest cross-omics alterations among these candidates (**Figure S12D**). SPP1 analysis using the OmniPath database of molecular biology prior knowledge^58^ confirmed integrin receptors as direct *SPP1* targets (**Figure S12E**). Notably, prior studies have reported mixed effects of *SPP1* on AD pathogenesis, where *SPP1*-activated phagocytic programmes may lead to protective debris clearance or detrimental synaptic pruning in AD^59–61^. Extensive AD-associated changes in *SPP1* biology and its unresolved effects on AD progression motivated us to explore this effector further.

Inspection of bulk omics datasets revealed a consistent increase in *SPP1* RNA and protein levels in the brain and protein levels in the CSF of individuals with AD as compared to healthy controls (**Figure 4F–G**). *SPP1* protein levels also increased with age in the plasma of healthy individuals (**Figure 4H**). In addition to these abundance-level changes, multiple *SPP1* phosphosites showed increased phosphorylation in AD, suggesting that *SPP1* could exert disease-associated effects through altered phosphorylation-dependent functions (**Figures 4I and S12F**). BRIDGE-AD Explorer also revealed that *SPP1* was a target of miR-181c-5p, one of eight miRNAs curated from the Takousis et al. meta-analysis (see **Supplementary Methods**)^42^ and found to be downregulated in AD (**Figure S12G**). In miRDB and TargetScan databases, *SPP1* was annotated as containing a conserved, high-confidence miR-181c-5p binding site in its 3’ untranslated region (UTR). A literature search further identified a study by Wang et al.^62^ in which *SPP1* was validated as a miR-181c-5p target using a dual-luciferase reporter assay. Taken together, decreased miR-181c-5p levels and increased expression of its target *SPP1* provide a plausible mechanistic hypothesis for altered *SPP1* levels in AD.

We next asked which cell types contributed to *SPP1* expression in the human brain by querying *SPP1* across the analysed single-cell and single-nucleus atlases. In the brain parenchyma snRNA-seq dataset, *SPP1* expression was enriched in microglia and CNS-associated macrophages (CAMs) (**Figures 4J and S12H**), consistent with a recent study identifying perivascular macrophages (PVMs) as an early source of *SPP1* in AD^60^. In the CSF-infiltrated immune cell scRNA-seq dataset, *SPP1* expression was enriched in non-classical monocytes, which could represent an alternative brain-boundary source of *SPP1*. In contrast, *SPP1* expression was not detected in brain vascular cells.

Given that CSF-infiltrated immune cells could represent an additional source of *SPP1* in the brain, we next sought to understand how non-classical monocytes accumulated in the CSF and patrolled brain-boundary compartments. We searched for secreted factors that could attract monocytes and noted that annexin A1 (ANXA1), an immunomodulatory protein involved in leukocyte trafficking, was among the Class 1 AD effectors and specifically enriched in the ependymal cells lining the brain ventricles (**Figure S13A**). Analysis of bulk omics datasets revealed that ANXA1 brain protein levels were increased in individuals with AD as compared to healthy controls (**Figure S13B**). OmniPath analysis further identified formyl peptide receptors (FPR1/2/3) involved in leukocyte chemotaxis as direct ANXA1 targets (**Figure S13C**). Among these receptors, FPR1 was most frequently expressed in CSF-infiltrated non-classical monocytes, being detected in 81% of this population, while FPR3 was observed in 30% and FPR2 was rarely detected (2%) (**Figure S13D–E**). Finally, quantification of non-classical monocyte abundance across diagnoses revealed an increasing trend from healthy controls to individuals with AD (**Figure S13F**). These observations support a hypothesis in which increased *ANXA1* expression by ependymal cells promotes FPR1/3^+^ non-classical monocyte infiltration and patrolling of the CSF in AD through ANXA1-FPR1/3 signalling (**Figure S13G**).

Taken together, SPP1-centred analysis (**Figure S12**) and the ANXA1–FPR1/3 hypothesis (**Figure S13**) illustrate how the BRIDGE-AD Explorer can be used to assemble an integrated trail of evidence about prioritised AD effectors, including their functional roles, regulatory mechanisms, and cross-tissue interactions in AD.

### SCARB2 is a top prioritised effector associated with lysosomal and lipid-handling changes in microglia

To select a candidate for experimental follow-up, we sought to identify a highly ranked but comparatively under-studied effector in AD. First, we performed a PubMed search for co-mentions between each prioritised effector and “Alzheimer’s disease” or “dementia” keywords to gauge the novelty of the top candidates (**Figure 5A**). Scavenger receptor class B member 2 (SCARB2, also known as lysosomal integral membrane protein 2, LIMP2) emerged as one of the effectors with the highest predicted probabilities of belonging to Class 1, yet it was comparatively underrepresented in the AD literature. SCARB2 is a lysosomal membrane protein best known for its role in transporting β-glucocerebrosidase (GCase) from the endoplasmic reticulum (ER) to the lysosomal lumen (**Figure S14A**). SCARB2 contains a large luminal domain that binds GCase as well as a hydrophobic tunnel that can bind cholesterol and contribute to lysosomal cholesterol export^63^. GCase is a lysosomal enzyme involved in sphingolipid metabolism, whereas mutations in the *GBA1* gene encoding GCase or its mis-sorting due to SCARB2 deficiency can cause lysosomal disorders, particularly Gaucher disease^64^. In further support of its lipid handling functions, SCARB2 was assigned to the same SOM cluster as APOE and other lipid-processing effectors (**Figure S11C**). Together, these observations place SCARB2 at an intersection of lysosomal function and lipid metabolism, processes highly relevant to molecular AD progression.

**Figure 5.**
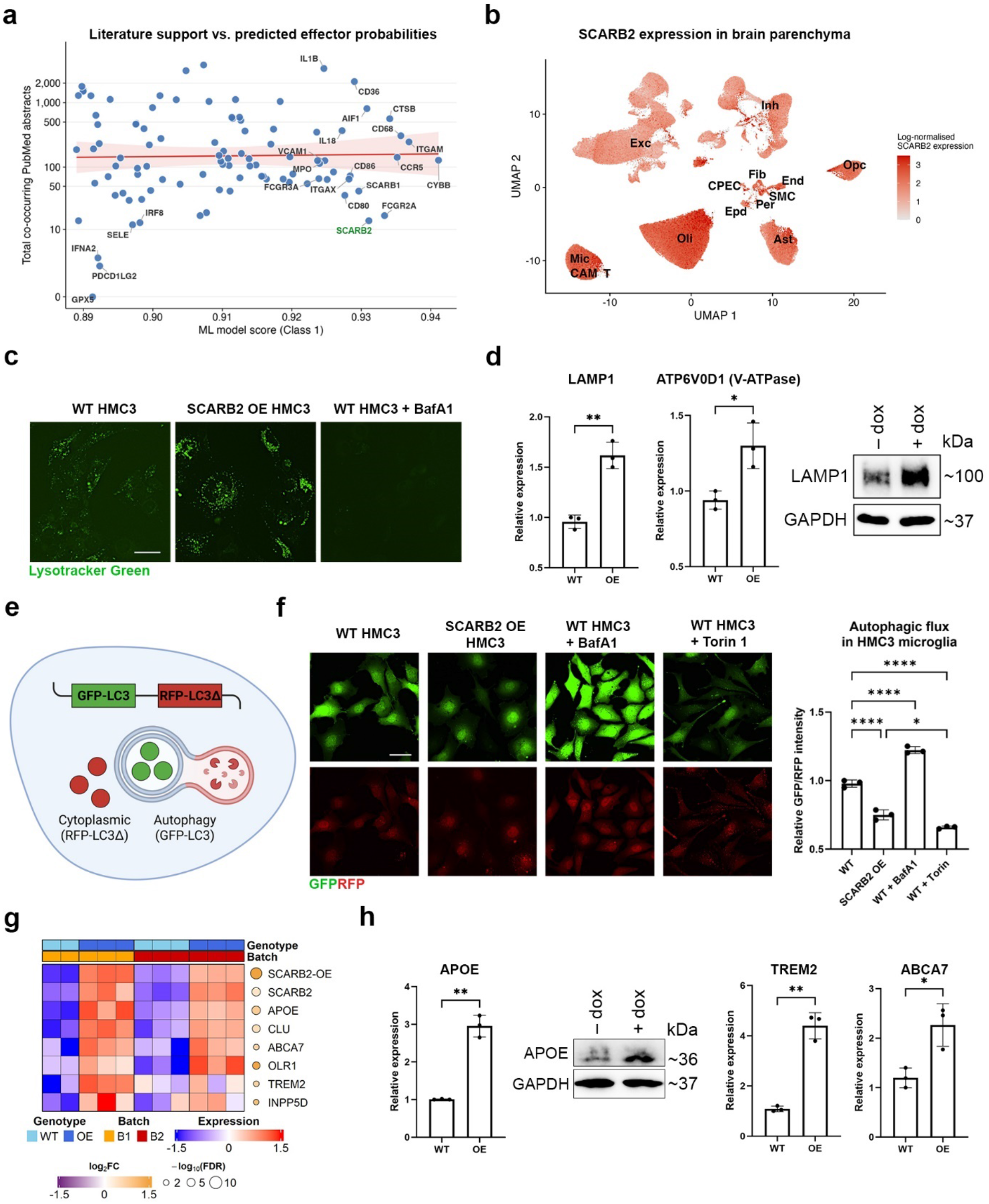
SCARB2 is a high-confidence Alzheimer’s disease effector candidate whose overexpression promotes lysosomal, autophagic, and lipid-processing programmes in HMC3 microglia. **(A)** Relationship between BRIDGE-AD effector probability and recent AD-related literature support. Each point represents a candidate effector plotted by predicted Class 1 probability and PubMed abstract co-mentions with “Alzheimer’s disease” or “dementia”. SCARB2 is highlighted as a high-confidence prediction with comparatively limited representation in AD literature. (B) *SCARB2* expression in the brain parenchyma snRNA-seq atlas from Mathys et al., shown as a UMAP feature plot. **(C)** Representative LysoTracker Green imaging of WT HMC3 microglia, *SCARB2*-overexpressing HMC3 microglia, and WT HMC3 treated with bafilomycin A1. *SCARB2* overexpression increases punctate acidic-compartment signal, consistent with expansion and/or increased acidification of the lysosomal compartment, whereas bafilomycin A1 serves as a lysosomal deacidification control. N = 3 independent biological replicates. Scale bar, 50 µm. **(D)** RT-qPCR analysis of *LAMP1* and *ATP6V0D1* expression, together with LAMP1 immunoblot validation, in WT and *SCARB2*-overexpressing HMC3 microglia. N = 3 independent biological replicates. **(E)** Schematic of the dual-colour GFP-LC3/RFP-LC3ΔG autophagic flux reporter. GFP-LC3 enters the autophagic pathway and is degraded, whereas RFP-LC3ΔG remains cytosolic as an internal control; therefore, a lower GFP/RFP ratio indicates increased autophagic flux. **(F)** Autophagic flux in WT and *SCARB2*-overexpressing HMC3 microglia. Representative GFP and RFP images and quantification of the GFP/RFP ratio are shown for WT, *SCARB2*-overexpressing, bafilomycin A1-treated, and Torin 1-treated conditions. Bafilomycin A1 and Torin 1 serve as negative and positive controls, respectively. Scale bar, 50 µm. **(G)** RNA-seq-based profiling of lipid-processing and AD-relevant genes altered by *SCARB2* overexpression in HMC3 microglia. N_WT_ = 5, N_OE_ = 6 independent biological replicates. **(H)** Orthogonal validation of the lipid-processing state induced by *SCARB2* overexpression. RT-qPCR and immunoblotting confirm increased *APOE* RNA and protein levels, and RT-qPCR shows increased expression of *TREM2* and *ABCA7* in *SCARB2*-overexpressing HMC3 microglia. N = 3 independent biological replicates. (**D, H**) RT-qPCR data are presented as mean ± SD. *p < 0.05, **p < 0.01 by a two-sided unpaired Welch’s t-test. (**F**) GFP/RFP ratios are presented as mean ± SD. *p < 0.05, ****p < 0.0001 by one-way ANOVA with Tukey’s multiple-comparisons test.

We queried SCARB2 with the BRIDGE-AD Explorer and found that SCARB2 protein levels were significantly increased in patients with AD as compared to healthy controls in the Morshed et al. dataset (log_2_FC = 0.2169, adj. p-value = 0.001257), but not in the other datasets examined (**Figure S14B**). Furthermore, SCARB2 was broadly expressed in the brain parenchyma and enriched in glial cells (**Figure 5B**). Pathway enrichment analysis highlighted lipid metabolism, scavenger receptor biology, and intracellular trafficking among SCARB2-associated processes (**Figure S14C**). We prioritised microglia as the model for mechanistic experiments for several reasons. First, many of the top prioritised candidates and the local SCARB2 SOM neighbourhood converged on myeloid and lipid-handling biology. Second, as professional phagocytes, microglia heavily rely on lysosomal programmes to process AD-relevant debris, including amyloid-β species. Third, Gaucher disease provides a myeloid-lineage precedent because its hallmark pathology arises from glucosylceramide-laden macrophages that experience a large burden of lysosomal lipids^64^. In microglia subtypes reported by Sun et al.^65^, mean SCARB2 expression was the highest in the MG4 cluster, annotated as lipid-processing microglia enriched for cholesterol and lipid homeostasis genes (**Figure S14D**).

Given that SCARB2 deficiency had already been associated with detrimental lysosomal phenotypes^66,67^, we sought to explore whether increasing SCARB2 levels was sufficient to enhance microglial lysosome– and lipid-processing-associated programmes, which could potentially be therapeutic in AD. Therefore, we overexpressed *SCARB2* in the HMC3 microglia cell line under a doxycycline-inducible promoter for downstream experiments (**Figure S14E**). We confirmed *SCARB2* overexpression (*SCARB2* OE) at the RNA and protein levels (**Figure S14F**) and found that SCARB2 colocalised with LAMP1^+^ lysosomes in HMC3 microglia (**Figure S14G**).

We first visualised the lysosomal compartment in wild-type (WT) and SCARB2 OE HMC3 using a pH-sensitive LysoTracker Green dye (**Figure 5C**). SCARB2 OE increased punctate LysoTracker Green signal, suggesting expansion and/or increased acidification of the lysosomal compartment upon SCARB2 OE. This phenotype was accompanied by increased *LAMP1* expression and protein levels, as well as increased expression of *ATP6V0D1*, a component of the lysosomal V-ATPase involved in lysosomal acidification, revealing structural and functional remodeling of the lysosomal compartment (**Figure 5D**). Our *in vitro* findings were supported by primary brain proteomics data, which revealed that *SCARB2* protein levels significantly correlated with those of various lysosomal components in the brain parenchyma (**Figure S15A– C**). Notably, the positive correlation between SCARB2 and LAMP1, LAMP2, and prosaposin (PSAP) was stronger in individuals with AD as compared to healthy controls, suggesting disease-enhanced coupling. Consistently, RNA-seq profiling of WT and *SCARB2* OE microglia revealed increased expression of various lysosomal genes after SCARB2 OE (**Figure S15E & Supplementary Table 7**). To test whether these lysosomal changes would influence autophagolysosomal processing, we measured autophagic flux using a dual-colour LC3 reporter (**Figure 5E**). In this system, GFP-LC3 serves as the autophagy-sensitive signal and RFP-LC3Δ serves as a non-lipidated cytoplasmic control; therefore, the GFP/RFP ratio decreases as autophagic flux increases. As expected, the autophagy inhibitor bafilomycin A1 increased the GFP/RFP ratio, whereas the autophagy activator Torin 1 decreased it (**Figure 5F**). Relative to WT microglia, *SCARB2* OE microglia exhibited a decreased GFP/RFP ratio, indicating that *SCARB2* OE is sufficient to increase autophagic flux in this microglia model.

We also found that various effectors related to lipid sensing and metabolism, including *APOE*, *CLU*, *TREM2*, *ABCA7*, *OLR1*, and *INPP5D*, were significantly upregulated after *SCARB2* OE, suggesting a coordinated remodelling of lipid handling programmes (**Figure 5G**). We confirmed increased *APOE* expression at the RNA and protein levels and increased *TREM2* and *ABCA7* expression at the RNA level in *SCARB2* OE HMC3 microglia by RT-qPCR and immunoblotting (**Figure 5H**). SCARB2 and APOE protein abundance also exhibited a significant positive correlation in the brain parenchyma (**Figure S15D**). We also noted increased *INPP5D* expression in *SCARB2* OE HMC3 microglia that, together with increased autophagic flux, suggested a possibility of reduced inflammasome tone^68^. We stimulated HMC3 cells with LPS and IFNγ, which robustly induced inflammatory gene expression (**Figure S15F**). In *SCARB2* OE HMC3 microglia, we detected reduced expression of *NLRP3* upon inflammatory stimulation as compared to WT HMC3, without a significant change in pro-inflammatory cytokine expression (**Figure S15G–H**).

Taken together, these results indicate that SCARB2 positively modulates lysosomal, autophagic and lipid-processing programmes in a microglial cell model. In the context of its computational nomination and its placement within a myeloid lysosome–lipid processing network neighbourhood, SCARB2 emerges as a candidate effector that intersects core molecular themes of AD, including lysosomal function, autophagy, lipid handling, and myeloid biology.

### Molecular consequences of dysregulated SCARB2 glycosylation in AD

To understand how SCARB2 is affected in AD, we first quantified SCARB2 and GCase protein levels in primary human brain tissues by immunoblotting **(Figure 6A & S16A– B)**. Mean SCARB2 and GCase abundance did not differ significantly between AD (19 subjects) and healthy control (16 subjects) brain samples, but SCARB2 levels were more heterogeneous in AD, with 2.67-fold higher variance by the Fligner-Killeen test (p = 0.037). We also noted that the SOM cluster 4 that contained SCARB2 was enriched for a Reactome term *Diseases of glycosylation,* as reflected by the SCARB2 SOM grid activation profile (**Figure S16C**). Furthermore, querying the BRIDGE-AD Explorer revealed dysregulated SCARB2 glycosylation in AD across multiple glycosites as reported by Zhang et al., whereas SCARB2 glycoforms were found to participate in AD-associated co-regulation modules^13,14^ (**Figure 6B**).

**Figure 6.**
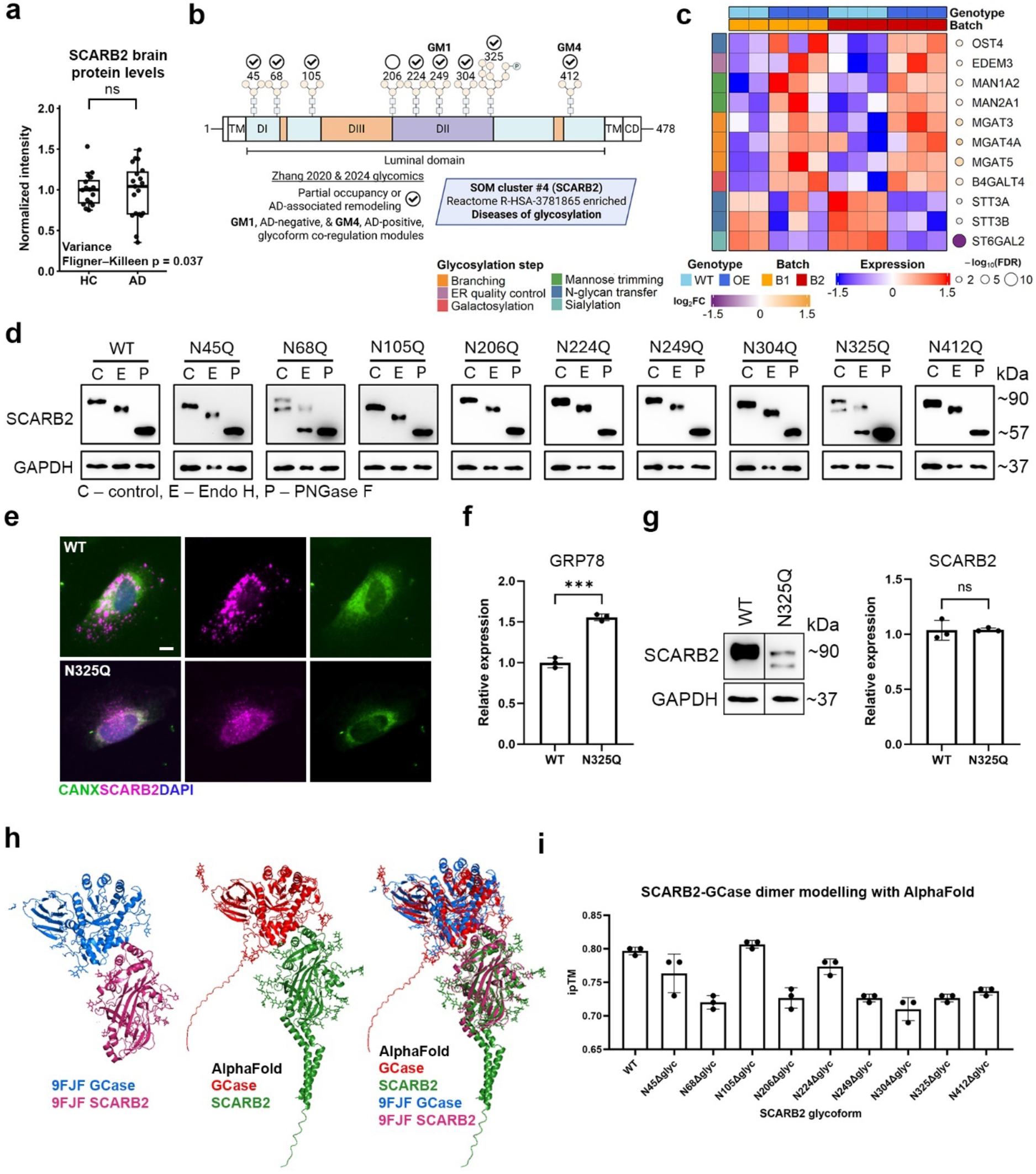
SCARB2 glycosylation is dysregulated in AD and influences SCARB2 maturation and fate. **(A)** Tukey box-and-whisker plot showing HC-median-normalised SCARB2 protein abundance in healthy control and AD brain samples. Boxes indicate the interquartile range, centre lines indicate medians, whiskers extend to values within 1.5×IQR, and points represent individual donors. Mean protein abundance was compared by Welch’s t-test and was not significantly different between diagnostic groups, whereas SCARB2 variance was increased in AD by the Fligner-Killeen test. N_HC_ = 16, N_AD_ = 19 brain tissue donors. **(B)** Schematic and glycoproteomic summary of SCARB2 N-glycosylation. Validated luminal N-glycosylation sites are shown across the SCARB2 protein, together with: partial occupancy or AD-associated glycosylation changes detected in Zhang et al. (2020) and Zhang et al. (2024) brain glycoproteomics datasets; assignment to glycoform co-regulation modules; and the relationship of SCARB2 to SOM cluster 4, which was enriched for the Reactome term *Diseases of glycosylation*. **(C)** Expression of glycosylation-related genes in WT and *SCARB2*-overexpressing HMC3 microglia, grouped by glycosylation step or pathway category. N_WT_ = 5, N_OE_ = 6 independent biological replicates. **(D)** Biochemical analysis of WT SCARB2 and single-site SCARB2 N→Q glycoforms using control buffer, Endo H, and PNGase F digestion followed by immunoblotting; GAPDH is shown as a loading control. **(E)** Representative immunofluorescence images of WT and *SCARB2-N325Q*-overexpressing HMC3 microglia immunolabelled for calnexin (CANX) and SCARB2 and counterstained for DAPI. Scale bar, 10 µm. **(F)** *GRP78* expression in WT and *SCARB2-N325Q*-overexpressing HMC3 microglia. N = 3 independent biological replicates. **(G)** *SCARB2* RNA levels and protein abundance in WT– and *SCARB2-N325Q*-overexpressing HMC3 microglia, showing reduced SCARB2-N325Q protein abundance despite comparable transcript levels. N = 3 independent biological replicates. **(H)** Structural modelling of the SCARB2-GCase complex. The *AlphaFold*-modelled full-length SCARB2-GCase complex is compared with an experimentally resolved SCARB2-GCase luminal-domain complex (PDB ID: 9FJF^74^). **(I)** Interface predicted template modelling (ipTM) scores for SCARB2-GCase complexes after removal of individual SCARB2 glycans, suggesting that disruption or remodelling of specific N-glycans may alter predicted SCARB2-GCase complex stability. N = 3 random seeds. (**F, G**) RT-qPCR data are presented as mean ± SD. ***p < 0.001 by a two-sided unpaired Welch’s t-test; ns, not significant. (**I**) Data are presented as mean ± SD.

Taken together, the increased variability of SCARB2 protein abundance in AD brain samples and the AD-associated remodelling of SCARB2 glycosylation suggested that post-translational regulation rather than uniform abundance change may influence SCARB2 maturation, trafficking, and functional availability in AD. Therefore, we next sought to characterise SCARB2 N-glycosylation.

First, we found that *SCARB2* overexpression altered expression of glycosylation-related genes in HMC3 microglia, including genes involved in branching, ER quality control, galactosylation, mannose trimming, N-glycan transfer, and sialylation (**Figure 6C**). Although it is challenging to model N-glycosylation gain using cellular models, we aimed to understand how a loss of individual SCARB2 glycans affects its fate. To this end, we generated WT SCARB2 and nine single-site N→Q mutant glycoforms, each lacking one validated N-glycosylation site, by replacing asparagine with a structurally similar glutamine residue at each position **(Figure S16D)**. We then used Endo H and PNGase F glycosidases to characterise SCARB2 glycoform maturation. N-glycosylation is initiated in the ER, where the oligosaccharyltransferase (OST) complex transfers high-mannose glycans onto the native peptide^69^. As the protein progresses through the Golgi complex, some high-mannose glycans are remodelled to generate mature hybrid and complex glycans (**Figure S16E**). High-mannose and hybrid glycans are sensitive to Endo H digestion, whereas complex glycans are Endo H-resistant; in contrast, PNGase F removes all glycans. Thus, glycosidase-dependent mobility shifts can be used to distinguish immature ER-associated glycoforms from more mature forms that have progressed through the Golgi apparatus. To characterise SCARB2 glycoforms, we overexpressed each glycoform in HMC3 microglia, performed digestions with Endo H and PNGase F, and visualised banding patterns with SDS-PAGE and immunoblotting (**Figure 6D**). WT SCARB2 exhibited the expected digestion pattern: a modest downward shift after Endo H digestion and a stronger downward shift after PNGase digestion, consistent with a mixed mature glycan population containing both Endo H-sensitive and Endo H-resistant glycan species. Several mutant glycoforms, such as N45Q and N105Q, migrated faster than the WT glycoform in the untreated control condition, confirming the loss of a single glycan. Notably, N68Q and N325Q glycoforms each exhibited two bands in the untreated control lane, suggesting the presence of a mature form and an immature form potentially retained in the ER. After Endo H digestion, the lower band collapsed to the PNGase F level, indicating that all glycans of this immature form remained sensitive to Endo H, and that N68Q and N325Q SCARB2 glycoforms were partially retained in the ER. We confirmed partial ER retention of the SCARB2-N325Q glycoform by immunostaining for SCARB2 and an ER marker calnexin (**Figure 6E**). We observed markedly increased colocalisation of SCARB2 and calnexin in *SCARB2-N325Q* OE HMC3 microglia as compared to WT SCARB2 OE HMC3 microglia. ER retention of the SCARB2-N325Q glycoform was accompanied by significantly increased *GRP78* (encoding an ER chaperone BiP) expression and an increasing trend in *GRP94* expression, indicating compensatory chaperone engagement in response to SCARB2 misprocessing (**Figures 6F & S16F**). At the same time, the SCARB2-N325Q glycoform underwent active degradation, as indicated by sharply reduced SCARB2-N325Q protein levels as compared to those of WT SCARB2 despite comparable expression at the RNA level (**Figure 6G**). To determine whether this phenotype reflected broad ER stress, we examined unfolded protein response (UPR) markers *ATF6* and *CHOP* (**Figure S16G**). We found that expression of *ATF6* and *CHOP* was unchanged in *SCARB2-N325Q* OE HMC3 microglia as compared to WT controls, suggesting a limited proteostatic response rather than global ER stress.

Next, we asked whether the changing N-glycosylation profile of SCARB2 could influence the stability of SCARB2-GCase complexes that are required for GCase delivery to the lysosome. We modelled the SCARB2-GCase dimer using *AlphaFold* and found that it showed near-superimposable geometry with an experimentally resolved luminal domain (**Figure 6H**). We subsequently modelled the removal of individual SCARB2 glycans (see **Methods**) and compared interface predicted template modelling (ipTM) scores for the resulting complexes (**Figure 6I**). We found that removal of various glycans reduced the ipTM score, suggesting that disruption or remodelling of specific N-glycans may weaken or destabilise SCARB2-GCase coupling. Finally, we extended the glycosylation analysis from SCARB2 to GCase, which contains four N-glycosylation sites, three of which showed AD-associated remodelling and were assigned to AD-positive GFM4 and GFM1 glycoform co-regulation modules by Zhang et al.^13,14^ (**Figure S16H**). In glycosidase digestion-based assays, WT and mutant GCase glycoforms were Endo H-sensitive, indicating that under these overexpression conditions the GCase protein remains in the ER (**Figure S16H**). One plausible explanation is that endogenous SCARB2 becomes saturated and cannot efficiently shuttle overexpressed GCase to generate a separate mature band on an immunoblot. Modelling SCARB2-GCase dimers with mutant GCase glycoforms revealed a decreased predicted modelling score of the GCase-N58Q glycoform but not other mutant glycoforms (**Figure S16I**). Taken together, these data support a model in which glycosylation defects can perturb both sides of the SCARB2-GCase axis, affecting protein maturation, trafficking, and cellular states.

## Discussion

Integrating and effectively leveraging heterogeneous disease evidence remains a central challenge in contemporary disease biology. This challenge is especially pronounced in AD, where disease progression unfolds across molecular, cellular, tissue and temporal scales and involves multiple cell and tissue compartments^70–72^. To address this challenge, we developed BRIDGE-AD as an interpretable network medicine framework that generates a unified, gene-level representation of AD molecular biology. BRIDGE-AD integrates more than 30 datasets and curated resources by transforming diverse outputs into harmonised disease signatures, projecting these signatures onto a common protein interactome, and learning the network proximity patterns that distinguish experimentally supported AD effectors from the rest of the proteome. Through this approach, BRIDGE-AD provides both a genome-wide prioritisation resource and a structured representation of AD-associated network biology.

A conceptual strength of BRIDGE-AD is that it represents heterogeneous evidence within a shared coordinate system while preserving the biological specificity of the underlying datasets. We achieved this by converting each analysis output into a distinct gene signature, including separate signatures for individual cell types, disease states, age-associated trajectories, regulatory programmes, and functional or genetic evidence. This structure allows each prioritised gene to be traced back to the particular signatures, disease contexts, and datasets that support its nomination. Such interpretability is essential for moving beyond ranked gene lists toward mechanistic hypotheses that can guide experimental follow-up.

Several aspects of the modelling strategy were designed to ensure robust gene prioritisation. Disease signatures were mapped onto the STRING PPI network^51^, and the resulting network proximity features were normalised against degree-matched random expectations, consistent with network medicine strategies for reducing biases associated with highly connected proteins^52^. BRIDGE-AD also incorporated three complementary network proximity measures – average shortest-path distance, minimum shortest-path distance, and ratio of neighbouring signature genes – capturing each gene’s relationship to disease-associated signatures at the global, nearest-member, and local-neighbourhood scales. Correlation-based pruning indicated that moderate feature reduction did not affect performance, whereas more aggressive pruning reduced recall, suggesting that BRIDGE-AD benefits from a broad and distributed feature space rather than relying on a small number of dominant predictors.

The benchmarking results showed that BRIDGE-AD captures AD-relevant gene relationships more effectively than recently published embeddings derived from the biomedical literature, transcriptomic data, protein sequences, and general network biology. BRIDGE-AD also demonstrated superior recovery of independent AD gene sets, indicating that it reflects disease biology not limited to the Class 1 labels used for training. This supports the central hypothesis that encoding multimodal AD evidence in a disease-specific gene representation improves disease effector discovery beyond what can be achieved with embeddings developed for broader, disease-agnostic applications.

Self-organising map analysis showed that the expanded AD effector set is organised into multiple functional neighbourhoods rather than converging on a single dominant disease axis: these span lipid metabolism, cytokine and chemokine signalling, lysosomal and protein-processing pathways, synaptic transmission, and extracellular matrix and adhesion programmes. Because newly prioritised candidates colocalise with well-characterised effectors, the map provides a functional context for genes whose disease relevance is not apparent from the literature alone.

The SPP1-centred analysis illustrates how the BRIDGE-AD evidence space can be used to progress from prioritisation to hypothesis generation. Filtering prioritised candidates for secreted integrin-binding proteins, followed by gene-centric review in the BRIDGE-AD Explorer, identified SPP1 as the extracellular effector with the broadest cross-omics dysregulation among the top candidates. Its increased abundance and phosphorylation in AD were supported by proteomic and phosphoproteomic evidence, whereas its regulation by miR-181c-5p was consistent with AD-associated decrease in miR-181c-5p levels. In addition to its expression in microglia and CAMs, *SPP1* was also expressed in CSF-infiltrated non-classical monocytes. Consistent with this cross-compartment model, the Class 1 effector ANXA1 and its receptors FPR1/FPR3 supported a candidate brain-boundary signalling axis that could link CSF-infiltrated non-classical monocytes to SPP1-associated activity in AD. Together these observations support a multifactorial model in which SPP1 may connect aging, brain-boundary immune activity, CSF-infiltrating monocytes, and integrin-linked phagocytic programmes.

Our experimental investigation of SCARB2 provides an example of how BRIDGE-AD can nominate a comparatively underexplored effector for mechanistic follow-up. SCARB2 was assigned one of the highest Class 1 probabilities despite limited representation in the AD literature, and its network context suggested a convergent programme of lysosomal function, lipid handling, and myeloid biology. In HMC3 microglia, SCARB2 overexpression increased punctate acidic-compartment signal, elevated expression of lysosomal components, enhanced autophagic flux and induced a coordinated lipid-processing programme involving APOE, TREM2, CLU, ABCA7, OLR1, and INPP5D and related effectors. These findings suggest that increased SCARB2 is sufficient to shift microglia toward a lysosome-competent, lipid-handling state.

A noteworthy aspect of SCARB2 biology in this study is that its disease relevance was not captured simply by mean changes in total abundance. Brain tissue analysis revealed unchanged average SCARB2 and GCase protein levels across diagnosis but substantial donor-to-donor variability, including significantly increased variance of SCARB2 protein levels in AD. Furthermore, glycoproteomic data revealed extensive site-selective remodelling of SCARB2 N-glycosylation in AD. Our mutagenesis and glycosidase digestion experiments indicated that loss of specific glycans, particularly at N68 and N325, can alter SCARB2 maturation. The N325Q glycoform showed partial endoplasmic reticulum retention, increased GRP78/GRP94 chaperone engagement without broad ATF6/CHOP induction, and reduced protein abundance despite comparable transcript levels. These results support a model in which AD-associated glycosylation changes may influence SCARB2 processing, trafficking and stability. Overall, BRIDGE-AD establishes an interpretable framework for integrating heterogeneous AD datasets, prioritising candidate effectors, and generating experimentally actionable hypotheses. The framework is useful not only because it produces high-confidence predictions, but also because the underlying evidence layers remain accessible for gene-centric exploration. In this study, that structure enabled progression from global AD effector prioritisation to pathway-level organisation, to SPP1-centred cross-tissue hypothesis generation, and finally to functional characterisation of SCARB2. More broadly, the same strategy should be adaptable to other complex diseases for which rich omics resources and curated perturbational knowledge are available.

Several limitations should be considered. First, BRIDGE-AD depends on the quality and scope of the curated Class 1 reference set. We used stringent criteria requiring experimental evidence that gene perturbation alters AD-relevant phenotypes, but the AD literature is shaped by historical research trends, unequal publication of positive and negative findings, and uneven evidence depth across genes. The Class 1 set is therefore a high-confidence training resource rather than an exhaustive catalogue of all causal AD effectors. Second, BRIDGE-AD prioritises genes as candidate AD effectors but does not infer whether increasing or decreasing each gene’s activity would be beneficial, because Class 1 labels combine protective and detrimental perturbation evidence to learn effector likelihood rather than therapeutic direction; nonetheless, we have successfully applied the BRIDGE-AD Explorer to infer directionality and design targeted experiments. Third, the current embedding is built on a protein interaction network and does not represent non-coding interactions that could potentially enrich the BRIDGE-AD model^73^. Fourth, the SPP1-centred and ANXA1–FPR hypotheses were generated by integrating cross-compartment evidence and will require experimental follow-up to test the predicted cellular interactions and signalling relationships. Fifth, our experimental work focused on SCARB2 as one high-priority candidate to demonstrate mechanistic follow-up, while other prioritised effectors and context-specific regulatory relationships remain to be tested. Taken together, BRIDGE-AD outputs should be interpreted as evidence-guided prioritisation for follow-up studies.

## Acknowledgements

The authors would like to thank Louise and Herbert Horvitz, the Christopher Family, the Judy and Bernard Briskin Fund, and the Sidell Kagan Foundation for their foresight and generosity; Banner Sun Health Research Institute for providing human brain tissues; and Dr. Qi Cui for providing support with immunoblotting and qPCR experimental design and interpretation. The authors would like to thank Drs. John Skidmore and Wei-Li Kuan for helpful suggestions. The authors thank Dr. Fred H. Gage for providing the pLVX-UbC-rtTA-Ngn2:2A:Ascl1 plasmid, Dr. Noboru Mizushima for providing the pMRX-IP-GFP-LC3-RFP-LC3ΔG plasmid, and Dr. Didier Trono for providing the psPAX2 and pMD2.G plasmids through Addgene. Figures 1C, 1D, 2A, 3A, 4A, 4J, 5E, 6B, S12B, S12C, S12E, S12F, S12G, S14E, S15F, S16D, S16E, S16H were created with BioRender.com. This work was supported by the National Institute on Aging of the National Institutes of Health R01 AG072291 and R01 AG079307 to Y.S.; the NIHR Cambridge Biomedical Research Centre (BRC-1215-20014) to K.S-P.; and the National Research Foundation of Korea grant funded by the Ministry of Science and ICT (RS-2025-18362970) and the Brain Pool Plus Fellowship Program funded by the Ministry of Science and ICT (RS-2025-25427881) to N.H. G.B. was funded by Standigm. J.C. was a predoctoral scholar in the Stem Cell Biology and Regenerative Medicine Research Training Program of the California Institute for Regenerative Medicine (CIRM) during a part of this study (EDUC4-12772). The funders had no role in study design, data collection and analysis, decision to publish, or preparation of the manuscript.

## Author Contributions

GB and JC contributed equally and share first authorship without precedence. GB, JC, KSP, YS, and NH conceived the project and designed the study. GB and JC jointly performed experiments and wrote the manuscript. GB led computational experiments reported in Figures 1–3. JC led laboratory experiments reported in Figures 4–6. HC developed an interactive website. Guoqiang Sun generated the GFP-LC3 reporter knock-in line. ZCEA performed APOE and LAMP1 immunoblotting. MR cloned FLAG-SCARB2 and GBA1-FLAG vectors. DW contributed to SCARB2 OE cloning and characterisation. Guihua Sun provided support for SCARB2 and GBA1 glycoform cloning design. TZ provided support with TREM2 characterisation. Guoqiang Sun, ZCEA, MR, and DW provided additional experimental support. DS provided senior support to HC. KSP, YS, and NH provided senior support and supervised the study.

**Figure S1.**
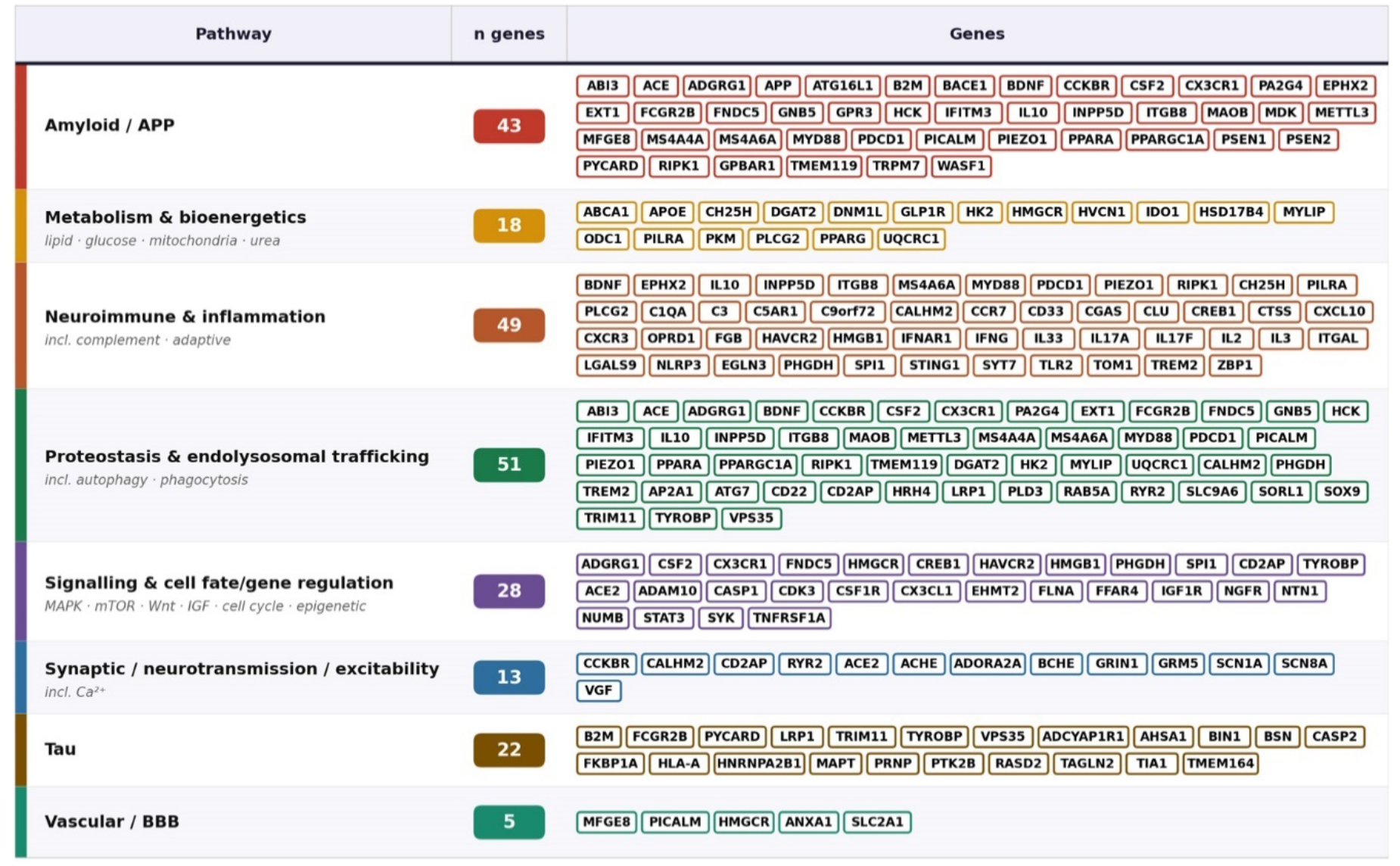
Pathway distribution of Class 1 effectors. Dominant pathway annotations were assigned as reported in the studies supporting each gene’s Class 1 designation.

**Figure S2.**
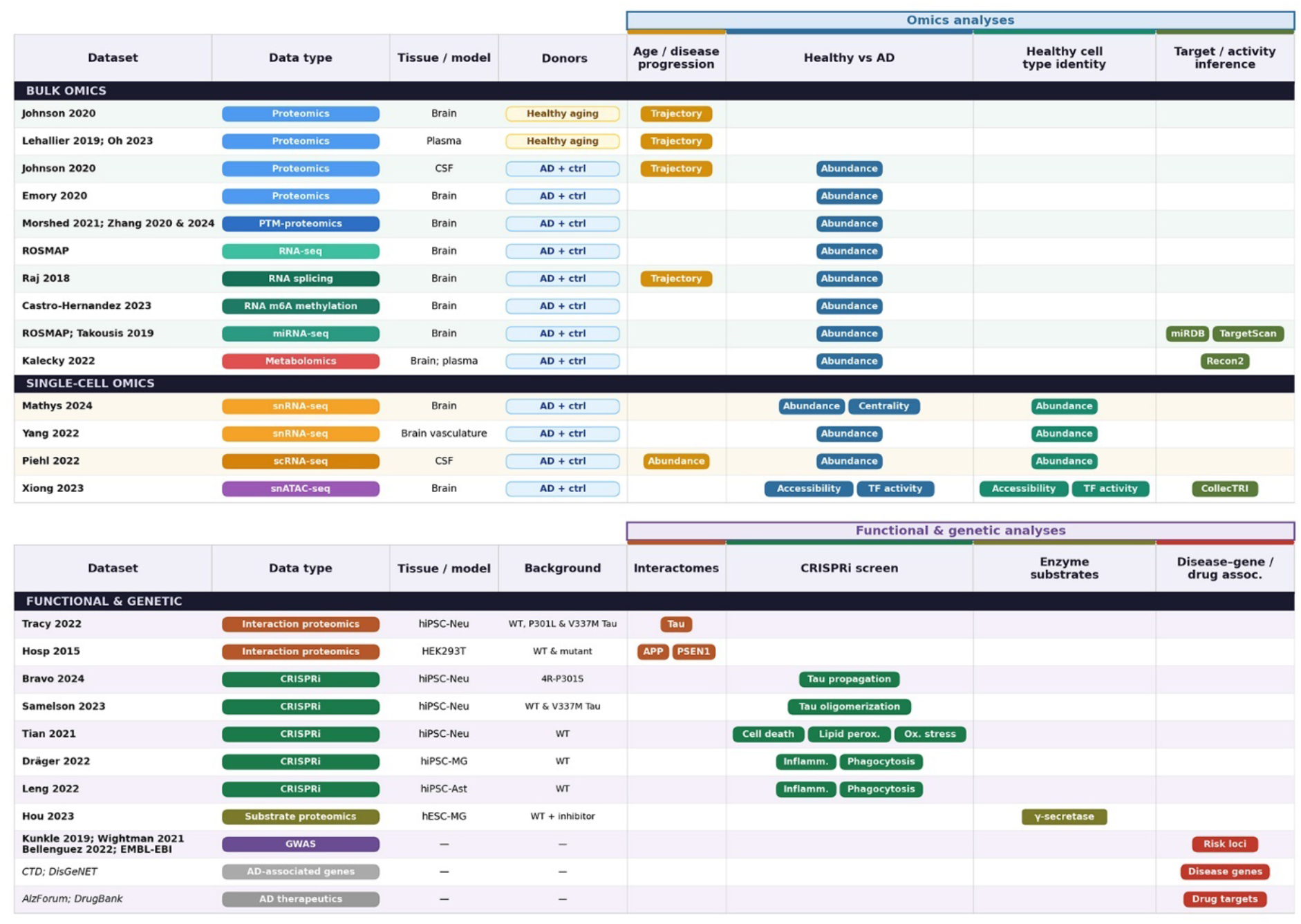
Datasets and resources used to construct the AD signature library. Overview of the bulk multi-omics datasets, single-cell and single-nucleus atlases, genetic studies, and functional knowledge resources analysed to generate AD gene signatures. For each resource, the analyses performed to derive the corresponding signatures are indicated.

**Figure S3.**
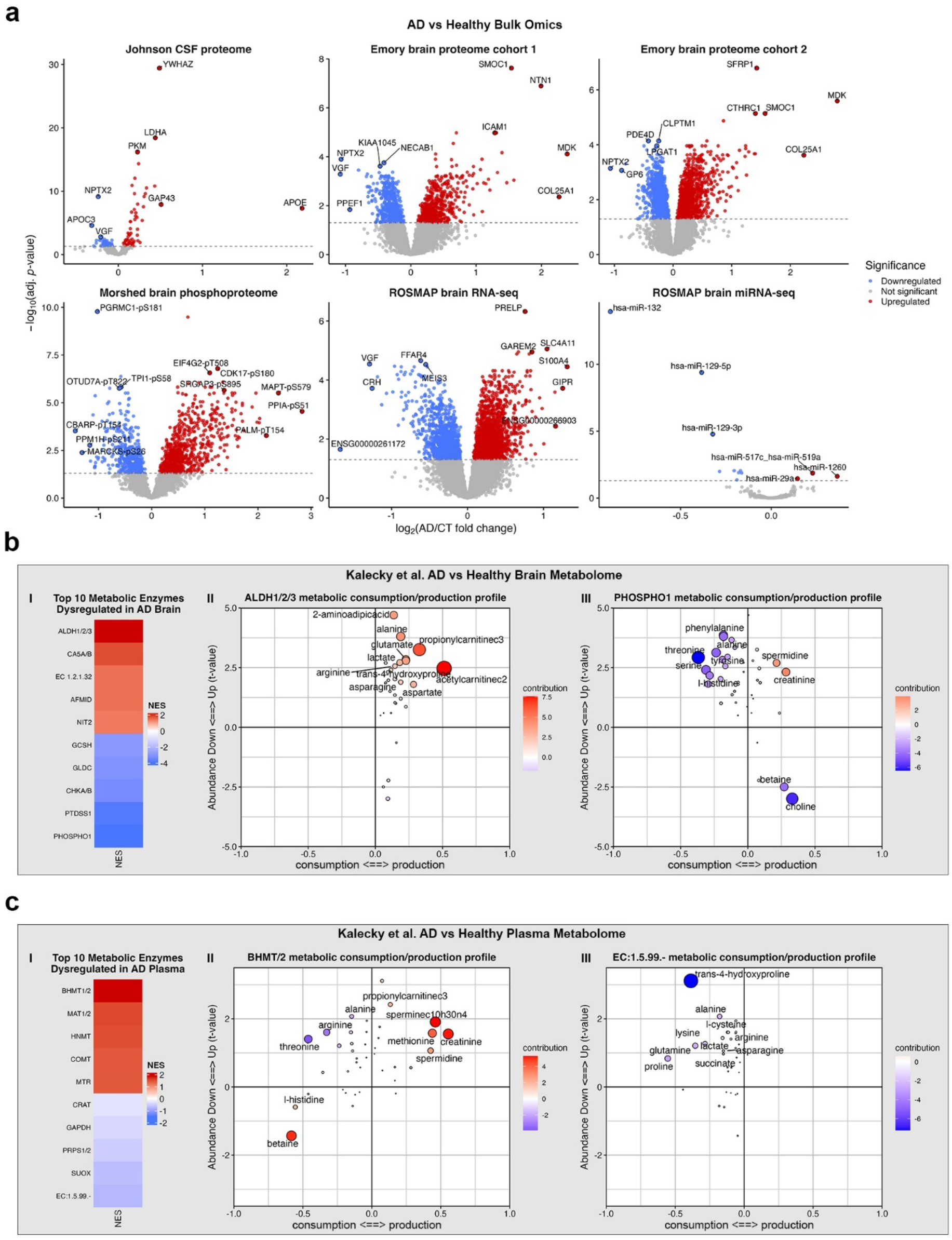
Bulk omics case-control analyses. **(A)** Volcano plots showing differentially expressed genes (DEGs) or differentially abundant proteins (DAPs) across bulk omics datasets. Features are coloured as significantly upregulated, significantly downregulated, or non-significant in AD donors relative to healthy controls (FDR < 0.05). **(B-C)** Top dysregulated enzymes inferred from **(B)** brain and **(C)** plasma metabolomics datasets. Panels I show enzyme-level normalised enrichment scores (NES), while panels II and III show example metabolite profiles used to infer enzyme activity changes (see **Methods**).

**Figure S4.**
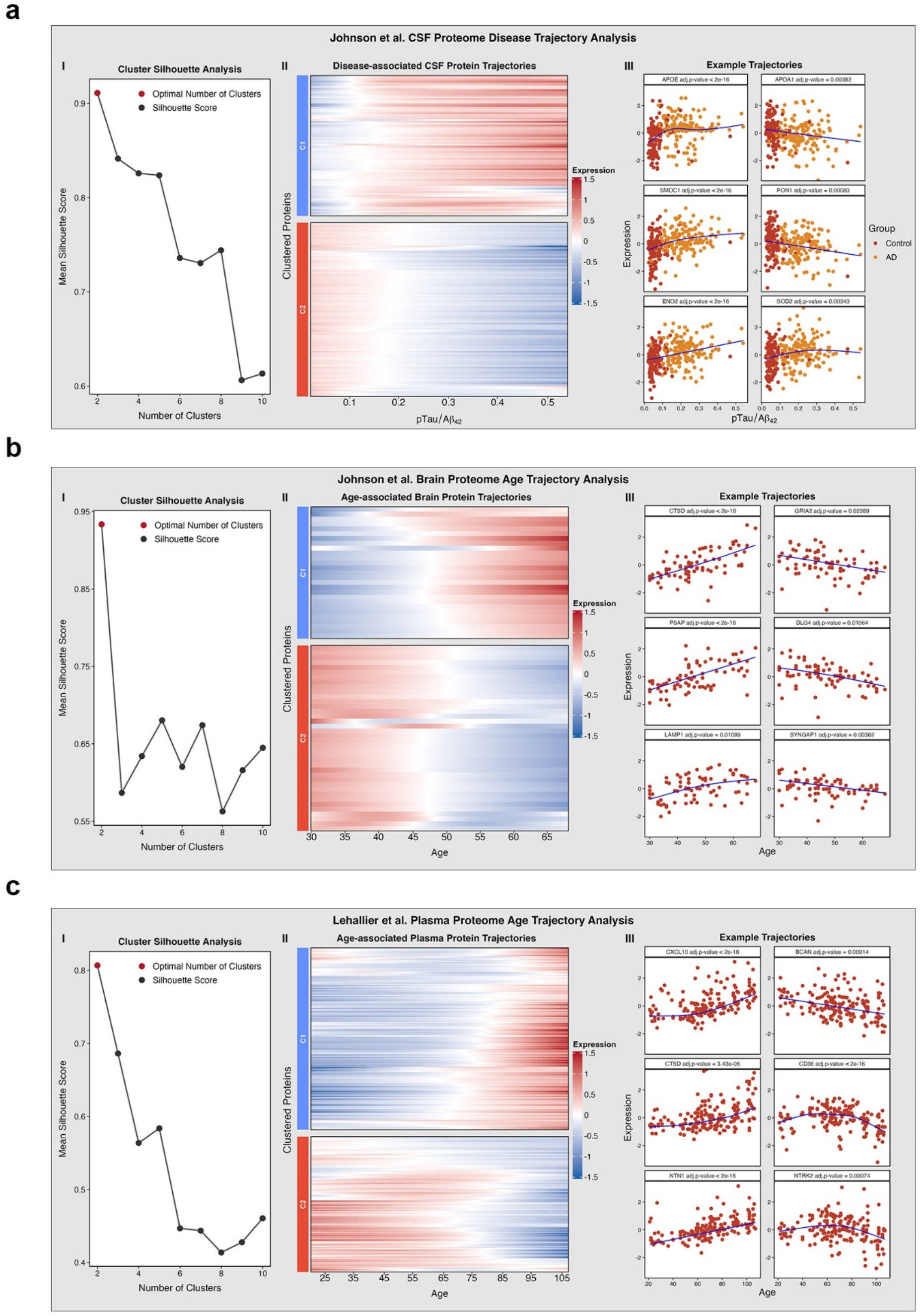
Proteomic trajectories across AD progression and aging. **(A)** Johnson et al. CSF proteomics data were modelled across the pTau/Aβ_42_ axis. (I) Silhouette analysis identified *k* = 2 as the optimal number of trajectory clusters. (II) Heatmap of significant CSF protein trajectories grouped into clusters C1–C2. (III) Representative protein trajectories, with points coloured by diagnostic group. **(B)** Johnson et al. brain proteomics data from healthy donors were modelled across age. (I) Silhouette analysis identified *k* = 2 as the optimal number of trajectory clusters. (II) Heatmap of significant brain protein trajectories grouped into clusters C1–C2. (III) Representative protein trajectories. **(C)** Lehallier et al. plasma proteomics data from healthy donors were modelled across age. (I) Silhouette analysis identified *k* = 2 as the optimal number of trajectory clusters. (II) Heatmap of significant plasma protein trajectories grouped into clusters C1–C2. (III) Representative protein trajectories.

**Figure S5.**
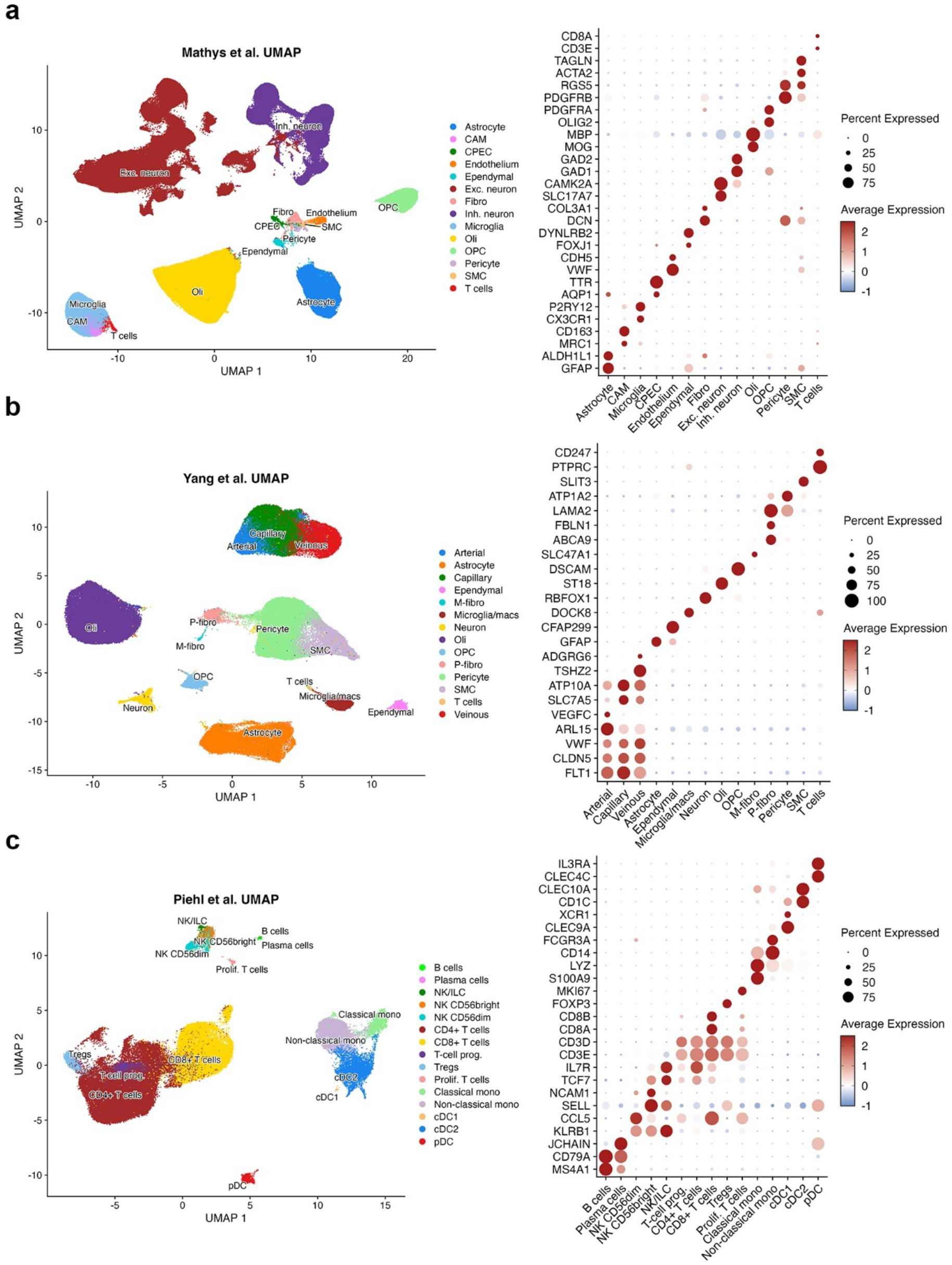
Single-cell and –nucleus gene expression atlases. (**A–C**) Brain parenchyma snRNA-seq atlas from Mathys et al. **(A)**, brain vasculature-enriched snRNA-seq atlas from Yang et al. **(B)**, and CSF-infiltrated immune cell scRNA-seq atlas from Piehl et al. **(C)**. UMAP embedding of nuclei coloured by annotated brain cell type (left panels) and dot plots showing canonical marker expression in annotated cell types (right panels).

**Figure S6.**
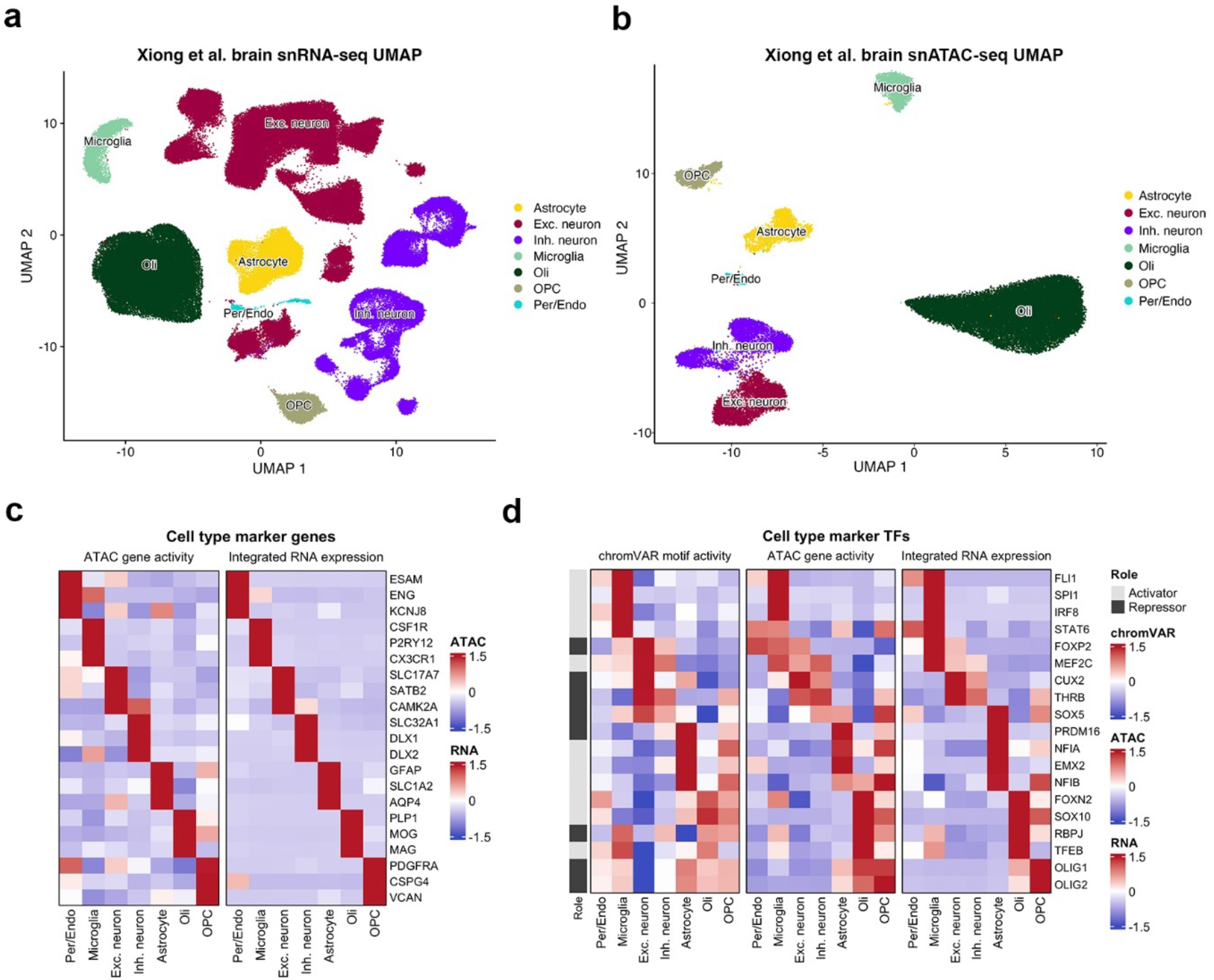
Cell type-resolved chromatin accessibility and gene expression atlas. (**A-B**) UMAP embedding of snRNA-seq **(A)** and snATAC-seq **(B)** nuclei from Xiong et al. **(C)** Heatmaps showing cell type marker genes from ATAC-inferred gene activity (left) and RNA expression (right). **(D)** Heatmaps showing cell type marker transcription factors (TFs) inferred from chromVAR motif activity (left), ATAC gene activity (middle), and RNA expression (right). For visualisation, chromVAR motif activity scores for repressor TFs were multiplied by –1.

**Figure S7.**
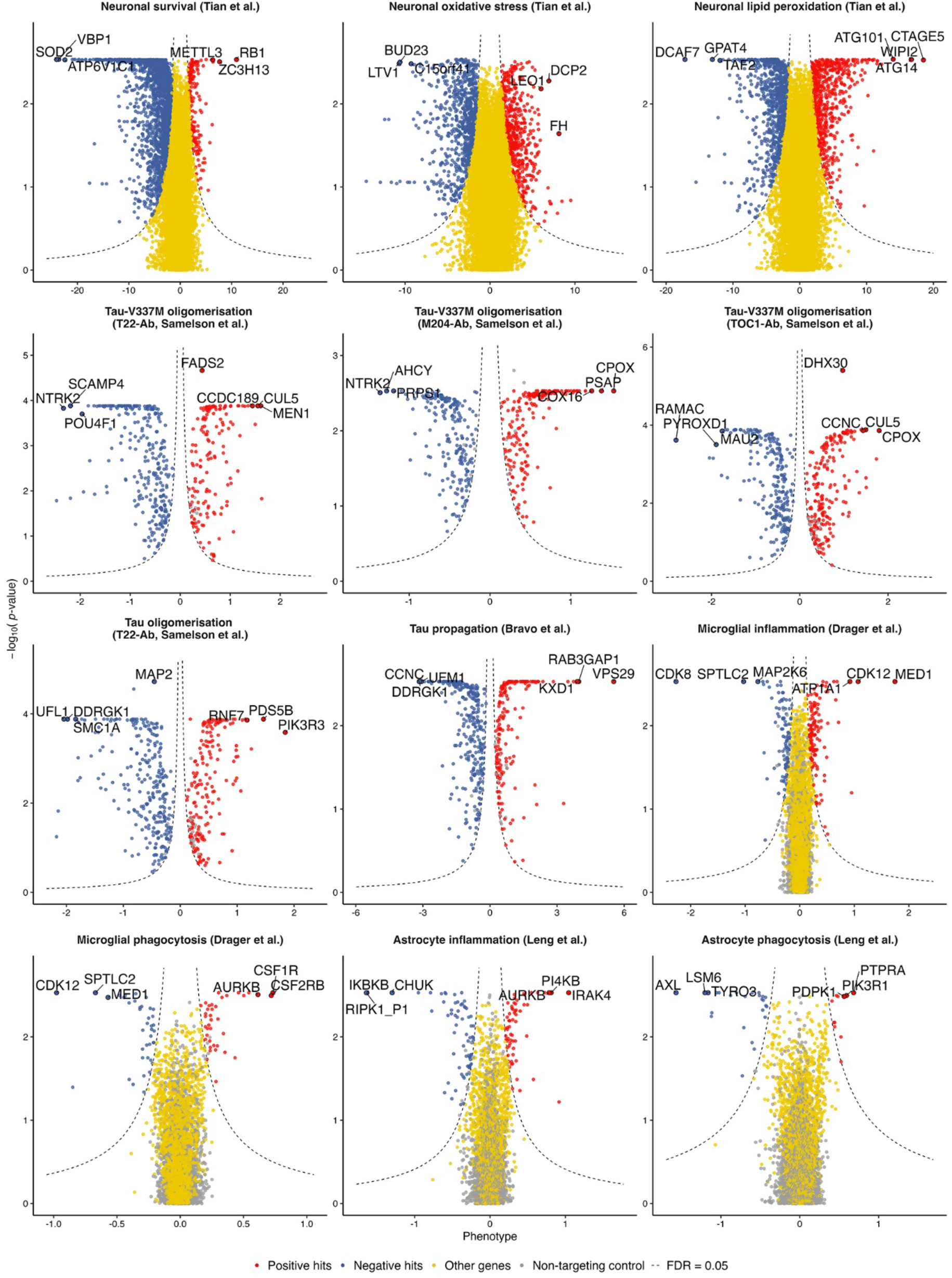
CRISPR screen-derived functional signatures. Volcano-style plots summarising AD-relevant CRISPRi/CRISPR screen results incorporated into the BRIDGE-AD signature library, including neuronal survival, oxidative stress and lipid peroxidation screens from Tian et al.; tau oligomerisation and tau propagation screens from Samelson et al. and Bravo et al.; and microglial or astrocytic inflammation and phagocytosis screens from Dräger et al. and Leng et al. The x-axis shows the reported phenotype score and the y-axis shows –log_10_(P value). Genes are coloured by screen-defined direction and significance: positive hits are shown in red, negative hits in blue, other assayed genes in yellow and non-targeting controls in grey. Dashed lines indicate the FDR = 0.05 threshold. Positive and negative hit sets from each screen were converted into functional perturbation signatures and integrated into BRIDGE-AD as evidence layers for machine learning-driven AD effector prioritisation.

**Figure S8.**
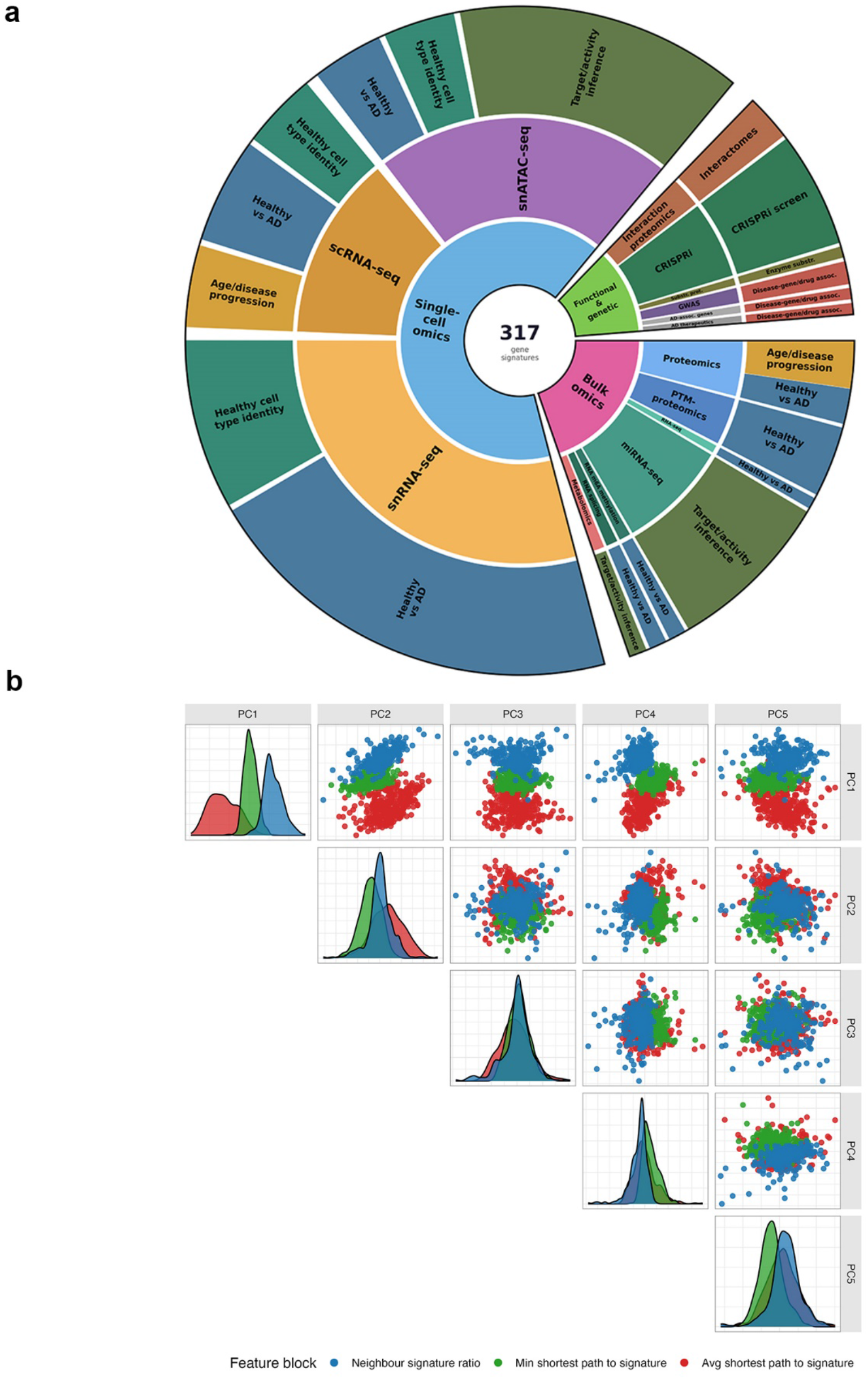
AD signature library and principal component analysis of the graph-derived gene-feature space. **(A)** Composition of the curated AD signature library. Sunburst plot summarising the 317 signatures used in this study, including single-cell and single-nucleus omics signatures (215/317), bulk omics signatures (65/317), as well as functional and genetic knowledge source-derived signatures (37/317). **(B)** Pairwise projections of the first five principal components, with each point representing one graph-derived proximity feature and coloured by feature block: ratio of neighbours (RN), minimum shortest path to signature (MSP), or average shortest path to signature (ASP). Diagonal panels show the distribution of each feature block along individual PCs, while off-diagonal panels show pairwise PC projections.

**Figure S9.**
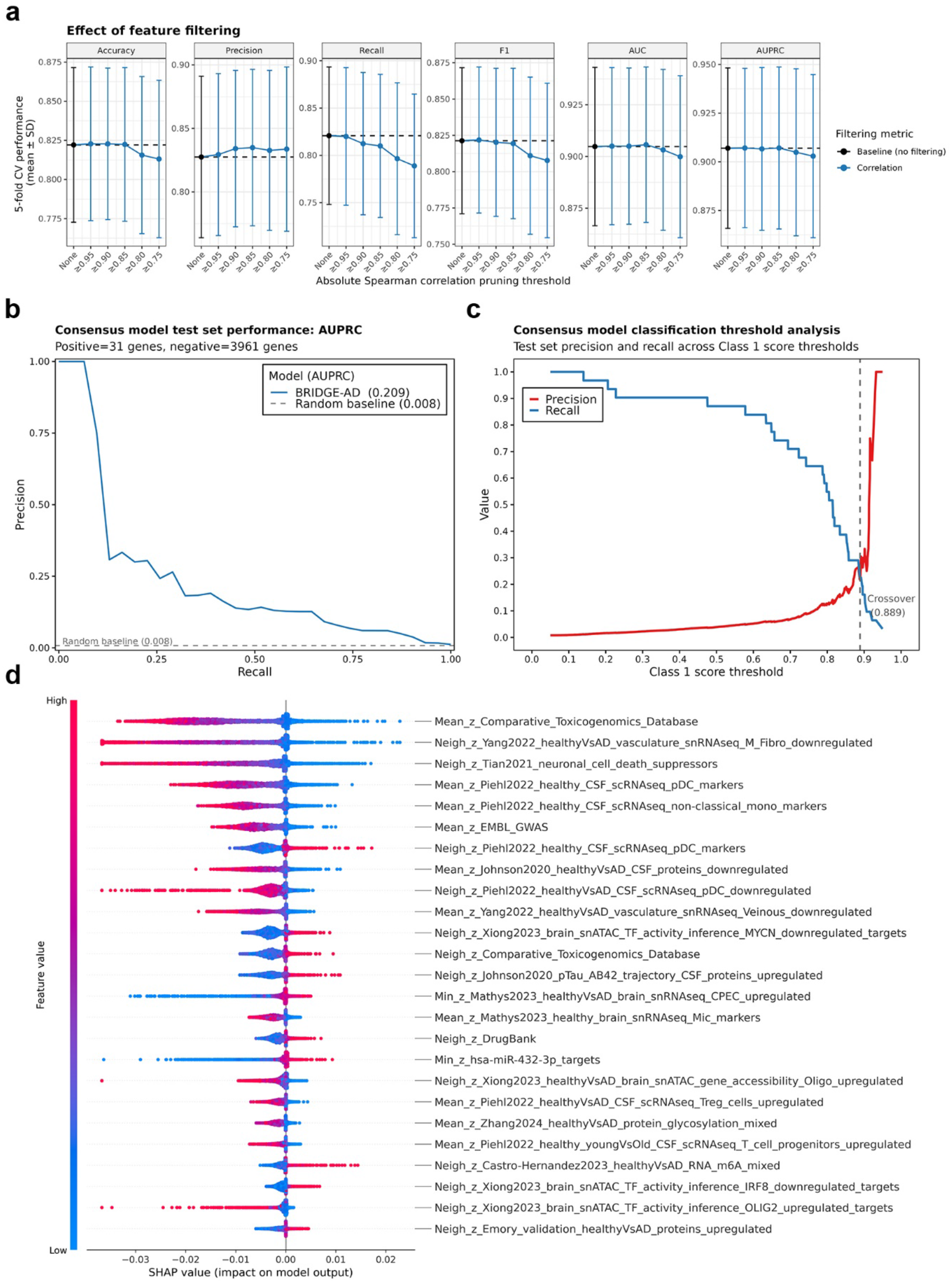
Machine learning pipeline evaluation: model optimisation, feature importance and test set performance. **(A)** Effect of correlation-based feature filtering on model performance. 5-fold cross-validation performance (mean ± SD) across feature matrices generated by pruning features with absolute Spearman correlations above the indicated threshold. The unfiltered baseline model, trained using all 951 features, is shown in black, with the dashed line indicating baseline performance for each metric. At each threshold, highly correlated feature pairs were grouped into redundancy clusters, from which one representative medoid feature was retained alongside all non-redundant singleton features. **(B)** Test set performance of the ensemble classifier on the full imbalanced held-out test set. Consensus predictions were generated by averaging the probability scores assigned to each held-out gene (Class 1 = 31 genes, Class 0 = 3,961 genes) across the 126 trained models. Model performance was then assessed across the full test split using AUPRC. **(C)** Test set precision–recall threshold analysis of the ensemble classifier. Precision and recall of the consensus ensemble predictions are shown across different Class 1 probability thresholds. The dashed vertical line indicates the crossover threshold of 0.889, at which precision and recall intersect. **(D)** SHAP-based feature importance of the ensemble classifier on the held-out test set. SHAP beeswarm plot showing the top 25 features ranked by mean absolute SHAP value after aggregating SHAP values across balanced test sets and model draws. Each point represents one gene, with the x-axis indicating the SHAP value and therefore the feature’s contribution to the model output. Positive SHAP values increase the predicted probability of the Class 1 AD effector label, whereas negative SHAP values decrease it. Point colour indicates the corresponding feature value, with blue denoting low values and pink denoting high values.

**Figure S10.**
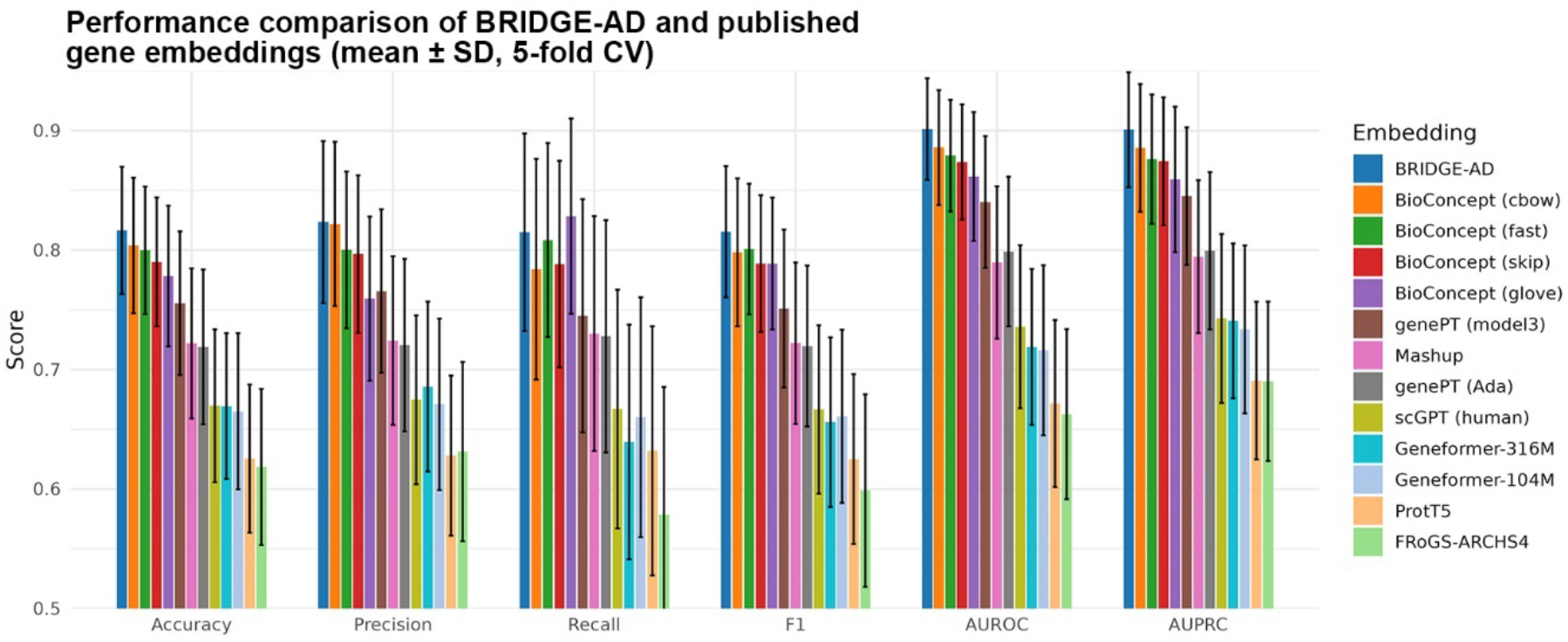
Training set performance across gene embeddings. 5-fold cross-validation performance (mean ± SD) of BRIDGE-AD and twelve recently published gene embeddings on the training split.

**Figure S11.**
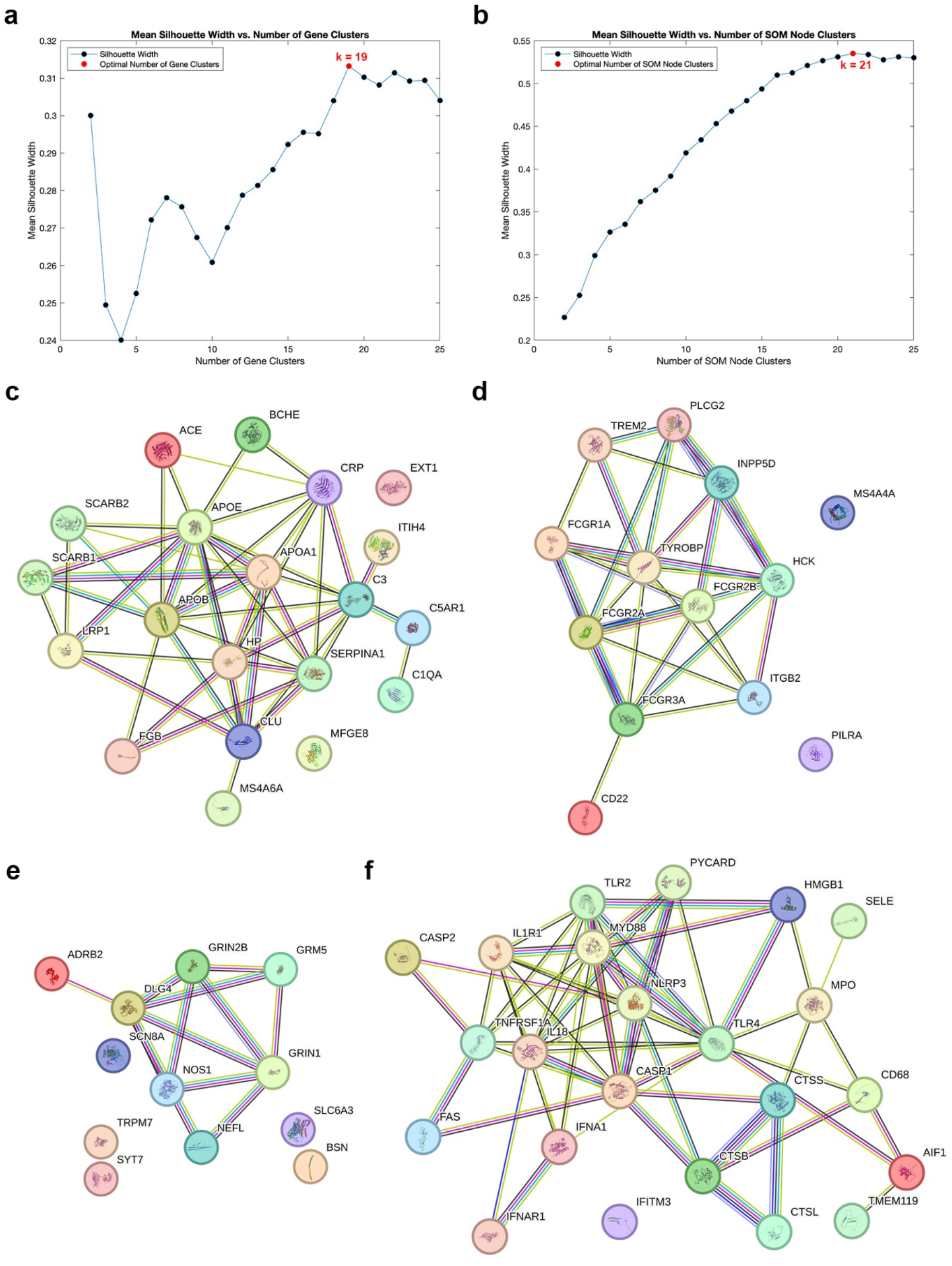
SOM QC analysis and example SOM clusters. **(A)** Silhouette-based selection of the optimal number of gene clusters, chosen by maximising the mean silhouette width (k = 19). **(B)** Silhouette-based selection of the optimal number of SOM node clusters, chosen by maximising the mean silhouette width (k = 21). Each node cluster was annotated with the dominant Reactome top-level pathway category among the pathways mapping to its nodes; clusters with the same dominant category were merged, yielding 20 final pathway regions. **(C-F)** SOM clusters representing lipid transport and complement (**C**, Cluster 4), myeloid immune receptor signalling (**D**, Cluster 7), synaptic transmission (**E**, Cluster 9), and innate immune activation (**F**, Cluster 2). For each example cluster, genes were visualised as STRING protein-protein interaction networks using a high-confidence interaction score cutoff of ≥ 700, with edges representing STRING-supported interactions among cluster members.

**Figure S12.**
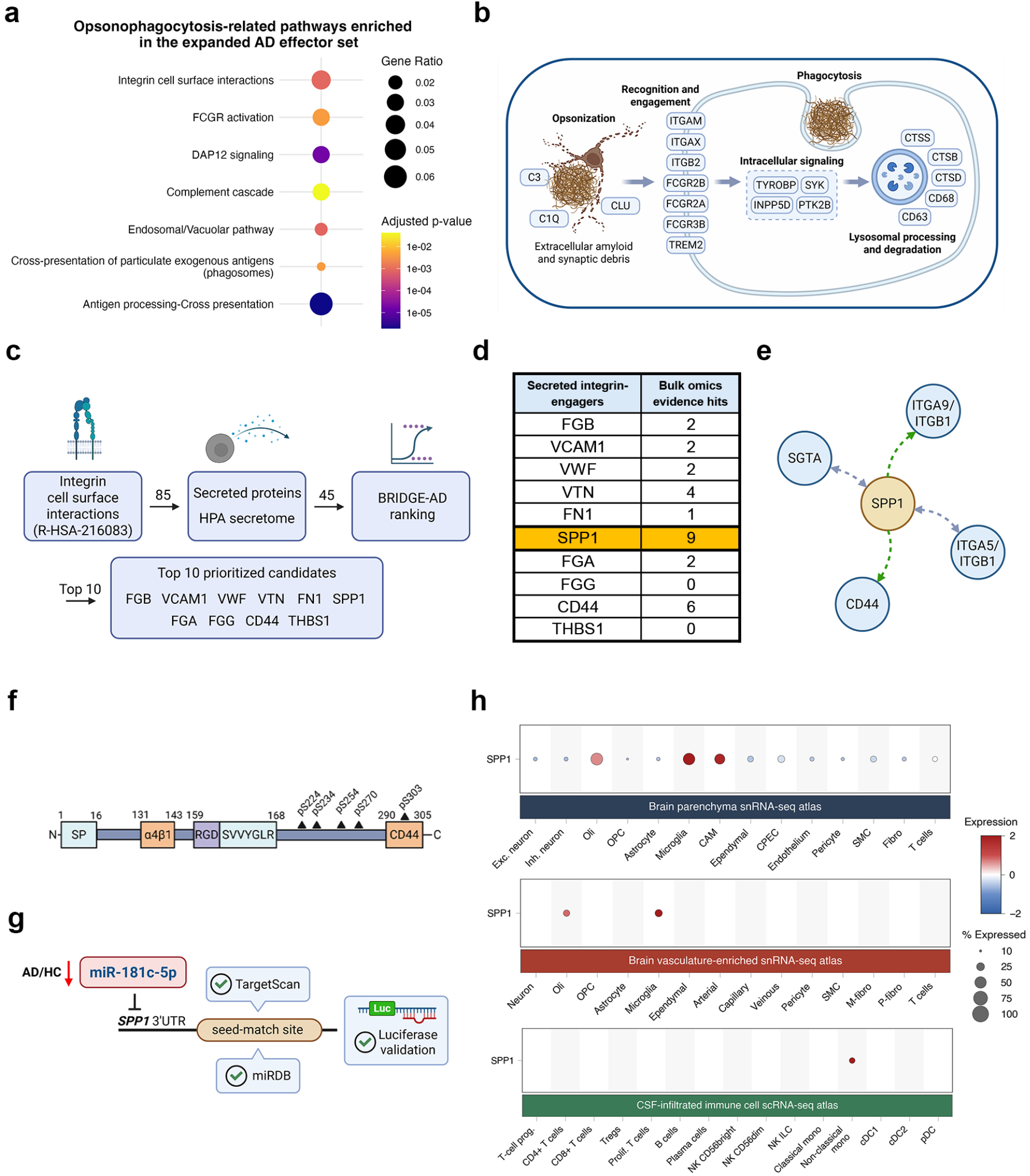
Expanded evidence trail supporting an SPP1-centred opsonophagocytosis hypothesis in AD. **(A)** Pathway enrichment analysis of the expanded AD effector set, highlighting opsonophagocytosis-related pathways. **(B)** Top prioritised AD effectors and selected Class 1 genes form a myeloid opsonophagocytosis-related programme comprising opsonisation, recognition and engagement, intracellular signalling, phagocytosis, and lysosomal processing and degradation modules. **(C)** Filtering workflow used to identify extracellular effectors capable of engaging the opsonophagocytosis programme. Genes annotated to Reactome ‘Integrin cell surface interactions’ were restricted to Human Protein Atlas (HPA) secretome proteins and ranked by BRIDGE-AD probability, nominating the top 10 prioritised secreted integrin-engaging candidates. **(D)** Summary of integrated bulk omics evidence for the top 10 nominated secreted integrin-engaging candidates, highlighting SPP1 as the candidate with the broadest bulk omics support. **(E)** OmniPath analysis of SPP1-centred molecular interactions, showing direct SPP1-associated targets including integrin receptor complexes. **(F)** Schematic of SPP1 protein domain architecture and AD-associated phosphorylation sites, including the signal peptide, integrin-binding regions, the RGD and SVVYGLR motifs, CD44-associated region, and C-terminal phosphosites dysregulated in AD. **(G)** miR-181c-5p as a candidate upstream regulator of *SPP1*, showing decreased miR-181c-5p abundance in AD and evidence layers for *SPP1* targeting. **(H)** SPP1 expression across cell types captured in the integrated brain parenchyma, brain vasculature-enriched, and CSF-infiltrated immune cell single-cell and single-nucleus RNA-seq atlases, highlighting *SPP1* expression in microglia, CNS-associated macrophages, and non-classical monocytes.

**Figure S13.**
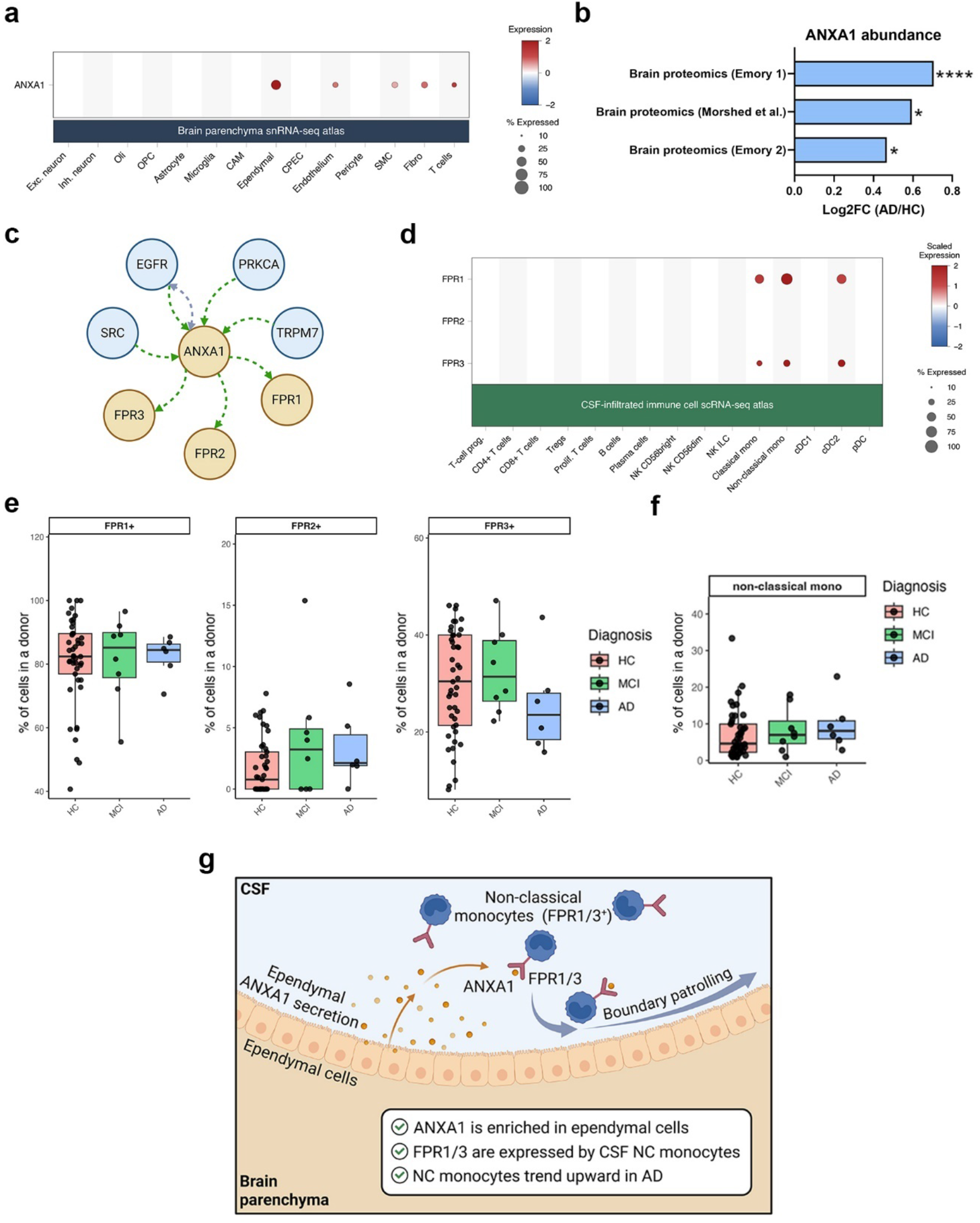
ANXA1–FPR evidence supporting a brain-boundary monocyte recruitment hypothesis in AD. **(A)** *ANXA1* expression across major cell types in the brain parenchyma snRNA-seq atlas, showing enrichment in ependymal cells. Dot size indicates the percentage of cells expressing *ANXA1*, and colour indicates scaled mean expression. **(B)** AD-associated changes in ANXA1 abundance across bulk brain proteomics datasets, showing increased ANXA1 protein abundance in AD relative to healthy controls. Asterisks indicate FDR-adjusted significance: *FDR < 0.05 and ****FDR < 0.0001. **(C)** OmniPath analysis of ANXA1-centred molecular interactions, showing direct targeting of formyl peptide receptors (FPRs) implicated in leukocyte chemotaxis, including FPR1, FPR2, and FPR3. **(D)** Expression of *FPR1*, *FPR2*, and *FPR3* across CSF-infiltrated immune cell populations in the Piehl et al. dataset, showing enriched *FPR1* and *FPR3* expression in non-classical monocytes. Dot size indicates the percentage of cells expressing each gene, and colour indicates scaled mean expression. **(E)** Diagnosis-stratified abundance of FPR1^+^, FPR2^+^, and FPR3^+^ CSF-infiltrated immune cell populations. **(F)** Quantification of non-classical monocyte abundance across diagnostic groups in the CSF-infiltrated immune cell dataset. **(G)** Proposed model in which *ANXA1* secretion promotes FPR1/3-dependent recruitment or patrolling of non-classical monocytes in the CSF, providing a potential source of SPP1 capable of engaging integrin-linked phagocytic programmes in AD. **(E–F)** Boxes represent the interquartile range, centre lines indicate medians, whiskers extend to values within 1.5×IQR, and points represent individual donors. HC, healthy control; IQR, interquartile range; MCI, mild cognitive impairment; NC, non-classical.

**Figure S14.**
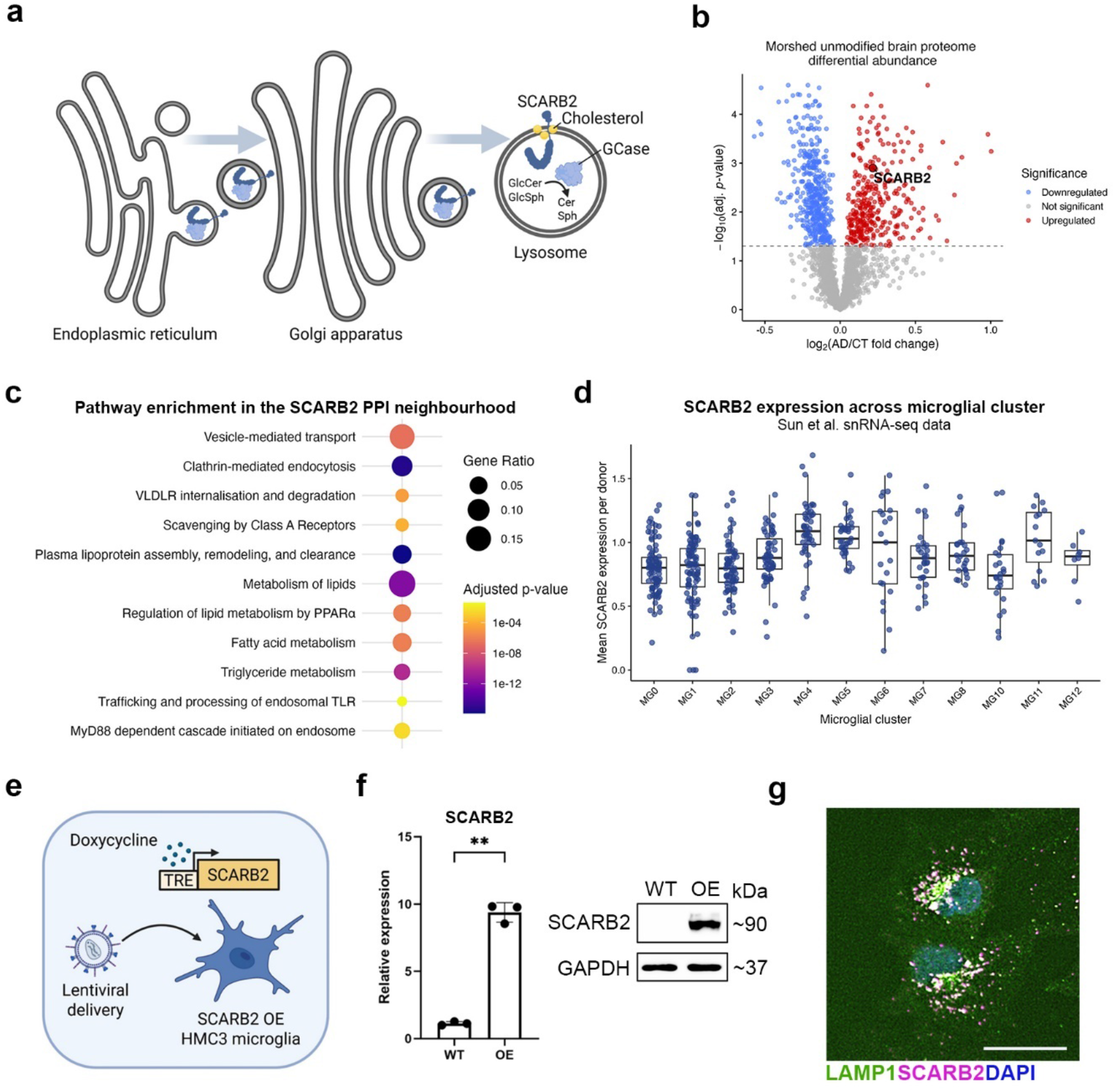
SCARB2 background, systems-level context, and validation of the HMC3 overexpression model. **(A)** Schematic overview of SCARB2/LIMP-2 biology. SCARB2 traffics through the endoplasmic reticulum and Golgi apparatus to the lysosome, where it delivers GCase and functions as a lysosomal membrane protein involved in lysosomal lipid-related processes, including cholesterol transport. GlcCer, glucosylceramide; GlcSph, glucosylsphingosine. **(B)** Volcano plot of the Morshed et al. unmodified brain proteome differential abundance analysis, highlighting increased SCARB2 protein abundance in AD. **(C)** Pathway enrichment analysis of the SCARB2 PPI neighbourhood. **(D)** Mean *SCARB2* expression across microglial subtypes in the Sun et al. brain parenchyma snRNA-seq atlas using microglial clusters annotated in the original study. **(E)** Experimental strategy used to establish *SCARB2* overexpressing HMC3 microglia through lentiviral delivery of a doxycycline-inducible *SCARB2* expression cassette. **(F)** Validation of *SCARB2* overexpression in HMC3 microglia by RT-qPCR and immunoblotting. RT-qPCR data are presented as mean ± SD. **p < 0.01 by a two-sided unpaired Welch’s t-test. N = 3 independent biological replicates. **(G)** Representative immunofluorescence imaging showing colocalisation of overexpressed SCARB2 protein with LAMP1^+^ lysosomal compartments in HMC3 microglia.

**Figure S15.**
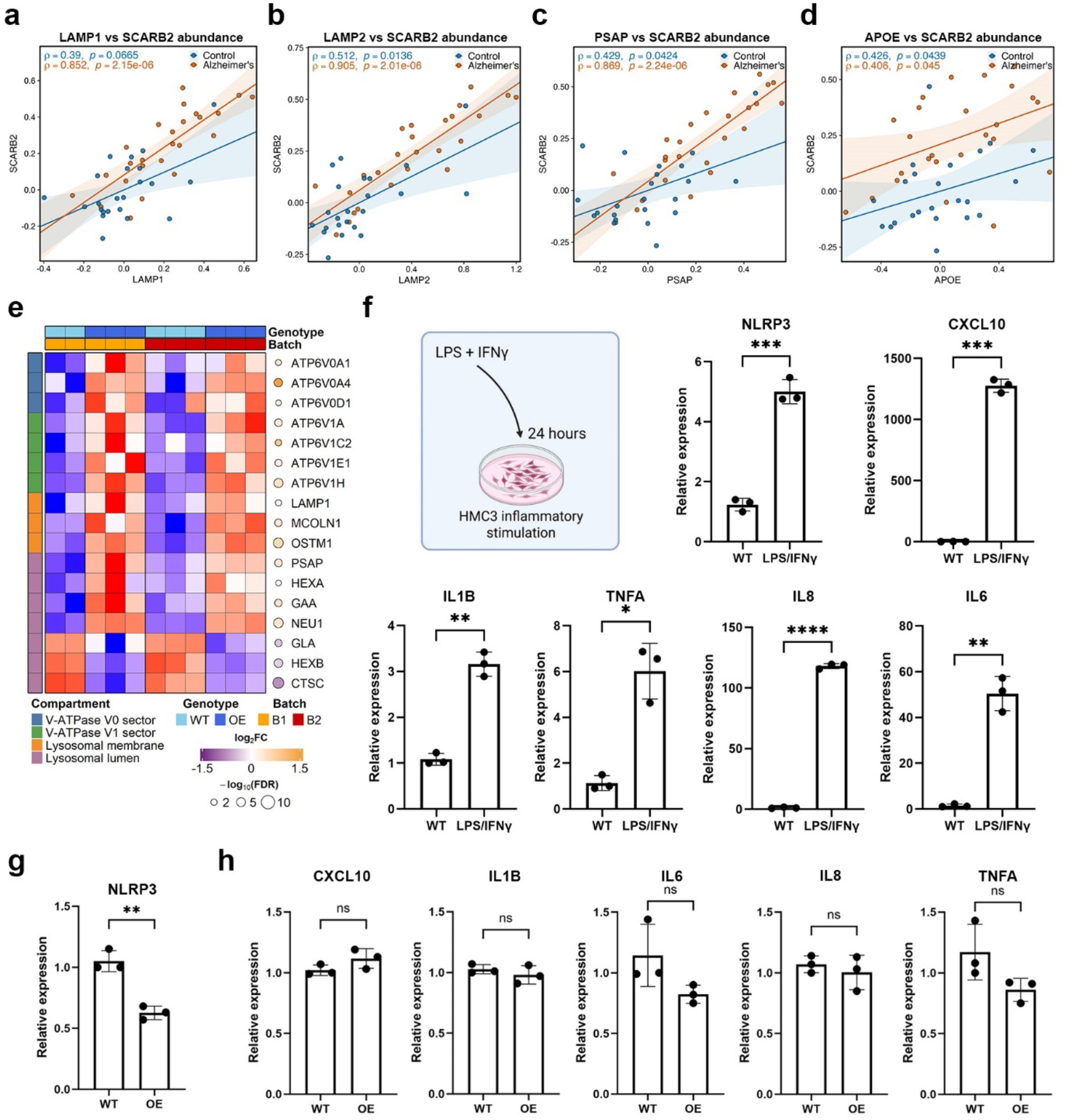
Cross-context support for SCARB2-associated lysosomal, lipid-processing, and inflammatory-state remodelling. (A-D) Diagnosis-stratified correlations between SCARB2 protein abundance and lysosomal or lipid-processing proteins in human brain proteomics data from Morshed et al., including LAMP1 **(A)**, LAMP2 **(B)**, PSAP **(C)**, and APOE **(D)**. Donors with AD and healthy controls are shown separately to assess disease-associated coupling between SCARB2 and each comparator protein. Solid lines show the group-specific linear regression fits, with shaded regions indicating the corresponding 95% confidence intervals. Spearman correlation coefficients and associated p-values are shown for each diagnostic group. **(E)** Focused RNA-seq heatmap of lysosomal genes altered by *SCARB2* overexpression in HMC3 microglia. Genes are grouped by lysosomal compartment or function, including V-ATPase V0, V-ATPase V1, lysosomal membrane, and lysosomal lumen annotations. N_WT_ = 5, N_OE_ = 6 independent biological replicates. **(F)** Inflammatory stimulation paradigm and positive-control induction of inflammatory genes in WT HMC3 microglia. WT HMC3 cells were stimulated with LPS and IFNγ for 24 hours, and expression of pro-inflammatory genes was assessed by RT-qPCR. N = 3 independent biological replicates. **(G)** NLRP3 expression in WT and *SCARB2*-overexpressing HMC3 microglia following inflammatory stimulation. N = 3 independent biological replicates **(H)** Expression of pro-inflammatory cytokines in WT and *SCARB2*-overexpressing HMC3 microglia following inflammatory stimulation. N = 3 independent biological replicates. (**G, H**) RT-qPCR data are presented as mean ± SD. **p < 0.01 by a two-sided unpaired Welch’s t-test; ns, not significant.

**Figure S16.**
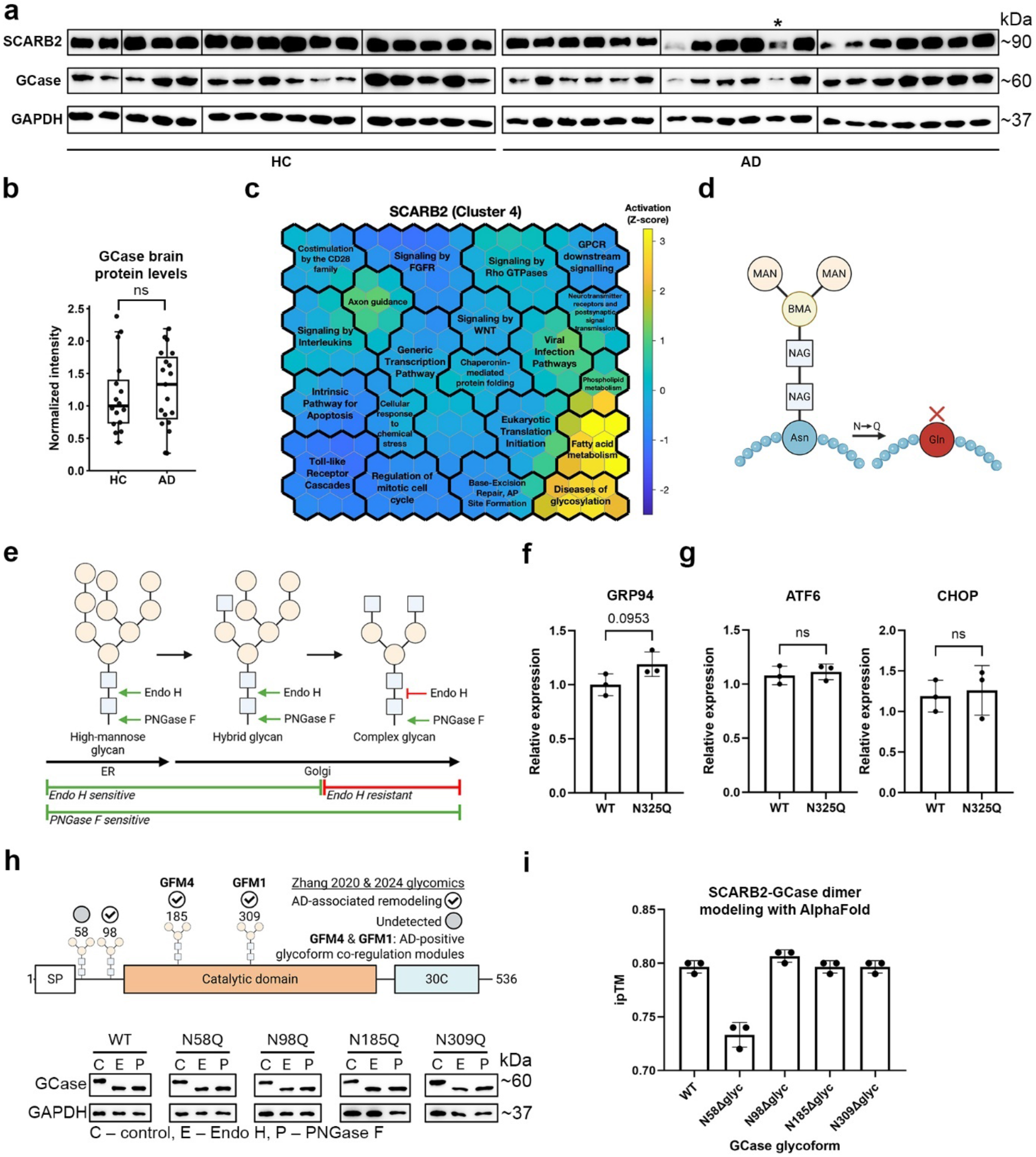
Supplementary context and extended glycosylation analyses for the SCARB2–GCase axis. (**A**) SCARB2 and GCase protein abundance in primary human brain tissue from donors with AD and healthy controls. GAPDH is shown as a loading control. The lane marked with an asterisk indicates the sample in which SCARB2 appeared as two closely migrating bands, suggesting that multiple SCARB2 maturation or glycoform states may be present in AD. **(B)** Tukey box-and-whisker plots showing median-normalised GCase protein abundance in healthy control and AD brain samples; boxes indicate the interquartile range, centre lines indicate medians, whiskers extend to values within 1.5×IQR, and points represent individual donors. Mean protein abundance was compared by Welch’s t-test and was not significantly different between diagnostic groups. **(A–B)** N_HC_ = 16, N_AD_ = 19 brain tissue donors. **(C)** SCARB2 SOM grid showing peak activation in regions enriched for fatty acid metabolism and diseases of glycosylation. **(D)** Schematic of the mutagenesis strategy used to remove individual N-glycans by substituting asparagine with glutamine at validated glycosylation sites. **(E)** Schematic of glycan maturation and glycosidase sensitivity. High-mannose glycans are Endo H-sensitive, complex glycans are Endo H-resistant, and all N-glycans are PNGase F-sensitive, enabling assessment of ER-to-Golgi maturation. **(F)** *GRP94* expression in WT and *SCARB2-N325Q*-overexpressing HMC3 microglia, showing a trend towards increased ER chaperone engagement. **(G)** Expression of ER stress or unfolded protein response markers *ATF6* and *CHOP* in WT and *SCARB2-N325Q*-overexpressing HMC3 microglia. **(F–G)** N = 3 independent biological replicates. **(H)** (Upper) Schematic and glycoproteomic summary of GCase N-glycosylation, including detected and undetected glycosylation sites in Zhang et al. (2020) and Zhang et al. (2024) glycoproteomics datasets, AD-associated remodelling, and assignment to AD-positive GFM4 and GFM1 glycoform co-regulation modules. (Lower) Biochemical analysis of WT GCase and single-site GCase N→Q glycoforms using control buffer, Endo H, and PNGase F digestion followed by immunoblotting. GAPDH is shown as a loading control. **(I)** Interface predicted template modelling (ipTM) scores for SCARB2–GCase complexes containing WT or mutant GCase glycoforms, based on *AlphaFold* modelling. N = 3 random seeds. **(F**–**G)** RT-qPCR data are presented as mean ± SD. ns, not significant by a two-sided unpaired Welch’s t-test. (**I**) Data are presented as mean ± SD.

**Extended Data Figure 1.**
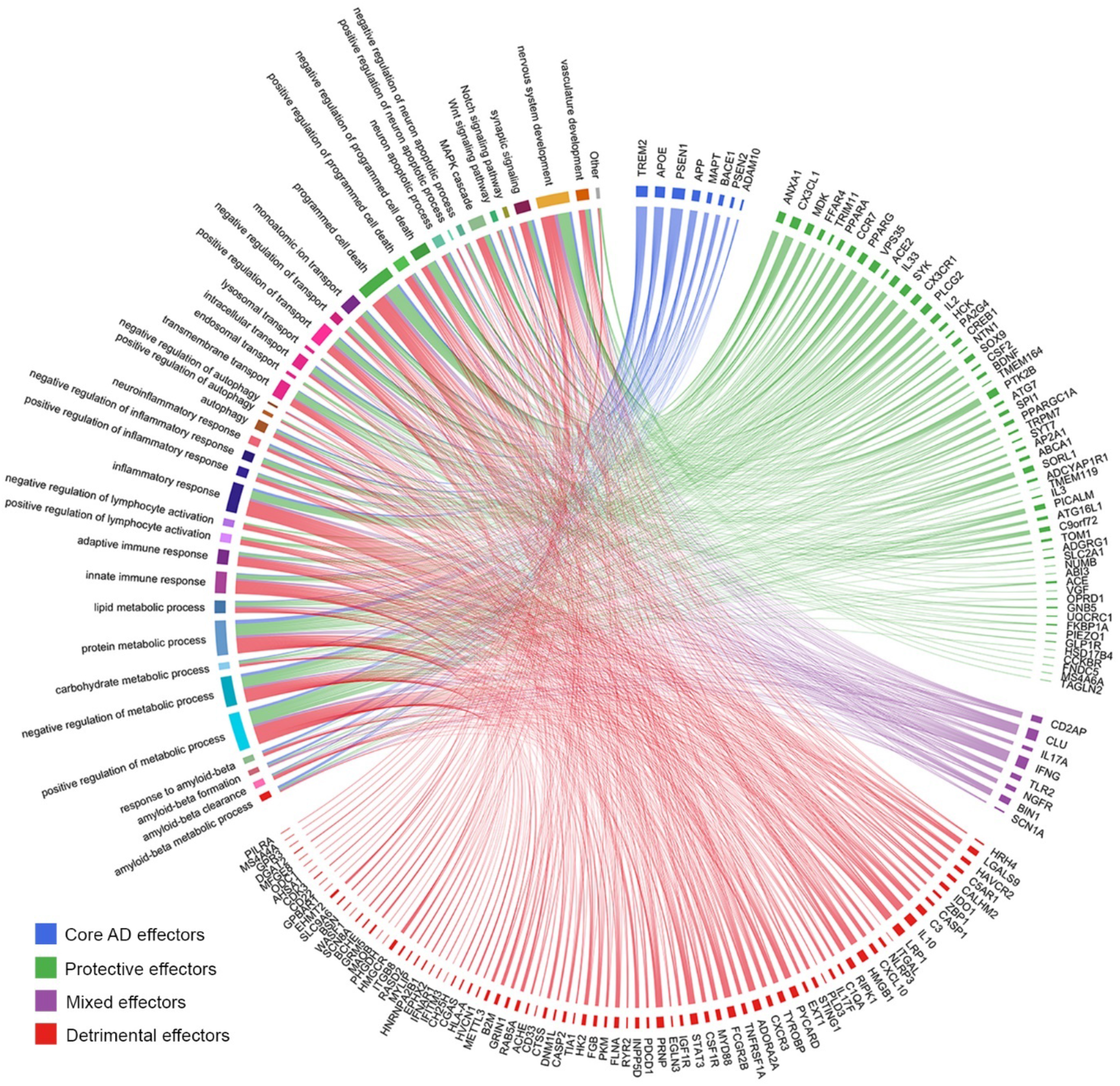
Functional enrichment of Class 1 effectors. Circos plot linking GO biological processes to Class 1 effectors, grouped as core (blue), protective (green), detrimental (red), or mixed (purple) AD genes.

**Extended Data Figure 2.**
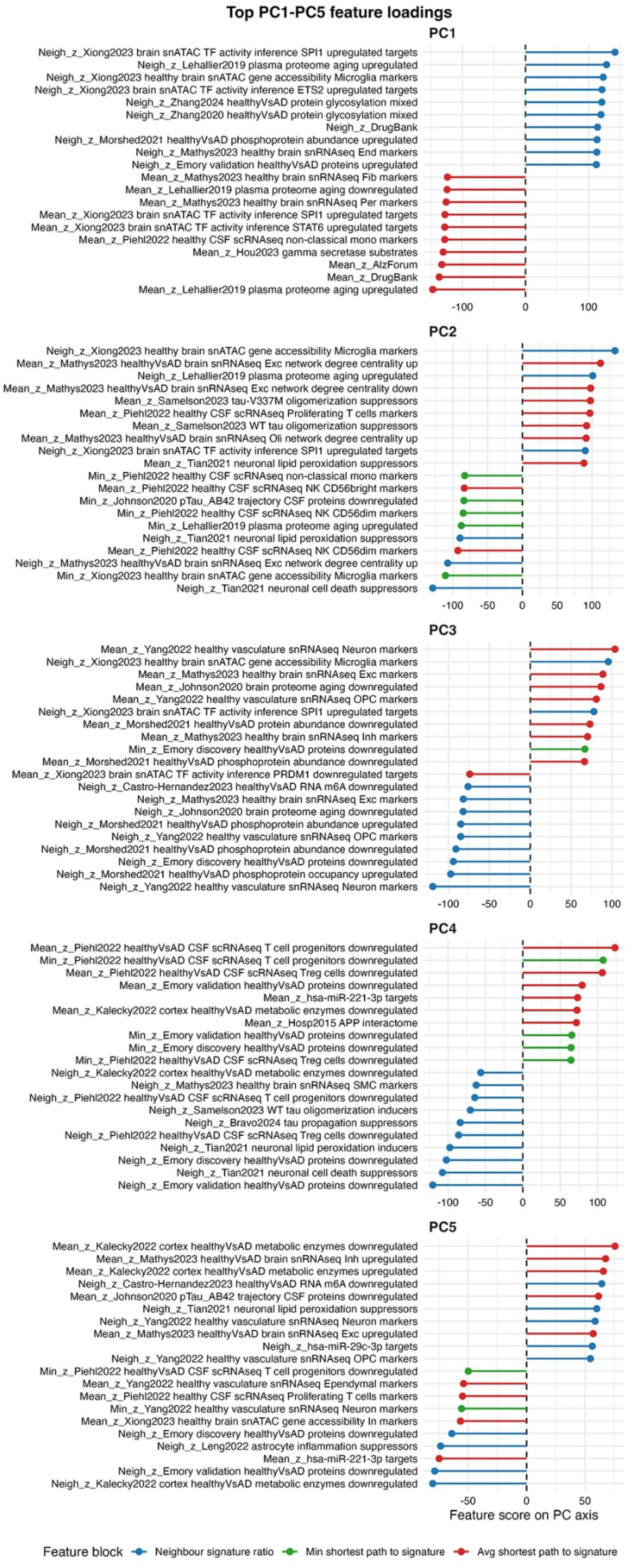
Top PC1–PC5 loadings of the graph-derived gene-feature space. Lollipop plots showing the top positive and negative feature loadings for PCs 1–5, annotated by the corresponding signature name and proximity metric.

## Supplementary Discussion

Across bulk omics datasets, we prioritised proteomics datasets, as they represent the effector layer of gene expression (**Figure S3A**). To that end, we incorporated brain proteomics of AD donors and healthy controls from two Emory cohorts^15^ and Morshed et al.^12^, the latter of which also mapped post-translational protein phosphorylation. We also integrated a CSF proteome of AD donors and healthy controls from Johnson et al.^11^, as well as two brain glycoproteomes from Zhang et al., 2020^13^, and Zhang et al., 2024^14^. At the transcriptome level, we included ROSMAP brain RNA-seq and miRNA-seq datasets^41,43^. Differential expression analysis (DEA) revealed expected differentially expressed genes (DEGs) and differentially abundant proteins (DAPs), such as increased APOE protein levels in the CSF of AD donors and increased SMOC1 protein levels in the brain proteomes of AD donors. For differentially expressed miRNAs, we generated target-gene signatures by mapping predicted targets using miRDB^44^ and TargetScan^45^, enabling miRNA-level dysregulation to be represented within the protein-coding BRIDGE-AD feature space.

We also obtained brain and plasma metabolomics datasets from Kalecky et al. (**Figure S3B-C**)^46^. To incorporate these datasets into our PPI network-based framework, we inferred enzyme activity programmes from differentially abundant metabolites using the ocEAn package^75^ (see **Methods**). Differential enzyme activity between the metabolomes of AD donors and healthy controls is represented by the normalised enrichment score (NES) and reveals top dysregulated enzymatic functions in AD (**Figure S3B-C, Panel I**). In the brain, aldehyde dehydrogenase (ALDH) and phosphoethanolamine/phosphocholine phosphatase 1 (PHOSPHO1) represent the top increased and decreased enzymatic activities in AD, respectively. In the plasma, top increased and decreased enzymatic activities are represented by betaine-homocysteine S-methyltransferase (BHMT) and a broad class of oxidoreductases (EC:1.5.99), respectively. Metabolites representing the enzymatic activities of ALDH, PHOSPHO1, BHMT, and EC:1.5.99 enzymes are shown in **Panels II-III**.

To represent temporal progression of AD, we reanalysed a CSF proteomics dataset from Johnson et al.^11^ We modelled protein changes as a function of the pTau/Aβ_42_ ratio, which is a well-established biomarker of AD severity (**Figure S4A**). We identified 179 CSF proteins that significantly change along the course of AD progression. Silhouette analysis supported a two-cluster solution, corresponding to proteins that increase or decrease along the pTau/Aβ_42_ axis (**Figure S4A, panels I-II**). Example trajectories representing emerging CSF biomarkers of AD are shown in **panel III**. Additionally, we incorporated two datasets of brain (Johnson et al.^11^) and plasma (Lehallier et al.^17^) proteomics profiling in healthy donors across age (**Figure S4B-C**). Silhouette analysis supported two-cluster solutions for both datasets, corresponding to proteins that increase or decrease with age (**Figure S4B-C, panels I-II**). Representative proteins and their trajectories are shown in **panels III**. Each disease– or age-associated protein trajectory cluster was converted into a signature and incorporated into the feature space of BRIDGE-AD providing an orthogonal evidence layer that captures temporal molecular programmes of AD progression.

To derive cell type-resolved gene expression signatures for the BRIDGE-AD feature space, we analysed three single-cell and –nucleus atlases: a brain snRNA-seq dataset from Mathys et al.^19^, a brain vasculature snRNA-seq dataset from Yang et al.^22^, and a CSF-infiltrated immune cell scRNA-seq dataset from Piehl et al.^20^ (**Figures S5**). The brain parenchyma snRNA-seq dataset resolved major CNS cell types, including excitatory and inhibitory neurons, astrocytes, oligodendrocytes, oligodendrocyte progenitor cells (OPCs), microglia, vascular cells (endothelial cells, pericytes, and smooth muscle cells), and other cell types (**Figure S5A**). The brain vasculature-enriched snRNA-seq (VINE-seq) resolved major vascular cell types, including endothelial subtypes (arterial, capillary, and venous endothelial cells), pericytes, smooth muscle cells, and associated parenchymal, immune, and stromal cell types (**Figure S5B**). The CSF-infiltrated immune cell scRNA-seq dataset resolved major lymphoid and myeloid lineages of peripheral immune cells, including CD4^+^ and CD8^+^ T cells, regulatory T cells (T_reg_), natural killer (NK) cell subtypes, B cells and plasma cells, monocyte subtypes, and dendritic cell subtypes (**Figure S5C**). Incorporation of the brain vasculature-enriched and the CSF immune cell datasets into BRIDGE-AD broadens our model beyond the brain parenchyma and captures disease-relevant compartments and cell types that are often underrepresented in standard brain atlases of gene expression. This broader coverage of the neurovascular compartment and the peripheral immune cell-CSF interface enables prioritisation and mechanistic investigation of disease effectors related to vascular dysfunction and peripheral immune cell infiltration that are both emerging themes in AD research.

For each cell type annotated in these atlases, we generated two types of gene signatures for the BRIDGE-AD feature space. First, we identified cell type-specific marker genes using nuclei from healthy donors to capture cellular identity programmes of the key cell populations involved in AD. Second, we performed DEA to identify genes significantly upregulated or downregulated in each cell type in AD donors relative to healthy controls. Lists of cell type upregulated or downregulated DEGs were used as cell context-specific disease signatures in the constructed feature space. In addition, the CSF immune cell dataset included young and aged healthy donors, allowing us to derive age-associated transcriptional signatures across immune cell types, while the brain parenchyma snRNA-seq dataset was also used to infer AD-associated cell type-specific interactome rewiring using SCINET.

To incorporate underlying regulatory programmes of gene expression in AD, we reanalysed a pre-processed paired single-nucleus ATAC-seq (snATAC-seq) and snRNA-seq dataset from Xiong et al. derived from the prefrontal cortex of ROSMAP participants (**Figure S6**)^21^. UMAP embedding of snRNA-seq (**Figure S6A**) and snATAC-seq (**Figure S6B**) nuclei, labelled using cell type annotations from Xiong et al., resolved major brain parenchyma cell types in both modalities, including excitatory and inhibitory neurons, oligodendrocytes, OPCs, astrocytes, microglia, and vascular cells. We then used these paired data to derive several types of regulatory signatures for the BRIDGE-AD feature space. First, we used healthy donor data to identify cell type marker genes by differential accessibility analysis (**Figure S6C**). This yielded one chromatin accessibility-derived marker signature for each major cell type. Second, we used AD and control donor data to identify differentially accessible genes in disease, generating two AD-associated chromatin accessibility signatures (increased and decreased accessibility) for each major cell type. Third, we integrated chromVAR motif accessibility, ATAC gene activity, and snRNA-seq expression analyses to infer cell type-specific marker transcription factors (TFs) (**Figure S6D**, see **Supplementary Methods**). We also used the same framework to identify differentially active TFs in AD donors as compared to healthy controls within each cell type. For each inferred cell type-enriched or differentially active TF, we retrieved its target genes from the CollecTRI^76^ regulatory network database and generated two regulatory signatures: one comprising positively regulated targets and the other negatively regulated targets. Together, bulk molecular profiling, trajectory, cell type-resolved, regulatory, functional and genetic evidence layers formed the final BRIDGE-AD signature library of 317 standardised entries, comprising 215 single-cell and single-nucleus omics signatures, 65 bulk multi-omics signatures, and 37 signatures derived from functional or genetic knowledge sources.

## Data availability

The raw RNA-sequencing data generated in this study have been deposited in Zenodo under DOI: 10.5281/zenodo.21973503.

Complete AD signatures are available at https://github.com/Greta-B/BRIDGE-AD.

Data obtained from the AD Knowledge Portal. The results published here are in whole or in part based on data obtained from the AD Knowledge Portal.

The Religious Orders Study and Memory and Aging Project (ROSMAP) Study. The data available in the AD Knowledge Portal would not be possible without the participation of research volunteers and the contribution of data by collaborating researchers. Study data were provided by the Rush Alzheimer’s Disease Center, Rush University Medical Center, Chicago. Data collection was supported through funding by NIA grants P30AG10161 (ROS), R01AG15819 (ROSMAP RNAseq), R01AG17917 (MAP), R01AG30146, R01AG36836 (RNAseq), U01AG46161 (TMT proteomics), U01AG61356 (ROSMAP AMP-AD), P30AG072975, the Illinois Department of Public Health (ROSMAP). Additional phenotypic data can be requested at www.radc.rush.edu.

ROSMAP TMT Proteomics. Study data were provided through the Accelerating Medicine Partnership for AD (U01AG046161 and U01AG061357) based on samples provided by the Rush Alzheimer’s Disease Center, Rush University Medical Center, Chicago. Data collection was supported through funding by NIA grants P30AG10161, R01AG15819, R01AG17917, R01AG30146, R01AG36836, U01AG32984, U01AG46152, the Illinois Department of Public Health, and the Translational Genomics Research Institute.

MIT ROSMAP Single-Nucleus Multiomics. Study data were generated from postmortem brain tissue provided by the Religious Orders Study and Rush Memory and Aging Project (ROSMAP) cohort at Rush Alzheimer’s Disease Center, Rush University Medical Center, Chicago. This work was supported in part by the Cure Alzheimer’s Fund, NIH grants AG058002, AG062377, NS110453, NS115064, AG062335, AG074003, NS127187, MH119509, HG008155 (M.K.), RF1AG062377,

RF1 AG054321, RO1 AG054012 (L.-H.T.) and the NIH training grant GM087237 (to C.A.B.). ROSMAP is supported by P30AG10161, P30AG72975, R01AG15819, R01AG17917. U01AG46152, U01AG61356.

Bulk brain RNA-seq (syn12104384)^41^ and brain miRNA array data (syn3387325)^43^ from ROSMAP; brain proteomics from the Emory cohorts (syn20824864)^15,38^; cerebrospinal fluid proteomics (syn21441787)^11^; brain proteomics across aging from the Johns Hopkins cohort (syn21441775)^11^; brain parenchyma snRNA-seq from ROSMAP (syn52383412; http://compbio.mit.edu/ad_multiregion/)^19^; and paired single-nucleus ATAC-seq and RNA-seq from ROSMAP (syn52293417; http://compbio.mit.edu/ad_epigenome/)^21^ are available via the AD Knowledge Portal (https://adknowledgeportal.org). The AD Knowledge Portal is a platform for accessing data, analyses, and tools generated by the Accelerating Medicines Partnership (AMP-AD) Target Discovery Program and other National Institute on Aging (NIA)-supported programs to enable open-science practices and accelerate translational learning. The data, analyses and tools are shared early in the research cycle without a publication embargo on secondary use. Data is available for general research use according to the following requirements for data access and data attribution (https://adknowledgeportal.synapse.org/Data%20Access).

Data obtained from the Gene Expression Omnibus. snRNA-seq of human brain vasculature was obtained under accession GSE163577^22^ and scRNA-seq of cerebrospinal fluid immune cells under accession GSE200164^20^, both openly available without restriction at https://www.ncbi.nlm.nih.gov/geo/.

Data obtained from published supplementary material. The following processed datasets were obtained from the supplementary information of their original publications, which is openly available from the publisher in each case: neuropathology-associated and case-control differential splicing results (Supplementary Tables 2 and 8 of Raj et al.^39^); m6A RNA methylation changes (Supplementary Table 24 of Castro-Hernández et al.^40^); plasma proteomic profiles across age (Supplementary Table 16 of Lehallier et al.^17^); plasma protein age associations (Supplementary Table 25 of Oh et al.^18^); brain phosphoproteomics (Supplementary Table 2 of Morshed et al.^12^); brain glycoproteomics (Supplementary Table 2 of Zhang et al., 2020^13^ and Zhang et al., 2024^14^); brain and plasma metabolomics (supplementary material of Kalecky et al.^46^); tau interactomes (Supplementary Tables 1, 3 and 5 of Tracy et al.^29^); PSEN1 and APP interactomes (Supplementary Table 1 of Hosp et al.^30^); γ-secretase substrates (Supplementary Table 1 of Hou et al.^28^); microRNA meta-analysis results of Takousis et al.^42^; CRISPRi screen results (Tian et al.^26^, Leng et al.^24^, Dräger et al.^23^, Parra Bravo et al.^27^, Samelson et al.^25^); and GWAS results (Kunkle et al.^31^, Wightman et al.^32^, Bellenguez et al.^34^).

Databases and knowledge resources. Alzheimer’s disease gene associations were retrieved from the Comparative Toxicogenomics Database (https://ctdbase.org, MeSH identifier D000544)^36^ and from DisGeNET (https://www.disgenet.org, disease identifier C0002395)^37^. GWAS results were retrieved from the NHGRI-EBI GWAS Catalog (https://www.ebi.ac.uk/gwas, MONDO identifier 0004975)^33^. MicroRNA target predictions were obtained from miRDB (https://mirdb.org)^44^ and TargetScan (https://www.targetscan.org)^45^. Clinical trial and drug-target information was obtained from DrugBank (https://go.drugbank.com)^49^, ClinicalTrials.gov (https://clinicaltrials.gov) and Alzforum (https://www.alzforum.org).

## Code Availability

The code for embedding generation and model training is available at https://github.com/Greta-B/BRIDGE-AD.

## Materials Availability

Plasmids generated in this study are available from the corresponding author (Y.S.) under a material transfer agreement.

## Supplementary table list

**Supplementary Table 1.** Curated Class 1 AD effectors and supporting evidence.

**Supplementary Table 2.** Primary cell-type annotations of curated Class 1 AD effectors.

**Supplementary Table 3.** AD gene-signature library and signature membership.

**Supplementary Table 4.** Genome-wide BRIDGE-AD effector prioritisation results.

**Supplementary Table 5.** Predicted Class 1 effectors nominated from the Class 0 gene set.

**Supplementary Table 6.** Self-organising map gene-cluster assignments.

**Supplementary Table 7.** Gene-level differential expression results for wild-type and *SCARB2*-overexpressing HMC3 microglia.

**Supplementary Table 8.** Diagnosis and available metadata for postmortem human brain tissues.

**Supplementary Table 9.** Antibodies used in this study.

**Supplementary Table 10.** DNA oligonucleotides used for cloning, mutagenesis and RT-qPCR.

## Methods

### Curation and annotation of Class 1 AD effectors

Class 1 AD effectors were curated from published experimental studies reporting genes whose perturbation altered AD-relevant phenotypes, with a focus on studies published within the past 10 years. Relevant studies were identified through a combination of programmatic PubMed screening using the RISmed package and manual literature review. Candidate genes were selected when experimental evidence linked their perturbation to measurable AD-relevant outcomes, including neuropathology, cognitive phenotypes, or molecular pathology. Class 1 genes were annotated for directionality as protective, detrimental, or mixed. Protective effectors were defined as genes whose loss-of-function worsened, and/or gain-of-function improved, AD-relevant phenotypes; detrimental effectors showed the opposite pattern. For each gene, we used the publication(s) supporting its Class 1 nomination to annotate the primary cell type, in vitro and in vivo evidence, as well as neuropathological and behavioural outcomes. Class 1 effectors were nominated as GWAS candidates or therapeutic targets if they appeared in curated GWAS and clinical target lists, respectively. Eight canonical AD effectors – ADAM10, BACE1, APOE, MAPT, APP, PSEN1, PSEN2, and TREM2 – were included in the Class 1 set based on extensive AD-relevant literature but were not assigned to specific source studies. The final Class 1 set comprised 153 genes and is intended as a high-confidence reference set for the task of supervised binary classification rather than an exhaustive catalogue of all AD-associated genes.

### Gene Embedding Construction

Human PPI network data were obtained from the STRING (v12.0)^51^ database and filtered to retain interactions with a combined confidence score of ≥ 400. The largest connected component of the resulting network was retained, comprising 19,486 nodes. To construct the Alzheimer’s disease-specific gene feature matrix, we defined signatures as gene sets containing at least two genes and quantified the network proximity of each PPI network node to each of the 317 curated AD gene signatures that met this condition. Shortest path lengths were calculated using the NetworkX package in Python^77^. Let *G = (V,E)* denote the PPI network, where *V* is the set of nodes and *E* is the set of edges. Let *S*⊆*V* denote a gene signature and *X*∈*V* the gene/protein for which network proximity features are being calculated. We define *d(s,X)* as the shortest path length between nodes *s* and *X*, and *N(X)* as the set of direct neighbours of *X* in the network. Then, for each node *X* and each signature *S*, we calculate three proximity metrics:

Average shortest path distance (ASP) – measures the average network distance from *X* to all genes in a signature:

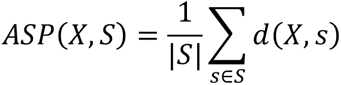

Minimum shortest path distance (MSP) – measures the distance from *X* to the closest gene in the signature:

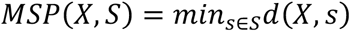

Ratio of neighbours (RN) – measures the fraction of the direct network neighbours of *X* that belong to the signature *S*:

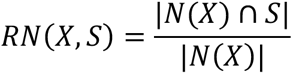

Applying these three proximity metrics across the 317 signatures produced an embedding matrix of 19,486 genes by 951 features.

To assess whether the observed proximity between *X* and *S* was greater or smaller than expected by chance, each feature was normalised using a degree-matched null distribution. For each metric *m∈ {*ASP, MSP, RN*}*, we generated an empirical null distribution by recalculating *m(X,S)* using randomly sampled source nodes matched to the degree of *X* and randomly sampled target signatures matched to the size and degree composition of *S*. This randomisation procedure was repeated 1,000 times. The mean *μ_m,null_* and standard deviation *σ_m,null_* of the resulting null distribution were then used to convert each observed metric value to a z-score:

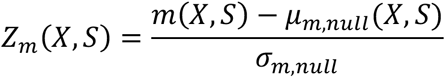

Because the human PPI network is scale-free^78^, a small number of nodes have very high degree centrality. To improve random sampling diversity while preserving degree structure, nodes were grouped into degree-based bins, with adjacent bins merged until each contained at least 100 nodes. Random source and target nodes were then sampled from the corresponding degree bins.

### ML framework

The constructed gene-feature space was used to train supervised classifiers to identify AD effector genes. The 153 curated AD effector genes constituted the positive class (Class 1) and were partitioned 80:20 into a training/cross-validation set (n = 122) and an independent test set (n = 31) reserved for final model evaluation. The remaining genes in the feature space formed the negative class (Class 0; n = 19,333) and were divided in the same proportion into training (n = 15,372) and test (n = 3,961) sets. Here, Class 0 labelling rests on the assumption that most genes are not involved in AD pathogenesis, such that treating the unlabelled pool as negatives introduces few false negatives into the training data. To mitigate class imbalance and to reduce variance through model averaging, the 15,372 Class 0 training genes were partitioned into 126 non-overlapping subsets of 122 genes, matching the number of Class 1 training genes. Each balanced training set therefore comprised the same 122 Class 1 genes together with a distinct subset of 122 Class 0 genes, ensuring that every negative gene was used exactly once across the 126 sets.

Four base classifiers (Logistic Regression (LR), Support Vector Machine (SVM), Random Forest (RF) and Gradient Boosting (GB)) together with a soft voting ensemble that averaged the predicted Class 1 probabilities of the four were trained independently on each of the 126 balanced training sets. For LR and SVM, feature standardisation (zero mean, unit variance) was implemented in the model pipeline and fitted within each cross-validation fold, while the tree-based RF and GB classifiers were trained on unscaled features. Hyperparameters were tuned separately within each balanced training set by optimising the AUROC under stratified 5-fold cross-validation. For LR, grid search was applied over the regularisation penalty (L1, L2, elastic net), regularisation strength C, and compatible solvers (liblinear, saga). For SVM, tuning spanned linear and radial basis function kernels and their associated C and γ parameters. For RF and GB, a two-stage strategy was used: an initial grid search over the principal hyperparameters (RF: number of trees, bootstrap sample fraction, and features per split; GB: number of trees, learning rate, subsample fraction, and features per split), followed by a randomised search over tree complexity and regularisation parameters (maximum depth, minimum samples per split and per leaf, maximum leaf nodes, minimum impurity decrease, and, for RF, cost-complexity pruning). Model performance was assessed by stratified 5-fold cross-validation within each balanced training set using accuracy, precision, recall, F1 score, AUROC and AUPRC. Accuracy was defined as the proportion of all genes classified correctly; precision as the fraction of genes predicted as Class 1 that were true Class 1 genes; recall as the fraction of true Class 1 genes recovered; and the F1 score as the harmonic mean of precision and recall. AUROC summarised discrimination across classification thresholds by comparing true-positive and false-positive rates, whereas AUPRC summarised the precision–recall trade-off.

To assess whether reducing redundancy among the network-derived features improved classification performance, correlation-based pruning was applied to the 951-feature matrix prior to model training. Pruning was carried out at five thresholds (|ρ| ≥ 0.95, 0.90, 0.85, 0.80 and 0.75) alongside the unpruned gene embedding. At each threshold, pairwise Spearman correlations were computed between all features, and a redundancy graph was built with features as nodes and edges joining pairs above the cut-off. Connected components of this graph defined redundancy clusters, each represented by its medoid (the feature with the highest mean absolute correlation to the other members). Singleton features belonging to no cluster were retained in full. Each resulting matrix was then supplied to the identical supervised learning pipeline described above, allowing performance to be compared directly across pruning thresholds and against the unpruned baseline.

### Model evaluation and final prediction

Cross-validation identified the soft-voting ensemble as the best-performing classifier. Feature pruning did not improve performance; therefore, the most conservative pruning threshold (|ρ| ≥ 0.95) was selected to reduce redundancy while preserving baseline performance, and this configuration was carried forward for evaluation on the held-out test data. Following the training procedure, balanced test sets were constructed by partitioning the 3,961 held-out Class 0 genes into groups of 31 – matching the number of held-out Class 1 genes – and pairing each group with the same 31 Class 1 test genes. Each of the 126 ensemble models was evaluated on every balanced test set using accuracy, precision, recall, F1 score, AUROC and AUPRC, and the resulting metrics were averaged across all models and test sets. To evaluate performance on the full imbalanced hold-out set, a consensus score was generated for each held-out gene by averaging its predicted Class 1 probability across the 126 ensemble models. AUPRC was then computed from these consensus scores over the complete hold-out set (31 positive and 3,961 negative genes).

Feature importance on the held-out test data was quantified using SHAP^53^, applied to the ensemble classifier’s predicted Class 1 probability. For each of the 126 balanced ensemble models, a background reference was built by summarising its balanced training set into 50 k-means centroids, and SHAP values were estimated for every held-out gene across all balanced test sets using KernelExplainer with nsamples = 300. SHAP values were averaged across the ensemble models and test sets, followed by feature ranking by mean absolute SHAP value.

To obtain final genome-wide scores, the framework was retrained on the full gene pool (153 Class 1 genes and 19,333 Class 0 genes partitioned across balanced training sets). Class 1 probabilities were then aggregated using an out-of-training average, such that a gene’s Class 1 score was derived only from models that did not use it during training.

### Literature co-mention analysis

To assess literature attention, PubMed co-mentions between each Class 1 or top-prioritised gene and AD were quantified using the RISmed package. Gene identifiers were first mapped to Entrez IDs to retrieve corresponding aliases, after which ambiguous aliases producing non-specific matches were manually excluded. AD-related search terms included “Alzheimer’s disease” and “dementia”, with searches restricted to article titles and abstracts.

### Benchmarking

The BioConcept^1^, genePT^2^, Mashup^3^, scGPT^4^, Geneformer^5^, and FRoGS-ARCHS4^6^ embeddings were sourced from their respective repositories; the protT5^7^ embedding was retrieved from the STRING database. Together with the BRIDGE-AD embedding, all representations were pre-processed by mapping gene identifiers to Entrez IDs and identifying the set of genes shared across every embedding (13,386 genes; Class 1 = 131, Class 0 = 13,255). Each embedding was restricted to this common gene set, and a single train/test split was applied across all of them. The training data (Class 1 = 105; Class 0 = 10,605) were partitioned into 101 balanced training sets using the same gene assignments for every embedding, and each embedding was then trained under the identical classification framework described above. Performance was evaluated on the held-out test data (Class 1 = 26, Class 0 = 2,650) using 102 test sets. In addition, consensus predictions were generated for every held-out gene and AUPRC was computed across the full imbalanced test set.

To assess how well each model recovered AD effector genes identified by independent studies, five external sets were sourced from AGORA^9^, NIAGADS^10^, Jaladanki et al. (2021)^54^, Bassil et al. (2021)^55^, and Liu et al. (2025)^56^. Each set was restricted to genes present in the 13,386 shared genes and filtered to remove the Class 1 genes used in training, ensuring an unbiased evaluation. For each embedding, the framework was retrained on the full common gene pool (13,386 genes) and every gene was scored by an out-of-training average across the balanced models, so that a gene’s Class 1 probability derived only from models that had not been trained on it. Recovery was quantified as AUPRC for each external set, treating that set’s genes as positives and all remaining Class 0 genes as negatives.

### Self-organising map (SOM) training and gene module identification

For each of the 251 genes, direct interactors were extracted from the STRING PPI network^51^ (confidence score ≥ 400) to define a gene neighbourhood. Each neighbourhood was tested for Reactome^57^ pathway enrichment, considering only the terminal (leaf) nodes of the Reactome hierarchy. Pathways enriched in at least three neighbourhoods were retained (g:SCS-adjusted p < 0.05; 877 pathways). For every neighbourhood-pathway pair, the resulting precision and recall values were combined into an F1 score, yielding an 877 × 251 pathway-by-gene matrix. A self-organising map (SOM Toolbox for MATLAB^79^) was trained on this matrix using a 14 × 14 hexagonal grid (196 nodes), initialised randomly and trained by batch learning for 500 epochs. Training embedded each gene as a 196-dimensional profile of activations over the node lattice, with topologically adjacent nodes capturing related pathway signatures. The 251 genes were subsequently clustered by applying k-means to these activation profiles, with the optimal number of clusters selected by maximising mean silhouette width. Final gene cluster assignments were obtained by consensus clustering: a gene-by-gene co-association matrix, recording the fraction of 1,000 k-means runs in which each pair co-clustered, was partitioned by spectral clustering at k = 19. For visualisation, the gene activation profiles were projected onto a UMAP. Within each gene cluster, two-dimensional kernel density estimates were computed, and density contours corresponding to the 5th, 25th, 50th, and 75th density quantiles were overlaid to indicate cluster extent and core density. Independently of the gene-level clustering, the 196 SOM nodes were also clustered by k-means to characterise pathway organisation across the map. The optimal number of node clusters was selected by maximising the mean silhouette width (k = 21). Each node cluster was annotated according to the dominant Reactome top-level pathway category among the pathways mapping to its nodes, and clusters with the same dominant category were merged, yielding 20 final pathway regions.

### AlphaFold modeling of SCARB2-GCase dimers

Structural models of SCARB2-GCase dimers were generated using AlphaFold 3^80^ to assess how perturbation of SCARB2 or GCase glycosylation could affect predicted complex formation. Protein sequences for human SCARB2 (UniProt ID: Q14108-1) and human GCase/GBA1 (UniProt ID: P04062-1) were obtained from UniProt. An experimentally resolved SCARB2-GCase luminal domain structure was obtained from the Protein Data Bank (PDB ID: 9FJF^74^) and used as a structural reference. As SCARB2 and GCase contain nine and four N-glycosylation sites, respectively, we generated 14 SCARB2-GCase models: one fully glycosylated reference model, in which all SCARB2 and GCase glycosylation sites were occupied, and 13 single-site perturbation models, in which one SCARB2 or GCase glycan was omitted at a time. With the exception of the SCARB2 N325 glycan, all SCARB2 and GCase glycans were modelled as the conserved Man₃GlcNAc₂ N-glycan core to represent experimentally supported N-glycosylation occupancy and proximal steric effects while avoiding assumptions about heterogeneous terminal glycan processing. The N325 SCARB2 glycan was modelled as an extended high-mannose glycan, Man₆GlcNAc₂, based on the glycan structure reported by Zhao et al., 2014^81^. For each glycosylation configuration, three AlphaFold 3 models were generated using random seeds 1, 2 and 3. Predicted SCARB2-GCase interface confidence was assessed using the interface predicted template modelling score (ipTM), and ipTM values were summarised across seeds as mean ± SD for each model configuration.

### Human postmortem brain tissues

Postmortem human brain tissues were obtained from the Banner Sun Health Research Institute. Donor diagnosis and other metadata are reported in **Supplementary Table 8**.

### Maintenance of HEK293T and HMC3 cell lines

Human embryonic kidney 293T (HEK293T) and human fetal microglia (HMC3) cell lines were cultured in DMEM (Corning, 15-013-CV) supplemented with 10% fetal bovine serum (Sigma-Aldrich, F4135), 1% GlutaMAX^TM^ (Gibco, 35050079), and 1% Antibiotic-Antimycotic supplement (Gibco, 15240112). The cells were maintained in a humidified 5% CO_2_ incubator at 37 °C and were passaged using Accutase (Sigma-Aldrich, A6964) once they reached 80% confluency. Cells were routinely tested for mycoplasma contamination using the MycoAlert^TM^ Mycoplasma Detection Kit (Lonza, LT07-318).

### Cloning of *SCARB2* and *GBA1* expression vectors

Human SCARB2 (Human, LIMPII/SR-B2 cDNA ORF Clone in Cloning Vector, HG11063-M) and GBA1 (Human, GBA/glucocerebrosidase cDNA ORF Clone in Cloning Vector, HG12038-G) cDNA cloning vectors were purchased from Sino Biological. SCARB2 and GBA1 coding regions were amplified by PCR with primers containing NotI and EcoRI restriction enzyme cutting sites. pLVX-UbC-rtTA (Addgene, 127289) PCR amplicons were purified with the QIAquick PCR & Gel Cleanup Kit (Qiagen, 28506), digested with NotI-HF (NEB, R3189) and EcoRI-HF (NEB, R3101L) restriction enzymes, and ligated into NotI/EcoRI digested and CIP (NEB, M0290) dephosphorylated pLVX-UbC-rtTA backbone using T4 DNA ligase (NEB, M0202) at 16°C for 12 hours. Ligated plasmids were propagated in DH5α *E. coli* bacteria and extracted using the E.Z.N.A.^®^ Endo-free Plasmid DNA Mini Kit II (Omega Bio-tek, D6950).

Coding regions for SCARB2-N45Q, SCARB2-N68Q, SCARB2-N105Q, SCARB2-N206Q, SCARB2-N224Q, SCARB2-N249Q, SCARB2-N304Q, SCARB2-N325Q, SCARB2-N412Q, GBA1-N58Q, GBA1-N98Q, GBA1-N185Q, and GBA1-N309Q isoforms were generated by PCR directed mutagenesis and cloned into the pLVX-UbC-rtTA vector as described above.

FLAG-SCARB2 and GBA1-FLAG vectors were PCR amplified from wild-type SCARB2 and GBA1 vectors, respectively, using primers containing 3xFLAG and flexible linker sequences. PCR amplicons were cloned into the pLVX-UbC-rtTA vector as described above.

All plasmids generated in this study were verified by whole-plasmid sequencing using Nanopore sequencing (Plasmidsaurus). Oligonucleotides used for cloning and mutagenesis are listed in **Supplementary Table 10**.

### Production of lentiviral particles

Lentiviral particles were produced in HEK293T cells using either a PEI-based or a calcium phosphate-based transfection workflow adapted from our previously described protocol. Briefly, HEK293T cells were seeded in 10-cm dishes 18 hours before transfection at 1:4 splitting ratio. The cells were transfected with the transfer plasmid and second-generation packaging plasmids psPAX2 (Addgene, 12260) and pMD2.G (Addgene, 12259), and the medium was replaced 6 hours after transfection. Virus-containing supernatant was collected 72 hours after transfection, clarified by low-speed centrifugation (600 g for 10 minutes) and filtered through a 0.45 µm filter. Subsequently, lentiviral particles were concentrated by ultracentrifugation (40,000 g for 2 hours at 4 °C) and resuspended in DPBS. Concentrated virus was stored at –80°C in 10 µL aliquots.

### Generation of stable overexpression lines

SCARB2 and GBA1 overexpression lines were generated by lentiviral transduction of doxycycline-inducible expression vectors. Target cells were seeded in 12-well plates at 20% confluence and transduced in the presence of 4 µg/mL polybrene and a survival-enhancing cocktail adapted from Chen et al.^82^ (10 µM ROCK inhibitor Y-27632 [Cellagen Tech, C9127], 5 μM emricasan [MedChemExpress, HY-10396], 0.7 μM trans-ISRIB [MedChemExpress, HY-12495], and 1X polyamine supplement [Sigma-Aldrich, P8483]). The medium was exchanged 24 hours after transduction, and 1 µg/mL puromycin was added 72 hours after transduction for 5 days.

Transgene expression was induced with a 1 µg/mL doxycycline treatment for at least 3 days before downstream experiments were performed. Overexpression was validated by RT-qPCR as the primary readout as well as immunoblotting where indicated.

### RNA isolation, reverse transcription, and qPCR

Total RNA was isolated using either a Quick-RNA Miniprep Kit (Zymo Research, R1054) or the TRIzol™ Reagent (Thermo Fisher Scientific, 15596018). RNA concentration was assessed by spectrophotometry. For cDNA synthesis, 3 µg of total RNA was reverse-transcribed using the Tetro cDNA synthesis kit (BioLINE, Bio-65043) following the manufacturer’s protocol. qPCR was performed using the iTaq™ Universal SYBR® Green Supermix (Bio-Rad, 1725124) or SYBR Green Master Mix/DyNAmo Flash SYBR Green (Thermo Scientific, F416/F416L) with three biological and three technical replicates per experiment. Relative gene expression was calculated using the ΔΔCt method with GAPDH as the endogenous reference gene. RT-qPCR primer sequences are provided in **Supplementary Table 10**.

### Protein extraction, SDS-PAGE, and immunoblotting

Primary tissue and cell lysates were prepared by sonication on ice in the RIPA buffer supplemented immediately before use with protease inhibitors. Lysates were clarified by centrifugation at 10,000 g for 30 min at 4 °C, and supernatants were transferred to fresh tubes. Protein concentration was measured using the Pierce BCA Protein Assay Kit (Thermo Fisher Scientific, 23227) according to the manufacturer’s instructions. 5-20 µg of total protein was combined with reducing Laemmli sample buffer, denatured at 95 °C for 5 min, and separated by SDS-PAGE using an 8-12% polyacrylamide gel.

Proteins were transferred to a PVDF membrane using semi-dry transfer conditions (5 V for 30 min). The membrane was blocked in 5% non-fat milk or 5% BSA in TBST for 1 hour at room temperature and incubated with primary antibodies diluted in TBST and incubated at 4 °C overnight. The membrane was washed with TBST 3 times for 10 minutes each and incubated with HRP-conjugated secondary antibodies for 2 hours at room temperature. After incubation, the membrane was washed with TBST 3 times for 10 minutes each and incubated with the SuperSignal West Pico PLUS Chemiluminescent Substrate (Thermo Fisher Scientific, 34577) for 1 min to develop a chemiluminescent signal. The signal was detected using the ChemiDoc Imaging System (Bio-Rad). Band intensities were quantified using Bio-Rad Image Lab and normalised to GAPDH. Antibody sources, catalogue numbers, applications and dilutions are provided in **Supplementary Table 9**.

### Glycan characterisation by Endo H and PNGase F digestion

N-linked glycosylation and glycoprotein maturation of SCARB2 and GCase glycoforms were assessed using Endoglycosidase H (Endo H, Thermo Fisher Scientific, 50-811-826) and PNGase F (Thermo Fisher Scientific, 50-591-168) digestion. Total protein was isolated as described earlier. 5-20 µg of protein from each condition was aliquoted into control, Endo H-treated, and PNGase F-treated reactions. For denaturing digestion conditions, samples were brought to 10 µL with 1× glycoprotein denaturing buffer and heated at 100 °C for 10 min to denature glycoproteins. Endo H reactions were then adjusted to a 20 µL final volume with 1× GlycoBuffer 3 and 0.5 µL Endo H and incubated at 37 °C for 3 h. PNGase F reactions were adjusted to a 20 µL final volume with 1× GlycoBuffer 2, NP-40, and 0.5 µL PNGase F and incubated at 37 °C for 3 h. After digestion, reactions were mixed with reducing Laemmli sample buffer and analyzed by SDS-PAGE and immunoblotting, with deglycosylation assessed by electrophoretic mobility shifts of SCARB2 or GCase glycoforms.

### Immunostaining

Cells were fixed in 4% paraformaldehyde (Millipore Sigma, 158127) for 15 min, washed with DPBS (Thermo Fisher Scientific, 14190144). Cells were then permeabilised in DPBS containing 0.5% Triton X-100 (MilliporeSigma, X100) for 15 min and blocked in DPBS containing 5% donkey serum (Thermo Fisher Scientific, 16210064) for 1 h at room temperature. Cells were incubated with primary antibodies at 4°C overnight. The primary antibodies were diluted in DPBS with 5% donkey serum at 1:300 dilution. The next day, cells were washed with DPBS and incubated with secondary antibodies for 1 h at room temperature. Secondary antibodies were diluted in DPBS at 1:500 dilution. After incubation, cells were washed with DPBS, incubated with DAPI (Thermo Fisher Scientific, 62248) at 1:10,000 dilution for 15 min, and mounted with Fluoromount-G reagent (SouthernBiotech, 0100-01).

### Immunofluorescence image acquisition and quantification

Immunolabeled cells were imaged using Nikon Ti-2, and the images were analyzed by Fiji/ImageJ. Images were acquired as single optical sections.

### Lysosome labelling in live cells

Acidic intracellular compartments in live HMC3 microglia were visualized using LysoTracker Green DND-26 (Thermo Fisher Scientific, L7526) following the manufacturer’s protocol. Briefly, wild-type and SCARB2-overexpressing HMC3 cells were plated on a glass-bottom 96-well plate at 10,000 cells/well density. As a lysosomal deacidification control, wild-type HMC3 cells were treated with 100 nM bafilomycin A1 (Cell Signaling Technology, 54645S) for 24 hours before LysoTracker labelling. Cells were incubated with LysoTracker diluted in pre-warmed culture medium at 37 °C for 45 min protected from light. After labelling, cells were washed with pre-warmed Hank’s balanced salt solution (HBSS) and imaged immediately using a Nikon Ti-2 microscope.

### Dual-colour LC3 reporter autophagic flux assay

Autophagic flux was measured using an HMC3 line carrying a dual-colour LC3 reporter generated using the pMRX-IP-GFP-LC3-RFP-LC3ΔG construct (Addgene, 84572). The reporter consists of autophagy-sensitive GFP-LC3 and a non-lipidated cytoplasmic RFP-LC3ΔG control. In this system, GFP-LC3 enters the autophagic pathway and is degraded during autophagic flux, whereas RFP-LC3ΔG remains cytosolic and provides an internal normalisation signal. Thus, a lower GFP/RFP ratio indicates increased autophagic flux, while an increased ratio indicates impaired autophagic degradation.

Wild-type and SCARB2-overexpressing HMC3 reporter cells were plated on glass-bottom 96-well plates at 10,000 cells per well. Cells were treated with 100 nM bafilomycin A1 as a lysosomal deacidification control or 250 nM Torin 1 as an autophagy-activation control for 24 h. Imaging was performed using identical GFP and RFP/TRITC acquisition settings across conditions. Fluorescence intensity was quantified using a Fiji macro. For each image, a duplicate segmentation image was converted to 8-bit, thresholded using a fixed intensity range of 10–252, converted to a binary mask, and separated by watershed. Cellular regions-of-interest (ROIs) were identified using particle analysis with a minimum size threshold of 5,000 pixels². Mean fluorescence intensity was then measured from the corresponding original-bit-depth measurement image for each ROI. ROI-level mean intensities were averaged per image, and GFP/RFP ratios were calculated by dividing the mean GFP intensity by the corresponding mean RFP/TRITC intensity.

### LPS/IFNγ inflammatory stimulation

Inflammatory responsiveness of HMC3 microglia was assessed by stimulation with 100 ng/mL lipopolysaccharide (LPS, Thermo Fisher Scientific, 00-4976-03) and 20 ng/mL recombinant human IFNγ (Cell Signaling Technology, 8927SC). Unstimulated cells were included as matched controls. 24 hours after stimulation, cells were harvested for RNA extraction and RT-qPCR analysis.

### RNA sequencing of WT and SCARB2-overexpressing HMC3 microglia

Total RNA was isolated using the Quick-RNA Miniprep Kit (Zymo Research, R1054) including on-column DNase treatment. RNA concentration and purity were measured by spectrophotometry. For each sample, 30 μL of purified total RNA was submitted to Plasmidsaurus at 10-100 ng/μL, corresponding to a minimum input of 300 ng total RNA per sample. RNA-seq libraries were generated using a stranded 3’-end-counting approach. In brief, mRNA was converted to cDNA using a poly(dT)VN primer, followed by second-strand synthesis, tagmentation, indexing, and amplification. Libraries were sequenced on Illumina NovaSeq instruments using single-end sequencing. Reads were generated from the 3’-end of transcripts, with an approximate read length of 90 bp, and unique molecular identifiers were included for deduplication. RNA-seq data were filtered to remove lowly expressed genes using the filterByExpr function^83^, normalised using the trimmed mean of M values method, and transformed using voom from the limma package^84^. Differential expression between WT and SCARB2-OE HMC3 microglia was assessed using the lmFit and eBayes functions from limma, with batch included as a covariate.

### Statistical analysis

Statistical analyses were performed using GraphPad Prism version 10. For cell-culture experiments, biological replicates were independent cultures, wells, or independently prepared cell batches as indicated; technical replicates were repeated measurements from the same biological sample. For human tissue analyses, each donor was treated as an independent biological replicate. Data are presented as mean ± SD. For two-group comparisons, a two-sided unpaired Welch’s t-test was used. For multiple-group comparisons, one-way ANOVA followed by Tukey was used. For correlation analyses, Spearman correlation was used as specified in the figure legend. Differences in variance between diagnostic groups were assessed using the Fligner–Killeen test. Omics analyses used Benjamini-Hochberg false-discovery-rate correction unless otherwise stated. Significance notation was defined as ns, not significant; *P < 0.05; **P < 0.01; ***P < 0.001; and ****P < 0.0001.

## Supplementary methods

### Brain bulk RNA-seq

Bulk RNA-seq count data from ROSMAP donors^41^ were downloaded from Synapse (syn12104384). Samples were excluded if they had missing RNA integrity number (RIN) values or RIN < 5, missing post-mortem interval (PMI) values or PMI > 36 hours, or missing age at death information. Donors were then stratified into healthy control and AD groups using stringent neuropathological and cognitive criteria to compare molecular profiles between cognitively healthy individuals and late-stage AD cases. The healthy cohort included participants with Braak scores of 0–3, CERAD scores of 3–4, cogdx score of 1, and MMSE scores ≥ 27. The AD cohort included participants with Braak scores of 4-6, CERAD scores of 1–2, cogdx scores of 4–5, and MMSE scores ≤ 14. The final dataset comprised 72 healthy control and 70 AD ROSMAP samples. Count data were filtered to remove genes with expression below 20 counts per million using the *filterByExpr* function^83^. Data were then normalised using the trimmed mean of M values (TMM) method and transformed using *voom* from the limma package^84^. Differential expression between AD and healthy control samples was performed using the *lmFit* and *eBayes* functions from limma, controlling for age at death, sex, PMI, RIN, and batch. Differentially expressed genes were defined as those with |log_2_FC| > 0.50 and FDR < 0.05.

### Brain bulk miRNA array

The ROSMAP miRNA dataset^43^ was downloaded from Synapse (syn3387325). The dataset contained 309 miRNAs and had already undergone initial preprocessing, including expression filtering, normalisation and batch correction. We then excluded samples if they had missing RIN values or RIN < 5, missing PMI values or PMI > 36 hours, or missing age at death information. Remaining donors were stratified into healthy control and AD groups using stringent neuropathological and cognitive criteria to compare molecular profiles between cognitively healthy individuals and late-stage AD cases. The healthy cohort included participants with Braak scores of 0–3, CERAD scores of 3–4, cogdx score of 1, and MMSE scores ≥ 27. The AD cohort included participants with Braak scores of 4–6, CERAD scores of 1–2, cogdx scores of 4–5, and MMSE scores ≤ 14. The final dataset comprised 71 healthy control and 60 AD ROSMAP samples. Differential expression between AD and healthy control samples was performed using the *lmFit* and *eBayes* functions from limma^84^, controlling for age at death, sex, PMI, and RIN. Differentially expressed miRNAs were defined as those with FDR < 0.05. For each differentially expressed miRNA, putative target genes were identified by intersecting predictions from miRDB^44^ and TargetScan^45^ databases. miRDB targets were retained using a target score > 70, while TargetScan targets were retained using a cumulative weighted context score < –0.2. When the mature miRNA strand was not specified in the array annotation, targets for both the 5p and 3p mature miRNA forms were considered separately. For miR-504-5p, only miRDB-predicted targets were used, as this miRNA was not available in TargetScan.

### Other miRNAs

We additionally included eight miRNAs reported as differentially expressed in AD brain tissue compared with healthy controls in the meta-analysis by Takousis et al., 2019^42^: miR-125b-5p, miR-501-3p, miR-138-5p, miR-195-5p, miR-181c-5p, miR-454-3p, miR-455-5p, and miR-29c-3p. These miRNAs were selected based on the “strong” evidence classification defined by Takousis et al. As above, putative target genes were identified by intersecting predictions from miRDB^44^ and TargetScan^45^. miRDB targets were retained using a target score > 70, while TargetScan targets were retained using a cumulative weighted context score < –0.2. For miR-29c-3p, only miRDB-predicted targets were used, as this miRNA was not available in TargetScan.

### RNA splicing

The RNA splicing dataset comprising 450 subjects of ROSMAP and MSBB was reported by Raj et al. (2018)^39^. Processed neuropathology-associated differential splicing results were obtained from Supplementary Table 2 and used to generate two signatures: significantly upregulated introns (FDR < 0.05, beta > 0) associated with neuritic plaques, amyloid and tangles, and significantly downregulated introns (FDR < 0.05, beta < 0) associated with the same neuropathological measures. Separately, an additional gene list was generated from introns reported as differentially spliced in AD patients compared with healthy controls in Supplementary Table 8 of the original study.

### RNA methylation

Fully processed RNA m^6^A methylation change results from 6 AD patients and 6 healthy controls were downloaded from Supplementary Table 24 of Castro-Hernández et al. (2023)^40^. Three signatures were generated based on mRNAs showing increased, decreased or mixed m6A level changes in AD patients compared with healthy controls.

### Brain proteomics (Emory)

The Emory dataset consisted of two cohorts reported by Ping et al. (2018)^38^ and Higginbotham et al. (2020)^15^, which were retrieved from Synapse (syn20824864). The first cohort comprised 40 samples, including 10 controls, 10 AD cases, 10 Parkinson’s disease (PD) cases, and 10 AD–PD cases. The dataset also included 10 global internal standard (GIS) samples, with proteomic profiling performed using tandem mass tag (TMT) quantification. Batch correction was applied to normalised TMT abundance data using the TAMPOR^85^ methodology. Data were then log_2_-transformed, and missing values were imputed using the K-nearest neighbours (KNN) method implemented in DreamAI^86^. Differential abundance analysis between AD and control samples was performed using the *lmFit* and *eBayes* functions from the limma package^84^, controlling for age at death, sex and PMI. Differentially abundant proteins were defined using |log_2_FC| > 0.25 and FDR < 0.05. The second cohort comprised 27 samples, including 10 controls, 9 AD cases and 8 asymptomatic AD (ASYM) cases. The dataset also included 6 GIS samples, with proteomic profiling performed using TMT quantification. Batch correction and missing value imputation were performed as described for the first cohort. After these preprocessing steps, one ASYM sample was excluded because age at death information was unavailable. Differential abundance analysis between AD and control samples was performed as described for the first cohort.

### CSF proteomics

The CSF proteomics dataset comprising 150 AD patients and 147 controls was reported by Johnson et al. (2020)^11^ and downloaded from Synapse (syn21441787). The dataset also included 41 GIS samples, with proteomic profiling performed using TMT quantification. Batch correction was applied on normalised TMT abundance data using the TAMPOR^85^ methodology, after which data were log_2_-transformed and missing values were imputed using the KNN method implemented in DreamAI^86^. Differential abundance analysis between AD and control samples was performed using the *lmFit* and *eBayes* functions from the limma package^84^, controlling for age and sex. Differentially abundant proteins were defined using |log_2_FC| > 0.20 and FDR < 0.05.

This dataset also contained quantitative AD biomarker measurements, including pTau and Aβ42, enabling modelling of disease-associated protein level abundance trajectories using generalised additive models (GAMs)^87^. Two samples were excluded before trajectory modelling because of biomarker-level quality issues: one had missing data and the other had outlier biomarker measurements. The imputed data were scaled and centred, and, for each protein, abundance was modelled as a function of the pTau/Aβ_42_ ratio, age and sex. The pTau/Aβ_42_ ratio and age were included as smooth terms using penalised regression splines, while sex was included as a categorical covariate. Models were fitted using a scaled-t family distribution with a gamma value of 1.5. The significance of the pTau/Aβ_42_ smooth term was extracted for each protein, and p-values were adjusted using the Benjamini-Hochberg method. Proteins with FDR < 0.05 were considered significantly associated with disease. To group proteins with similar disease-associated trajectories, cosine similarity was calculated between predicted trajectories for all significant proteins. The resulting similarity matrix was converted to a dissimilarity matrix and used for hierarchical clustering with the Ward.D method. Silhouette analysis supported an optimal solution of two clusters, yielding two signatures.

### Brain proteomics across aging

The brain proteomics dataset from 84 healthy donors aged 30–68 in a Johns Hopkins cohort, reported by Johnson et al. (2020)^11^, was downloaded from Synapse (syn21441775). The dataset included 9 GIS samples, with proteomic profiling performed using label-free quantification (LFQ). Batch correction was applied using the TAMPOR^85^ methodology, as described previously^11^, after which data were log_2_-transformed and missing values were imputed using the KNN method implemented in DreamAI^86^. Samples with PMI >36 h were excluded, and the imputed data were scaled and centred. Age-associated protein abundance changes were modelled using GAMs^87^. For each protein, abundance was modelled as a function of age, PMI and sex, with age and PMI included as smooth terms using penalised regression splines and sex included as a categorical covariate. Models were fitted using a scaled-t family distribution with a gamma value of 1.25. The significance of the age smooth term was extracted for each protein, and p-values were adjusted using the Benjamini-Hochberg method. Proteins with FDR < 0.05 were considered significantly associated with age. To group proteins with similar age-associated trajectories, cosine similarity was calculated between predicted trajectories for all significant proteins. The resulting similarity matrix was converted to a dissimilarity matrix and used for hierarchical clustering with the Ward.D method. Silhouette analysis supported an optimal solution of two clusters, yielding two signatures.

### Plasma proteomics across aging 1

Plasma proteomic data from 171 healthy individuals aged 21–107, reported by Lehallier et al. (2019)^17^, were downloaded from Supplementary Table 16 of the original study. The dataset was generated using the SomaScan platform and comprised log_10_-transformed relative fluorescence unit (RFU) protein abundance values. Data were scaled and centred, and, for each protein, abundance was modelled as a function of age, sex, and cohort using GAMs^87^. Age was included as a smooth term using penalised regression splines, sex as a categorical covariate, and cohort as a random effect. Models were fitted using a scaled-t family distribution with a gamma value of 1.25. The significance of the age smooth term was extracted for each protein, and p-values were adjusted using the Benjamini-Hochberg method. Proteins with FDR < 0.05 were considered significantly associated with age. To group proteins with similar age-associated trajectories, cosine similarity was calculated between predicted trajectories for all significant proteins. The resulting similarity matrix was converted to a dissimilarity matrix and used for hierarchical clustering with the Ward.D method. Silhouette analysis supported an optimal solution of two clusters, yielding two signatures.

### Plasma proteomics across aging 2

The plasma proteomics dataset comprising 4,980 proteins measured across 5,676 individuals aged 19-104 was reported by Oh et al. (2023)^18^. Processed age association results, including the estimated slope of protein abundance change with age, were downloaded from Supplementary Table 25. Applying thresholds of Q value < 0.05 and absolute Age_beta > 0.0025 yielded two signatures, corresponding to plasma proteins that increased or decreased with age.

### Brain phosphoproteomics

The brain phosphoproteomics dataset of 25 AD patients and 23 non-demented controls was reported by Morshed et al., 2021^12^, and downloaded from Supplementary Table 2 of the original study. The data had already undergone initial pre-processing, including within-run normalisation of TMT quantification values using the CONSTrained STANdardization algorithm, followed by normalisation of the final relative TMT intensities to the non-demented, amyloid-negative control samples across TMT 10-plex analyses. Prior to differential abundance analysis, peptides supported by ≤ 3 PSMs were removed, and the data were further filtered to retain peptides detected in at least 50% of samples within each diagnostic group. The resulting intensities were log_2_-transformed, and non-phosphorylated and phospho-modified peptides were analysed separately. Differential abundance analysis was performed using msqrob2^88^, modelling diagnosis while adjusting for batch, PMI, sex and age at death. Proteins and phosphosites supported by multiple peptides were analysed by peptide-level robust mixed modelling with aggregation to the protein or phosphosite level (*msqrobAggregate*, with a sample-level random effect), whereas single-peptide features were analysed by robust fixed effect modelling (*msqrob*). For each layer, test statistics from the multi– and single-peptide models were pooled and a single Benjamini-Hochberg correction was applied across all features. For phosphosite-level analyses, both unadjusted phosphopeptide abundance and protein-adjusted phosphosite occupancy, calculated by subtracting the corresponding log_2_-transformed parent protein abundance from each phosphosite abundance, were assessed. Differentially abundant proteins and phosphosites, and differentially occupied phosphosites, were defined using |log_2_FC| > 0.25 and FDR < 0.05.

### Brain glycoproteomics

Protein glycosylation data from 8 AD and 8 control cases were reported by Zhang et al., 2020^13^, with an additional glycosylation dataset generated from the same donor cohort reported by Zhang et al., 2024^14^. Processed glycopeptide-level results, indicating whether each glycopeptide was detected in AD and/or control samples, were downloaded from Supplementary Tables 2 of the respective studies. Each dataset was processed separately by removing glycopeptides with identical detection status in AD and control samples, thereby retaining only glycopeptides detected exclusively in one condition. Based on these retained glycopeptides, proteins were assigned to one of three signatures per dataset: AD-associated increased glycosylation, comprising proteins with glycopeptides detected only in AD samples; AD-associated decreased glycosylation, comprising proteins with glycopeptides detected only in control samples; and mixed glycosylation, defined by the presence of both AD-specific and control-specific glycopeptides from the same protein.

### Metabolomics

Brain metabolomics data reported by Kalecky et al. (2022)^46^ were downloaded from the Supplementary Materials of the published study. A total of 136 samples were available from 35 AD subjects, 36 healthy controls, and 65 unrelated subjects. Initial preprocessing followed the approach described in the original study. Briefly, limits of detection (LODs) were calculated as the mean blank signal plus two standard deviations. Metabolites with more than 50% of values below the LOD in both AD and control samples were excluded. Zero or missing values were then replaced with LOD/2 values, as described in the Biocrates manual. To account for batch effects, metabolite values were normalised across plates using median scaling. The resulting data were log_2_-transformed and analysed using the ocEAn pipeline^75^. Metabolites were first mapped to KEGG identifiers, and metabolites sharing the same KEGG ID were averaged within each sample. Differential abundance analysis between AD and healthy control samples was performed using the *lmFit* and *eBayes* functions from the limma package^84^, controlling for age at death, sex, PMI, body mass index (BMI) and storage duration. The resulting metabolite t-statistics were used as input to the ocEAn framework, where the distance penalty was set to 8, the minimum branch length was set to 1 upstream and 1 downstream, and the ratio of upstream to downstream branch length for enzymes was left unbounded. Scores for reactions annotated as “reverse” were inverted, and multiple scores for the same enzyme across different reactions were averaged. Enzymes with an absolute normalised enrichment score >1.5 were used to generate signatures corresponding to increased or decreased inferred enzyme activity in AD.

Plasma metabolomics data reported by Kalecky et al. (2022) were also downloaded from the Supplementary Materials of the published study. The dataset comprised 158 samples, including 94 AD subjects and 64 healthy controls. The same analysis pipeline described for the cortex metabolomics data was applied to the plasma metabolomics dataset. During linear regression, sex, BMI, storage duration, blood collection site and fasting duration were included as covariates.

### Brain parenchyma snRNA-seq

A pre-processed and annotated snRNA-seq dataset from 48 ROSMAP participants^19^ was downloaded from Synapse (syn52383412). Donors were filtered and stratified into healthy and AD groups using stringent neuropathological and cognitive criteria to compare molecular profiles between cognitively healthy individuals and late-stage AD cases. The healthy cohort included participants with Braak scores of 0–3, CERAD scores of 3–4, cogdx score of 1, and MMSE scores ≥ 27. The AD cohort included participants with Braak scores of 4–6, CERAD scores of 1–2, cogdx scores of 4–5, and MMSE scores ≤ 14. The final dataset comprised 20 participants, including 11 healthy controls and 9 AD cases, and 560,760 nuclei from six brain regions: prefrontal cortex, entorhinal cortex, midtemporal cortex, hippocampus, angular gyrus, and thalamus. The snRNA-seq data were processed using Seurat^89^. Counts were log-normalised with a scale factor of 10,000, and the top 2,000 highly variable genes were used for principal component analysis. Region-associated batch effects were then corrected using Harmony^90^, followed by UMAP visualisation. Cell type-specific differential expression between AD and healthy controls was performed using the MAST method (*FindMarkers*), controlling for age at death, sex, percentage of mitochondrial genes, PMI, and brain region. DEGs were defined using |log_2_FC| > 0.25 and FDR < 0.05. Cell type-specific marker genes were identified using nuclei from healthy donors only, using the MAST method (*FindAllMarkers*) with the same covariates; markers were defined as genes with log_2_FC > 1.5 and FDR < 0.05.

To obtain additional insights into the brain snRNA-seq dataset and better understand the rewiring of molecular networks in AD, we also performed SCINET^47^ analysis. SCINET reconstructs cell type-specific protein-protein interaction networks by combining single-cell transcriptomic profiles with a reference interactome (PCNet^91^). Using SCINET workflow implemented in ACTIONet^92^, cells were grouped by disease state and cell type, and gene specificity scores were calculated for each group using the *compute.cluster.feature.specificity* function. These scores were then used to infer cell type– and disease state-specific interactomes with *run.SCINET.clusters*. Within each inferred network, weighted degree centrality was calculated for every gene using node strength from the igraph package^93^. To enable comparison across networks of differing sizes, centrality values were converted to within-network percentile ranks. Differential centrality between AD and healthy networks was assessed separately for each cell type, retaining cell types with ≥ 500 nuclei in both conditions. For each gene, the differential centrality score was calculated as the AD centrality percentile minus the healthy control centrality percentile. To assess the statistical significance of observed centrality changes, an empirical null distribution was constructed for each cell type by generating randomised networks using the *rewire* function in igraph, followed by recalculation of weighted degree centrality percentile ranks. This procedure was repeated to generate 10⁶ null centrality differences, against which the observed centrality differences were compared. The resulting p-values were adjusted for multiple testing using the Benjamini-Hochberg method. Genes with FDR<0.05 were used to define cell type-specific signatures of increased or decreased centrality in Alzheimer’s disease.

### Brain endothelium snRNA-seq

A pre-processed and annotated snRNA-seq dataset from 17 donors (9 AD patients and 8 healthy controls) was obtained from Yang et al. (2022)^22^ via GEO (GSE163577). The dataset included 143,793 nuclei enriched for vascular cells, generated using the vessel isolation and nuclei extraction for sequencing (VINE-seq) methodology described by the authors. Count data were log-normalised with a scale factor of 10,000, and the top 2,000 highly variable genes were used for principal component analysis using Seurat^89^. Batch effects were then corrected using Harmony^90^, followed by UMAP visualisation. Cell type-specific differential expression between AD and healthy controls was performed using the MAST method (*FindMarkers*), controlling for age at death, sex, percentage of mitochondrial genes, PMI, batch, and brain region. DEGs were defined using |log_2_FC| > 0.25 and FDR < 0.05. Cell type-specific marker genes were identified using nuclei from healthy donors only, using the MAST method (*FindAllMarkers*) with the same covariates; markers were defined as genes with log_2_FC > 1.5 and FDR < 0.05.

### CSF immune cell scRNA-seq

A pre-processed but unannotated scRNA-seq dataset of immune cells isolated from the cerebrospinal fluid of 59 donors was obtained from Piehl et al. (2022)^20^ via GEO (GSE200164). The cohort comprised 6 AD patients, 8 individuals with mild cognitive impairment (MCI), and 45 healthy controls. Count data were log-normalised using Seurat^89^ with a scale factor of 10,000. Principal component analysis was performed using the top 2,000 highly variable genes, followed by sample integration using Harmony^90^ and UMAP visualisation. Cell types were annotated using canonical marker genes. Pan-NK, myeloid, B-cell and T-cell clusters were then subsetted, and the clustering procedure was repeated to resolve more specific immune cell populations. Cell type-specific differential expression between AD patients and healthy controls aged >60 years was performed using the MAST method (*FindMarkers*), controlling for age, sex, and percentage of mitochondrial genes. DEGs were defined using |log_2_FC| > 0.25 and FDR < 0.05. To assess age-associated transcriptional changes, healthy donors were divided into two groups: < 70 years old (24 individuals) and ≥ 70 years old (21 individuals). Cell type-specific DEGs between these age groups were determined using the same MAST-based approach, controlling for sex and percentage of mitochondrial genes and applying the same significance thresholds. Cell type-specific marker genes were identified using cells from healthy donors only, using the MAST method (*FindAllMarkers*) while controlling for age, sex, and percentage of mitochondrial genes; markers were defined as genes with log_2_FC > 1.5 and FDR < 0.05.

### Brain snATAC-seq

A pre-processed and annotated dataset consisting of single-nucleus ATAC-seq (snATAC-seq) and snRNA-seq data^21^ from 92 ROSMAP participants was downloaded from Synapse (syn52293417). Donors were filtered and stratified into healthy and AD groups using stringent neuropathological and cognitive criteria to compare molecular profiles between cognitively healthy individuals and late-stage AD cases. The healthy cohort included participants with Braak scores of 0–3, CERAD scores of 3–4, cogdx score of 1, and MMSE scores ≥ 27. The AD cohort included participants with Braak scores of 4–6, CERAD scores of 1–2, cogdx scores of 4–5, and MMSE scores ≤ 14. The final dataset comprised prefrontal cortex samples from 35 participants, including 17 healthy controls and 18 AD cases, and contained 163,411 snRNA-seq nuclei and 38,646 snATAC-seq nuclei after retaining those with transcription start site (TSS) enrichment > 6 and 1,000–100,000 fragments. snATAC-seq data was analysed using the ArchR package^94^. The analysis included iterative LSI dimension reduction and clustering using a 500 bp tile matrix, with parameters “iterations = 3, resolution = 0.2, varFeat = 35000”, followed by UMAP visualisation. Peak calling was performed using MACS2, with maxPeaks set to 300,000 and a q-value threshold 0.01. Transcription factor motif annotations were added to the snATAC-seq peak set with the CIS-BP motif database, followed by motif deviation score calculation using *addDeviationsMatrix* function. snATAC-seq nuclei were integrated with the snRNA-seq reference using ArchR’s *addGeneIntegrationMatrix* function.

Cell type-specific differentially accessible marker genes were identified using nuclei from healthy donors. ArchR’s *getMarkerFeatures* function was applied to the GeneScoreMatrix, adjusting for TSS enrichment and library size, and genes were retained using FDR < 0.05 and log_2_FC > 2 thresholds. To identify AD-associated changes in chromatin accessibility, the analysis was repeated using the full 35-donor dataset, comparing AD nuclei against healthy nuclei within each cell type while adjusting for the same technical covariates. Differential gene score results were extracted for each cell type, and genes with FDR < 0.05 were classified as showing increased or decreased inferred accessibility in AD according to the direction of log_2_FC.

Cell type-specific TF activity was inferred using healthy donor data. Differential motif activity across cell types was assessed using *getMarkerFeatures* on the chromVAR^95^-derived MotifMatrix, adjusting for TSS enrichment and library size. To link motif activity with TF accessibility and expression, ArchR’s *correlateMatrices* function was used to correlate motif deviation scores (MotifMatrix) with chromatin accessibility-derived gene scores (GeneScoreMatrix), and gene scores (GeneScoreMatrix) with snRNA-seq-derived integrated gene expression values (GeneIntegrationMatrix). Candidate cell type-specific TFs were retained if they met the following criteria:

1. Significant differential motif activity in the relevant cell type (FDR < 0.05);
2. Absolute motif activity-gene score correlation > 0.6 with FDR < 0.05;
3. Gene score-gene integration correlation > 0.6 with FDR < 0.05;
4. Increased expression in the corresponding snRNA-seq cell type marker analysis (log_2_FC > 1 with FDR < 0.05)

TFs satisfying all four criteria were classified as cell type marker TFs if they showed either:

1. increased motif activity in that cell type (activator TFs) together with a positive correlation between motif activity and chromatin accessibility-derived gene scores, or
2. decreased motif activity in that cell type (suppressor TFs) together with a negative correlation between motif activity and chromatin accessibility-derived gene scores.

To infer cell type-specific differences in TF activity between AD and healthy control donors, the full 35-donor dataset was used. The analysis was performed for each cell type individually and involved the following steps with the corresponding cutoffs:

1. differential motif activity between AD and healthy nuclei was calculated using *getMarkerFeatures* on the chromVAR-derived MotifMatrix, adjusting for TSS enrichment and library size, retaining TFs with FDR < 0.05;
2. Pearson correlation between motif activity and ATAC-inferred gene scores was calculated using *correlateMatrices* function, and TFs with absolute correlation > 0.6 and FDR < 0.05 were retained;
3. Pearson correlation between gene scores and snRNA-seq-derived integrated gene expression values was calculated using *correlateMatrices*, retaining TFs with correlation > 0.6 and FDR < 0.05.

TFs that met all 3 criteria in a given cell type were classified as showing increased activity in AD if they had:

1. increased motif activity (activator TFs) together with a positive motif activity-gene score correlation, or
2. decreased motif activity (suppressor TFs) together with a negative motif activity-gene score correlation.

Otherwise, they were deemed to be more active in healthy controls.

For each differentially active TF identified above, its positive and negative targets were obtained from the CollecTRI^76^ regulatory network used to generate separate positive and negative TF target signatures.

### Tau interactomes

Tau interactomes across different contexts were reported by Tracy et al., 2022^29^, and downloaded from the Supplementary Materials of the original study. The authors used ascorbic acid peroxidase (APEX) methodology combined with a proximity ligation assay to identify proteins interacting with the N– and C-termini of wild-type tau, as well as activity-dependent N– and C-terminal tau interactomes and interactomes associated with the P301L and V337M tau mutants. Interacting proteins were defined using the stringent inclusion criteria described in the original study.

### PSEN1 and APP interactomes

PSEN1 and APP interactomes were reported by Hosp et al., 2015^30^, and obtained from the Supplementary Materials of the original study. The authors used quantitative affinity purification coupled to mass spectrometry to identify interaction partners of PSEN1 and APP. Interacting proteins were defined according to the stringent inclusion criteria described in the original study.

### CRISPRi studies

We obtained CRISPR interference (CRISPRi) screen results from Tian et al., 2021^26^; Leng et al., 2022^24^; Dräger et al., 2022^23^; Bravo et al., 2024^27^; and Samelson et al., 2026^25^. Hit genes across all studies were defined as those with FDR < 0.05. We included CRISPRi screens designed to identify genetic inducers and suppressors of the following AD-relevant cellular phenotypes:

- Tian et al., 2021: neuronal cell death, neuronal oxidative stress, and neuronal lipid peroxidation (genome-wide screens);
- Leng et al., 2022: astrocyte phagocytosis and astrocyte inflammation (druggable genome screens of 2,318 genes);
- Dräger et al., 2022: microglia inflammation and microglia phagocytosis (druggable genome screens of 2,325 genes);
- Bravo et al., 2024: seeding-induced tau propagation (a targeted screen of 1,037 genes);
- Samelson et al., 2026: WT and mutant tau-V337M oligomerisation (a targeted screen of 1,037 genes).

### Substrates of γ-secretase

The substrates of γ-secretase were reported by Hou et al., 2023^28^, and downloaded from the Supplementary Materials of the original study. The authors used mass spectrometry-based G-SECSI method to identify membrane-bound γ-secretase substrates in human induced pluripotent stem cell (hiPSC)-derived microglia. The criteria for defining proteins as γ-secretase substrates are described in the original study.

### GWAS

Three gene signatures were derived from Alzheimer’s disease GWAS results reported by Kunkle et al., 2019^31^; Wightman et al., 2021^32^; and Bellenguez et al., 2022^34^. A fourth GWAS-derived signature was generated from Alzheimer’s disease associations downloaded from the NHGRI-EBI GWAS Catalog^48^ (https://www.ebi.ac.uk/gwas), retaining genes with genome-wide significant associations (P < 5 x 10^-8^) that were reported in at least three GWAS studies.

### Comparative Toxicogenomics Database

The Comparative Toxicogenomics Database (CTD)^36^ was accessed on 20 June 2024 to retrieve genes associated with Alzheimer’s disease using the MeSH disease identifier D000544. Only genes with direct supporting evidence were retained in the final signature.

### DisGeNet

The DisGeNet database^37^ was accessed on 3 June 2024 to retrieve genes associated with Alzheimer’s disease using the disease identifier C0002395. The resulting signature was restricted to curated gene-disease associations.

### Drug lists

Alzheimer’s disease clinical trial data were downloaded from DrugBank^49^, focusing on trials investigating potential therapeutic interventions rather than diagnostic, preventative or supportive-care approaches. Trials were further filtered to retain only those that had progressed beyond phase 1 and to exclude studies that had been withdrawn, terminated or suspended. For the retained trials and associated drugs, drug targets were identified using DrugBank’s drug-target interaction annotations, restricting to interactions marked as having pharmacological action. For drugs with missing target information, targets were identified through manual literature review. All unique targets were retained, with one exception: for targets belonging to multi-protein complexes, only one representative subunit was included in the final gene signature to avoid over-representation of such complexes. Additional drug candidates were manually curated from clinicaltrials.gov and alzforum.org^50^.

