## Supplementary Table 1 for "BRIDGE-AD reveals Alzheimer’s disease effectors through interpretable large-scale omics integration"

**Table 1. Class 1 Candidates.**

| Gene | Primary cell type | Dominant pathway | Intervention | Protective or detrimental | Neuro-pathology assessed? | Behavioral/ Cognitive function assessed? | Model system | Clinical develop-ment? | GWAS gene? |
|---|---|---|---|---|---|---|---|---|---|
| ADAM10 | Neurons | Core AD machinery | NA | NA | NA | NA | NA | | |
| ANXA1[1] | Endothelial cells | Cerebrovascular integrity | Recombinant protein | Protective | Yes | Yes | 5xFAD, TauP301L mice | | |
| VGF[2] | Neurons | Synaptic integrity | OE | Protective | Yes | Yes | 5xFAD mice | | |
| CX3CL1[3] | Neurons | Neurogenesis | OE | Protective | Yes | Yes | TauP301S mice | | |
| MDK[4] | Astrocytes | Amyloid aggregation | KO | Protective | Yes | No | 5xFAD mice | | |
| CD2AP[5,6] | Neurons, microglia | MAPK signaling, synapse phagocytosis | cKO | Protective/ Detrimental | Yes | Yes | APP/PS1, 5xFAD mice; SH-SY5Y cells | | Yes |
| IGF1R[7] | Neurons | Insulin/IGF signaling | cKO | Detrimental | Yes | Yes | APP/PS1 mice | | |
| TMEM119[8] | Microglia | Aβ clearance / phagocytosis | OE, Pharm activation | Protective | Yes | Yes | 5xFAD mice | | |
| MS4A4A[9,10] | Microglia | Aβ clearance, phagocytosis, lysosome, TREM2 regulation | KO, OE, Ab | Detrimental | Yes | No | 5xFAD, AppNL-G-F Rag2-/- Il2rγ-/- hCSF1KI mice; primary human macrophages, hiPSC-derived microglia; non-human primates | | Yes |
| MS4A6A[11,10] | Microglia | Aβ clearance / phagocytosis, neuroinflammation | KO, OE | Protective | Yes | Yes | APP/PS1 mice; HMC3 cells | | Yes |
| EBP1[12] | Neurons | APP processing / secretase | cKO, OE | Protective | Yes | Yes | 5xFAD, Ebp1-CKO mice; human postmortem | | |
| SYT7[13] | Microglia | Neuroinflammation | OE | Protective | Yes | Yes | APP/PS1 mice; BV2 microglia cells | | |
| AP2A1[14] | Neurons | Autophagy, neuronal trafficking | OE | Protective | Yes | Yes | APP/PS1 mice; N2a/APPswe cells | | |

| Gene | Primary cell type | Dominant pathway | Intervention | Protective or detrimental | Neuro-pathology assessed? | Behavioral/ Cognitive function assessed? | Model system | Clinical develop-ment? | GWAS gene? |
|---|---|---|---|---|---|---|---|---|---|
| CTSS[15] | Neurons | Neuroinflammation, neuron-microglia cross-talk | KD, OE, inhibitor | Detrimental | Yes | Yes | APP/PS1, WT mice; BV2 cells, HT-22 cells | | |
| GNB5[16] | Neurons | APP processing / secretase | cKO, LOF, OE | Protective | Yes | Yes | 5xFAD mice | | |
| METTL3[17] | Neurons | APP processing / secretase | cKO, inhibitor | Detrimental | Yes | Yes | 5xFAD mice; human postmortem | | |
| CREB1[18] | Neurons, microglia | Neuroinflammation, neuronal resilience | KO, inhibitor | Protective | Yes | Yes | APP/PS1 mice; human iPSC-N cells / iMG cells | | |
| HRH4[19] | Microglia | Phagocytosis | Inhibitor | Detrimental | Yes | Yes | 3xTg-AD, APPswe/PSEN1dE9 mice | | |
| NTN1[20] | Neurons | Neuronal apoptosis | KO, KD, LOF, inhibitor, Ab | Protective | Yes | Yes | APP/PS1, 3xTg, Netrin-1flox/flox mice; primary neurons | | |
| UQCRC1[21] | Neurons | Lysosome function, mitochondria bioenergetics | cKO, OE | Protective | No | Yes | APP/PS1 mice; primary neurons | | |
| LGALS9[22] | Microglia | Microglia activation, neuroinflammation | KO | Detrimental | Yes | Yes | TauP301S mice | | |
| DOR[23] | Microglia | Complement signaling | OE, agonist | Protective | Yes | Yes | APP/PS1 mice; BV2 cells, HT22 cells | | |
| CDK3[24] | Neurons | Cell cycle, cell death | cOE, inhibitor | Detrimental | Yes | Yes | 5xFAD, APP/PS1, Tau-PS19 mice | | |
| GPR120[25] | Microglia | mTORC1 signaling | cKO, GOF | Protective | Yes | Yes | 5xFAD, APP/PS1 mice | | |
| HVCN1[26] | Microglia | Mitochondrial bioenergetics | Inhibitor | Detrimental | Yes | Yes | Hvcn1–/–, 3×Tg-AD mice; BV2 cells, primary neurons and microglia | | |
| SOX9[27] | Astrocytes | Phagocytosis | cKO, OE | Protective | Yes | Yes | APP−NLGF mice | | |

| Gene | Primary cell type | Dominant pathway | Intervention | Protective or detrimental | Neuro-pathology assessed? | Behavioral/ Cognitive function assessed? | Model system | Clinical develop-ment? | GWAS gene? |
|---|---|---|---|---|---|---|---|---|---|
| MFE2[28] | Microglia | Fatty acid β-oxidation, mitochondrial bioenergetics | cKO, activator | Protective | Yes | Yes | 5xFAD mice; primary microglia, BV2 cells | | |
| ACE[29] | Microglia | Aβ clearance / phagocytosis, endolysosomal trafficking | cOE | Protective | Yes | Yes | 5xFAD, Cx3Cr1+/CreERT2; Rosa+/hACE-loxP, ACE10/10 mice | Yes | Yes |
| HAVCR2[30] | Microglia | TGFβ signaling, microglia homeostasis | cKO | Detrimental | Yes | Yes | 5xFAD mice; primary microglia, iPSC-MG cells | | Yes |
| C5AR1[31,32,33] | Microglia | Complement cascade | KO, OE, inhibitor | Detrimental | Yes | Yes | Arc, Tg2576 mice | | |
| PILRA[34] | Microglia | Lipid metabolism, neuroinflammation, bioenergetics | KO, OE, Ab | Detrimental | Yes | No | 5xFAD-hCSF1 mice; hiMG cells | | Yes |
| PHD3[35] | Microglia | Type I interferon signaling | KO, cOE, inhibitor | Detrimental | Yes | Yes | APP-PSEN1/+ mice | | |
| TGR5[36] | Neurons | APP processing, secretase | KO, cKO | Detrimental | Yes | Yes | 5xFAD, CamKIIα,Tgr5F/F 5×FAD, Gad2,Tgr5F/F 5×FAD mice | | |
| CALHM2[37] | Microglia | Calcium homeostasis, neuroinflammation, phagocytosis | KO, cKO | Detrimental | Yes | Yes | 5xFAD mice | | |
| TRIM11[38] | Neurons | Tau protein quality control | KO, OE | Protective | Yes | Yes | TauP301S, 3×Tg-AD mice; SH-SY5Y cells, Neuro-2a cells, primary neurons | | |
| IDO1[39] | Astrocytes | Glucose metabolism | Inhibitor | Detrimental | Yes | Yes | 5xFAD, APP/PS1, TauP301S mice; primary mouse astrocytes, hiPSC-AST cells, hiPSC-N cells | | |

| Gene | Primary cell type | Dominant pathway | Intervention | Protective or detrimental | Neuro-pathology assessed? | Behavioral/ Cognitive function assessed? | Model system | Clinical develop-ment? | GWAS gene? |
|---|---|---|---|---|---|---|---|---|---|
| PHGDH[40] | Astrocytes | NF-κB, mTOR, autophagy | KD, LOF, cOE, inhibitor | Detrimental | Yes | Yes | 3xTg-AD, 5xFAD, APP KI mice; brain organoids | | |
| CCKBR[41] | Neurons | APP processing / secretase, calcium signaling | Agonist | Protective | Yes | Yes | 5xFAD mice | | |
| DGAT2[42] | Microglia | Lipid metabolism, phagocytosis | Inhibitor | Detrimental | Yes | No | 5xFAD mice | | |
| ADGRG1[43] | Microglia | Microglia homeostasis, Aβ phagocytosis | cKO | Protective | Yes | Yes | 5xFAD mice; hESC-MG cells | | |
| ZBP1[44] | Microglia | Neuroinflammation | KO, cKO, inhibitor | Detrimental | Yes | Yes | 5xFAD mice | | |
| CD33[45] | Microglia | TREM2 signaling | KO | Detrimental | Yes | Yes | 5xFAD mice | | Yes |
| CASP1[46,47] | Neurons, microglia | Neuronal resilience and survival | KO, inhibitor | Detrimental | Yes | Yes | J20, APP/PS1 mice; human primary neurons | | |
| PPARA[48-51] | Astrocytes, microglia, neurons | Aβ clearance, autophagy | cKO, Pharm activation | Protective | Yes | Yes | 5xFAD, APP-PSEN1ΔE9, APP/PS1 mice; primary astrocytes, primary human microglia, SH-SY5Y cells | | |
| APOE[52-54] | Astrocytes, microglia | Lipid metabolism | NA | NA | NA | NA | NA | NA | Yes |
| CSF2[55-57] | Microglia | Aβ clearance, microglia homeostasis | Recombinant protein | Protective | Yes | Yes | TgF344-AD rats, clinical trial | Yes | |
| TREM2[58-61] | Microglia | Microglial activation / phagocytosis | NA | NA | NA | NA | NA | | |
| ACHE[62-65] | Neurons | Cholinergic neurotransmission | Inhibitor | Detrimental | Yes | Yes | APdE9 mice, 5xFAD, APP/PS1 | Yes | |
| NGFR[66,67] | Astrocytes | Neurogenic signaling | OE | Bidirectional | Yes | Yes | APP/PS1dE9 mice; primary human astrocytes; clinical trial | Yes | |

| Gene | Primary cell type | Dominant pathway | Intervention | Protective or detrimental | Neuro-pathology assessed? | Behavioral/ Cognitive function assessed? | Model system | Clinical develop-ment? | GWAS gene? |
|---|---|---|---|---|---|---|---|---|---|
| C3[68,69,70] | Microglia | Complement cascade | KO | Detrimental | Yes | Yes | APP/PS1, PS2APP, TauP301S mice | | |
| HMGCR[71-73] | Neurons, endothelial cells | Wnt/β-catenin signaling, vascular integrity, mitochondrial function | Inhibitor | Detrimental | Yes | Yes | APP mice; Wistar rats | Yes | |
| CLU[74,75,76] | Astrocytes, endothelial cells | Immune/neuroinflammatory signaling | KO, LOF, GOF, recombinant protein | Bidirectional | No | No | APP, 5xFAD mice; hiPSC-AST-N-MG co-cultures | | Yes |
| IL10[77] | Microglia | Neuroinflammation, Aβ clearance | KO, recombinant protein | Detrimental | Yes | Yes | APP/PS1 mice; primary microglia | | |
| PRNP[78] | Neurons | Tau aggregation and neurotoxicity | KO, KD, Ab | Detrimental | Yes | No | Primary neurons, iPSC-N cells | | |
| PDCD1[79,80] | Microglia, T cells, astrocytes | Immune/neuroinflammatory signaling, Aβ clearance | KO, Ab | Detrimental | Yes | No | APP/PS1-21, 5xFAD, TE4 mice | | |
| LRP1[81] | Neurons | Tau uptake and spread | KD | Detrimental | Yes | No | WT mice; H4 neuroglioma cells, hiPSC-N cells | | |
| CCR7[82] | T cells | Immune/neuroinflammatory signaling | KO, Ab | Protective | Yes | Yes | 5xFAD, WT mice | | |
| BDNF[83-85] | Neurons | Neuroinflammation, APP processing / secretase | KO, KD, OE, Pharm activation, Ab | Protective | Yes | Yes | TauP301L mice; rat primary neurons, primary neurons; clinical trial | Yes | |
| ABCA1[86] | Microglia, astrocytes | Lipid metabolism | OE, Pharm activation | Protective | Yes | Yes | TauP301S, TE4, TEKO, TE3 mice | | Yes |
| HLA-A[87] | Neurons | Tau processing | KO, KD | Detrimental | Yes | No | TauP301S mice; primary neurons | | |
| PSEN2[88,89] | Neurons | γ-secretase / Aβ generation | NA | NA | NA | NA | NA | | |
| IL3[90] | Astrocytes, microglia | Microglial activation | KO, recombinant protein | Protective | Yes | Yes | 5xFAD mice; hiPSC-MG-AST-N co-cultures | | |

| Gene | Primary cell type | Dominant pathway | Intervention | Protective or detrimental | Neuro-pathology assessed? | Behavioral/ Cognitive function assessed? | Model system | Clinical develop-ment? | GWAS gene? |
|---|---|---|---|---|---|---|---|---|---|
| MAOB[91,92] | Neurons, astrocytes | APP processing / secretase, astrogliosis | KD, OE, inhibitor | Detrimental | No | Yes | APP/PS1 mice; primary neurons | Yes | |
| ITGAL[93] | Neutrophils | Leukocyte adhesion & transmigration | KO, Ab | Detrimental | Yes | Yes | 5xFAD, 3xTg-AD mice | | |
| SLC2A1[94] | Endothelial cells | Vascular integrity | KO, cKO | Protective | Yes | Yes | APPSw/0 mice | | |
| PSEN1[88,89] | Neurons | γ-secretase / Aβ generation | NA | NA | NA | NA | NA | | |
| NLRP3[95,47] | Microglia | Neuroinflammation | KO | Detrimental | Yes | Yes | APP/PS1, Tau22 mice; primary microglia, primary neurons | | |
| PPARG[96,97] | Neurons | Neuronal energy metabolism | Pharm activation | Protective | Yes | Yes | Clinical trial, WT rats | Yes | |
| APP[88,89] | Neurons | Aβ generation | NA | NA | NA | NA | NA | | |
| FKBP1A[98] | Neurons | Tau processing | KD, OE | Protective | Yes | No | Primary neurons, hiPSC-N-AST cells | | |
| GRM5[99,100] | Neurons | Synaptic transmission | KO, inhibitor | Detrimental | Yes | Yes | APPswe/PS1ΔE9, 3xTg-AD mice | | |
| RAB5A[101] | Neurons | Endosomal trafficking | cOE | Detrimental | Yes | Yes | WT mice | | |
| EPHX2[102,103] | Hepatocytes, microglia, astrocytes | Aβ metabolism, neuroinflammation | cKO, cKD, cOE, inhibitor | Detrimental | Yes | Yes | 5xFAD, 3xTg-AD mice | | |
| CXCL10[104] | T cells, microglia | Neuroinflammation | Inhibitor, Ab, recombinant protein | Detrimental | No | No | hiPSC-MG-N-AST co-cultures, primary human T cells | | |
| HMGB1[105] | Astrocytes | Cellular senescence, neuroinflammation | inhibitor | Detrimental | Yes | Yes | hTau mice | | |
| INPP5D[106,107,108] | Microglia | TGFβ signaling, microglia activation, Aβ phagocytosis | KO, cKO | Detrimental | Yes | Yes | APP/PS1, 5xFAD mice | | Yes |

| Gene | Primary cell type | Dominant pathway | Intervention | Protective or detrimental | Neuro-pathology assessed? | Behavioral/ Cognitive function assessed? | Model system | Clinical develop-ment? | GWAS gene? |
|---|---|---|---|---|---|---|---|---|---|
| RYR2[109,110,111,112] | Neurons | Calcium signaling, autophagy, lysosome function | LOF, GOF, Pharm activation, inhibitor, hiPSC-N | Detrimental | Yes | Yes | 3xTg-AD, 5xFAD, APP/PS1, WT mice; primary neurons | | |
| RIPK1[113] | Microglia | Microglia activation, Aβ phagocytosis | LOF, inhibitor | Detrimental | Yes | Yes | APP/PS1 mice; primary microglia | | |
| PICALM[114,115] | Astrocytes, endothelial cells | Endosomal trafficking, vascular integrity, Aβ metabolism | KD, haploinsufficiency, OE, cOE | Protective | Yes | No | WT, APPsw/0 mice; hiPSC-AST cells, primary human brain endothelial cells, hiPSC-END cells; yeast | | Yes |
| SORL1[116] | Neurons | Endosomal function | Haploinsufficiency | Protective | Yes | No | Göttingen minipigs | | Yes |
| CD22[117] | Microglia | Phagocytosis | KO, Ab | Detrimental | No | Yes | WT mice; BV2 cells | | |
| FLNA[118,119] | Neurons | Protein aggregation | Inhibitor | Detrimental | Yes | Yes | Clinical trial | Yes | |
| BACE1[88,89,120,121] | Neurons | β-secretase / Aβ generation | NA | NA | NA | NA | NA | | |
| BIN1[122,123,124] | Neurons | Tau processing, endocytosis | cKO, KD | Protective/Detrimental | Yes | No | TauP301S mice; rat primary neurons | | Yes |
| MAPT[88,89,125,126] | Neurons | Tau stability / aggregation | NA | NA | NA | NA | NA | | |
| BCHE[65,127,128] | Neurons | Cholinergic neurotransmission | KO, inhibitor | Detrimental | Yes | Yes | APP/PS1, 5xFAD | Yes | |
| NUMB[129] | Neurons | Neuronal integrity | cKO | Protective | Yes | No | TauP301S mice | | |
| PKM[130] | Microglia | Glucose metabolism | cKO, KD, inhibitor | Detrimental | Yes | Yes | 5xFAD mice; BV2 cells | | |
| ATG16L1[131] | Microglia | Aβ processing | LOF | Protective | Yes | Yes | WT mice | | |
| MFGE8[132] | Endothelial cells | Vascular integrity, Aβ aggregation | KO | Detrimental | Yes | No | APP/PS1, APP23 mice | | |
| FGB[133] | Microglia | Immune/neuroinflammatory signaling | Ab | Detrimental | Yes | No | 5xFAD mice; primary macrophages, human | | |

| Gene | Primary cell type | Dominant pathway | Intervention | Protective or detrimental | Neuro-pathology assessed? | Behavioral/ Cognitive function assessed? | Model system | Clinical develop-ment? | GWAS gene? |
|---|---|---|---|---|---|---|---|---|---|
| | | | | | | | primary macrophages, rat primary neurons | | |
| SCN8A[134,135,136,137] | Neurons | Neuronal/synaptic activity | KD, inhibitor | Detrimental | Yes | Yes | APP/PS1, 5xFAD mice; SH-SY5Y cells | | |
| SCN1A[134,135,136,137] | Neurons | Neuronal/synaptic activity | Haploinsufficiency, inhibitor | Protective/Detrimental | Yes | Yes | WT, J20, 5xFAD mice | Yes | |
| C9orf72[138] | Microglia | Microglia activation | KO | Protective | Yes | Yes | WT, 5xFAD mice | | |
| C1QA[139,140,141,142] | Microglia | Complement cascade | KO, Ab | Detrimental | Yes | No | TauP301S, WT, APP/PS19 mice; primary rat neurons, primary rat microglia | | |
| GRIN1[143] | Neurons | Glutamatergic transmission | Inhibitor | Detrimental | NA | NA | FDA-approved therapeutic | Yes | |
| GLP1R[144-147] | Neurons | Glucose metabolism | Pharm activation | Protective | Yes | Yes | 3xTg mice; HT22 cells; clinical trial | Yes | |
| WASF1[148] | Neurons | Aβ processing | KD, haploinsufficiency, OE | Detrimental | Yes | Yes | 2×Tg mice; N2a-APPswe.PS1ΔE9 cells | | |
| HK2[149] | Microglia | Glucose metabolism, microglia phagocytosis | cKO, inhibitor | Detrimental | Yes | Yes | 5xFAD mice; primary mouse microglia, hiPSC-MG cells | | |
| CGAS[150,151,152] | Microglia | Immune/neuroinflammatory signaling | KO, cOE, GOF, inhibitor | Detrimental | Yes | Yes | WT, TauP301S, 5xFAD mice | | |
| TIA1[153,154] | Neurons | Tau processing | KO, KD, haploinsufficiency, OE | Detrimental | Yes | Yes | TauP301S, tau-/- mice; primary neurons | | |
| HNRNPA2B1[155] | Neurons | Tau processing | KD | Detrimental | Yes | No | TauP301S mice; primary neurons | | |
| IL17F[156] | Neutrophils, microglia | Immune signaling | cKO, Ab, recombinant protein | Detrimental | Yes | Yes | Ragγc female, APP/PS1, 5xFAD-E4 mice | | |
| PLD3[157] | Neurons | Endosomal trafficking, neuronal function | cKO, OE, GOF | Detrimental | No | No | 5xFAD mice | | |

| Gene | Primary cell type | Dominant pathway | Intervention | Protective or detrimental | Neuro-pathology assessed? | Behavioral/ Cognitive function assessed? | Model system | Clinical develop-ment? | GWAS gene? |
|---|---|---|---|---|---|---|---|---|---|
| FNDC5[158, 159] | Astrocytes, neurons | Aβ clearance, cAMP-PKA-CREB signaling | KD, OE, Ab, recombinant protein | Protective | Yes | Yes | WT, APP/PS1ΔE9 mice; ReNcell VM human neural stem cells, primary neurons | | |
| VPS35[160,161,162] | Neurons | Autophagy, tau processing, endosomal trafficking | cKO, KD, OE | Protective | Yes | No | WT mice; hESC-N cells | | |
| ADCYAP1R1[163] | Neurons | Proteasome activity, tau processing | Recombinant protein | Protective | Yes | Yes | rTg4510 mice; primary neurons | | |
| CH25H[164] | Microglia | Lipid metabolism, neuroinflammation | KO | Detrimental | Yes | No | TauP301S mice | | |
| AHSA1[165] | Neurons | Tau processing | OE, inhibitor | Detrimental | Yes | Yes | rTg4510 mice | | |
| CASP2[166] | Neurons | Tau processing | KD, OE | Detrimental | Yes | Yes | rTg4510, WT mice; primary neurons | | |
| STING1[150,152] | Microglia | Immune/neuroinflammatory signaling | KO, inhibitor | Detrimental | Yes | Yes | WT, 5xFAD mice | | |
| ODC1[167] | Astrocytes | Urea metabolism | KD | Detrimental | Yes | Yes | APP/PS1 mice; primary astrocytes | | |
| MYLIP[168] | Microglia, neurons, astrocytes | Lipoprotein clearance | KO, haploinsufficiency, Pharm activation | Detrimental | Yes | No | WT, APP/PS1 mice; primary microglia, mouse Neuro2a neuronal cells, primary astrocytes, BV2 microglia cells, primary neurons | | |
| RASD2[169] | Neurons | Tau processing | KD, OE, inhibitor | Detrimental | Yes | Yes | rTg4510 mice; hiPSC-N cells, primary neurons, MAPT-P301L hiPSC-N cells, MAPT-V337M hiPSC-N cells | | |
| IFITM3[170] | Neurons, astrocytes | APP processing / secretase | KO, KD | Detrimental | Yes | No | 5xFAD, WT mice; mouse hippocampal neurons, hiPSC-N | | |

| Gene | Primary cell type | Dominant pathway | Intervention | Protective or detrimental | Neuro-pathology assessed? | Behavioral/ Cognitive function assessed? | Model system | Clinical develop-ment? | GWAS gene? |
|---|---|---|---|---|---|---|---|---|---|
| | | | | | | | cells, primary human astrocytes | | |
| BSN[171] | Neurons | Tau aggregation and neurotoxicity | KD, OE, GOF | Detrimental | Yes | Yes | TauP301S, WT mice; hTau-P301L Drosophila | | |
| TOM1[172] | Microglia | Immune/neuroinflammatory signaling | KD, OE | Protective | Yes | Yes | 3xTg-AD mice | | |
| ABI3[173,174] | Microglia | Microglia migration and phagocytosis, Aβ clearance | KO, KD | Protective | Yes | Yes | 5xFAD mice; HMC3 cells | | Yes |
| GPR3[175] | Neurons | APP processing / γ-secretase | KO | Detrimental | Yes | Yes | APP/PS1, APPDutch, AppNL, AppNL-F, WT mice | | |
| TMEM164[176] | Astrocytes | Astrocyte activation and neurotoxicity | OE, cOE | Protective | Yes | Yes | 5xFAD mice; primary astrocytes, primary neurons, human primary astrocytes, hESC-N cells | | |
| PIEZO1[177] | Microglia | Microglia activation, Aβ phagocytosis | cKO, Pharm activation | Protective | Yes | Yes | 5xFAD mice | | |
| SLC9A6[178] | Neurons | Endosomal trafficking | KO, cKO, KD, inhibitor | Detrimental | Yes | No | WT, APOE3 and APOE4, AppNL-F, AppNL-F;ApoeAPOE4 mice; primary neurons | | |
| EXT1[179] | Neurons | Aβ clearance | cKO | Detrimental | Yes | No | APP/PS1 mice | | |
| IL17A[180,181] | T cells | Immune signaling | OE, Ab | Detrimental/ Protective | Yes | Yes | 3xTg-AD, TgAPPswe/PS1dE9 mice | | |
| B2M[182] | Microglia | Aβ aggregation and neurotoxicity | KO, KD, OE, Pharm inhibition, Ab, recombinant protein | Detrimental | Yes | Yes | B2MKI/KI, 5×FAD, 5×FAD; B2MKI/KI mice | | |
| ACE2[183] | Neurons | Renin–angiotensin system, neuromodulation | Pharm activation, Pharm inhibition | Protective | Yes | Yes | Tg2576 mice | | |

| Gene | Primary cell type | Dominant pathway | Intervention | Protective or detrimental | Neuro-pathology assessed? | Behavioral/ Cognitive function assessed? | Model system | Clinical develop-ment? | GWAS gene? |
|---|---|---|---|---|---|---|---|---|---|
| PYCARD[184] | Microglia | Aβ aggregation and neurotoxicity | KO, LOF, Ab, recombinant protein | Detrimental | Yes | Yes | APP/PS1 mice; primary microglia, THP-1 cells | | |
| TYROBP[185,186] | Microglia | Microglia homeostasis and activation, tau clearance | KO | Detrimental | Yes | Yes | APP/PS1, TauP301S mice; primary microglia | | |
| IL-33[187,187] | Microglia | Immune/neuroinflammatory signaling | Recombinant protein | Protective | Yes | Yes | APP/PS1 mice; primary microglia | | |
| SYK[188,189] | Microglia | Microglia homeostasis and activation | cKO, Pharm activation, Ab activation | Protective | Yes | Yes | 5xFAD mice; primary macrophages | | |
| PTK2B[190] | Neurons | Tau processing and neurotoxicity | KO, inhibitor | Protective | Yes | Yes | TauP301S mice; hiPSC-N cells | | Yes |
| CX3CR1[191] | Microglia | Aβ clearance / phagocytosis, microglia homeostasis | KO | Protective | Yes | Yes | 5xFAD mice | | |
| ITGB8[106] | Microglia | TGFβ signaling, microglia activation, Aβ phagocytosis | cKO, Ab | Detrimental | Yes | No | WT, APP/PS1 mice | | |
| TAGLN2[192] | Neurons | Tau processing and neurotoxicity | KD, OE, Pharm activation, peptide activator | Protective | Yes | Yes | TauP301S mice; primary neurons, SH-SY5Y cells | | |
| PLCG2[193,194,195] | Microglia | Immune/neuroinflammatory signaling, microglia activation, lipid metabolism | KO, LOF, GOF | Protective | Yes | No | WT, 5xFAD mice; hiMG cells | | Yes |
| CXCR3[104] | T cells | T cell recruitment | Inhibitor, Ab | Detrimental | No | No | hiPSC-MG-N-AST co-cultures, primary human T cells | | |
| IFNG[196,79] | Microglia, T cells | Immune/neuroinflammatory signaling | cKO, Ab | Detrimental/ Protective | Yes | No | TE3 mice | | |

| Gene | Primary cell type | Dominant pathway | Intervention | Protective or detrimental | Neuro-pathology assessed? | Behavioral/ Cognitive function assessed? | Model system | Clinical develop-ment? | GWAS gene? |
|---|---|---|---|---|---|---|---|---|---|
| ATG7[197] | Microglia | Autophagy | cKO | Protective | Yes | No | 5xFAD mice | | |
| ADORA2A[198,199,200,201] | Neurons | Synaptic plasticity | KD, cOE, inhibitor | Detrimental | Yes | Yes | WT, THY-Tau22, APP/PS1 mice | Yes | |
| TNFRSF1A[202] | Choroid plexus epithelial cells | Inflammation | KO, nanobody | Detrimental | Yes | Yes | APP/PS1, WT mice | | |
| DNM1L[203,204] | Neurons | Mitochondrial dynamics | Inhibitor, inhibitory peptide | Detrimental | Yes | Yes | 5xFAD, APP/PS1 mice; SH-SY5Y cells, N2a APPSwe cells, primary human fibroblasts, human SK-N-MC neuroblastoma cells, primary rat neurons | | |
| EHMT2[205,206] | Neurons | Epigenetic regulation of gene expression | KD, inhibitor | Detrimental | Yes | Yes | TauP301S, 5xFAD mice | | |
| IL2[207,208] | Astrocytes, T cells | Astrocyte activation, immune signaling | OE, recombinant protein | Protective | Yes | Yes | APP/PS1 mice | | |
| TLR2[209,210,211] | Microglia | Neuroinflammation | KO, OE, Ab | Detrimental/ Protective | Yes | Yes | rTg4510, APP/PS1 mice; primary neurons, primary microglia, SH-SY5Y cells, THP-1 cells | | |
| FCGR2B[212,213] | Neurons | Aβ uptake and neurotoxicity | KO, cKO, KD, LOF, OE, Ab, recombinant protein | Detrimental | Yes | Yes | 3xTg-AD mice; primary neurons, primary astrocytes, SH-SY5Y cells | | |
| MYD88[214,215] | Microglia, astrocytes | Immune/neuroinflammatory signaling, Aβ clearance, astrocyte reactivity | cKO, haploinsufficiency | Detrimental | Yes | Yes | APP/PS1 mice | | |

| Gene | Primary cell type | Dominant pathway | Intervention | Protective or detrimental | Neuro-pathology assessed? | Behavioral/ Cognitive function assessed? | Model system | Clinical develop-ment? | GWAS gene? |
|---|---|---|---|---|---|---|---|---|---|
| SPI1[216] | Microglia | Immune signaling, complement activity, microglia homeostasis | KD, haploinsufficiency, OE | Protective | Yes | No | APP/PS1 mice; BV2 cells | | Yes |
| CSF1R[217,218] | Microglia | Microglia homeostasis and activation | Inhibitor, recombinant protein | Detrimental | Yes | Yes | APP/PS1, TauP301S, WT mice; N13 murine microglia cells | Yes | |
| HCK[219,220] | Microglia | Aβ clearance, phagocytosis | KO, inhibitor | Protective | Yes | Yes | Tg2576, J20 mice; primary microglia; BV2 mouse microglia cells | Yes | |
| STAT3[221,222,223] | Astrocytes | Astrogliosis | cKO, OE, inhibitor | Detrimental | Yes | Yes | 5xFAD, APP/PS1, WT mice; primary astrocytes | | |
| IFNAR[224,225] | Microglia | Immune/neuroinflammatory signaling | KO, cKO, Ab, recombinant protein | Detrimental | Yes | Yes | 5xFAD, TauP301S mice; primary neural cells | Yes | |
| PPARGC1A[226] | Neurons | APP processing / secretase | OE | Protective | Yes | Yes | APP23 mice | | |
| TRPM7[227] | Neurons | Aβ processing | LOF, OE | Protective | Yes | Yes | 5xFAD mice; primary neurons | | |

**Abbreviations**: Aβ, amyloid-β; Ab, antibody; AD, Alzheimer's disease; APP, amyloid precursor protein; AST, astrocytes; END, endothelial cells; FDA, U.S. Food and Drug Administration; GOF, gain of function; hESC, human embryonic stem cell; MG, microglia-like cells; hiPSC, human induced pluripotent stem cell; AST, astrocytes; END, endothelial cells; N, neurons; KD, knockdown; cKD, conditional knockdown; KO, knockout; cKO, conditional knockout; LOF, loss of function; OE, overexpression; cOE, conditional overexpression; Pharm activation/inhibition, pharmacological activation/inhibition; WT, wild-type. Common model systems and cell lines include 3xTg-AD, 5xFAD, APP/PS1, APP23, APPNL-G-F, J20, Tg2576, TauP301S, TauP301L, Tau22, THY-Tau22, and rTg4510 transgenic mouse models; BV2, HMC3, HT22, N2a/Neuro-2a, SH-SY5Y, THP-1, and N13 cell lines; and Drosophila, Göttingen minipigs, non-human primates, organoids, primary cells, and clinical trial cohorts where indicated.
