## Supplementary Table 8 for "BRIDGE-AD reveals Alzheimer’s disease effectors through interpretable large-scale omics integration"

**Table S8. Brain tissue donor information.**

| **Donor #** | **Diagnosis** | **Braak score** |
| --- | --- | --- |
| 1 | HC | III |
| 2 | HC | II |
| 3 | HC | II |
| 4 | HC | I |
| 5 | HC | II |
| 6 | HC | 0 |
| 7 | HC | I |
| 8 | HC | I |
| 9 | HC | III |
| 10 | HC | 0 |
| 11 | HC | II |
| 12 | HC | II |
| 13 | HC | III |
| 14 | HC | I |
| 15 | HC | I |
| 16 | HC | II |
| 17 | AD | V |
| 18 | AD | V |
| 19 | AD | V |
| 20 | AD | V |
| 21 | AD | V |
| 22 | AD | V |
| 23 | AD | VI |
| 24 | AD | V |
| 25 | AD | V |
| 26 | AD | VI |
| 27 | AD | VI |
| 28 | AD | VI |
| 29 | AD | V |
| 30 | AD | V |
| 31 | AD | V |
| 32 | AD | V |
| 33 | AD | VI |
| 34 | AD | V |
| 35 | AD | V |
