## Supplementary Table 9 for "BRIDGE-AD reveals Alzheimer’s disease effectors through interpretable large-scale omics integration"

**Table S9. Antibodies used in the study.**

| **Target** | **Antibody** | **Application** | **Vendor** | **Catalog number** | **Working concentration** |
| --- | --- | --- | --- | --- | --- |
| SCARB2 | Rabbit monoclonal | WB, IF | Cell Signaling Technology | 27960 | WB: 1:1000, IF: 1:300 |
| GCase | Mouse monoclonal | WB | Bio-Techne | MAB7410-SP | 1:500 |
| APOE | Goat polyclonal | WB | Millipore | 178479 | 1:1,000 |
| LAMP1 | Mouse monoclonal | WB, IF | Cell Signaling Technology | 15665 | WB: 1:1,000, IF: 1:50 |
| Calnexin | Mouse monoclonal | IF | Proteintech | 66903-1-Ig | 1:300 |
| GAPDH | Mouse monoclonal | WB | GeneTex | GTX627408 | 1:5,000 |
| Rabbit IgG (H+L) | Donkey polyclonal, HRP | WB | Jackson ImmunoResearch | 711-035-152 | Rabbit IgG (H+L) |
| Mouse IgG (H+L) | Donkey polyclonal, HRP | WB | Jackson ImmunoResearch | 715-035-151 | Mouse IgG (H+L) |
| Goat IgG (H+L) | Donkey polyclonal, HRP | WB | Jackson ImmunoResearch | 705-035-147 | Goat IgG (H+L) |
| Rabbit IgG (H+L) | Donkey polyclonal, Alexa Fluor 647 | IF | Jackson ImmunoResearch | 711-605-152 | Rabbit IgG (H+L) |
| Mouse IgG (H+L) | Donkey polyclonal, Alexa Fluor 488 | IF | Jackson ImmunoResearch | 715-545-150 | Mouse IgG (H+L) |
