## Supplementary Table 10 for "BRIDGE-AD reveals Alzheimer’s disease effectors through interpretable large-scale omics integration"

**Table S10. DNA oligonucleotides used in the study.**

| **Oligonucleotide name** | **Sequence** |
| --- | --- |
| SCARB2_ORF_NotI_F | ATCCGCGGCCGCTGTACAAACTGGTGGCAGACCGGTGCTAGCCGCCACCATGGGCCGATGCTGCTTCTA |
| SCARB2_ORF_EcoRI_R | TTCAGAATTCGCATGCTTAGGTTCGAATGAGGGGTG |
| GBA1_ORF_NotI_F | ATCCGCGGCCGCTGTACAAACTGGTGGCAGACCGGTGCTAGCCGCCACCATGGAGTTTTCAAGTCCTTC |
| GBA1_ORF_EcoRI_R | TTCAGAATTCGCATGCTCACTGGCGACGCCACAGGTAG |
| SCARB2_N45Q_mut_F | GAAGAAAATTGTGTTAAGGCAGGGTACTGAGGCATTTGACTC |
| SCARB2_N45Q_mut_R | GAGTCAAATGCCTCAGTACCCTGCCTTAACACAATTTTCTTC |
| SCARB2_N68Q_mut_F | CTCAGTTCTATTTCTTCCAGGTCACCAATCCAGAGGAG |
| SCARB2_N68Q_mut_R | CTCCTCTGGATTGGTGACCTGGAAGAAATAGAACTGAG |
| SCARB2_N105Q_mut_F | CAAATATTCAATTTGGAGATCAGGGAACAACAATATCTGCTG |
| SCARB2_N105Q_mut_R | ACAGCAGATATTGTTGTTCCCTGATCTCCAAATTGAATATTTGC |
| SCARB2_N206Q_mut_F | GGCCTATTCTATGAGAAACAGGGGACTAATGATGGAGACTATG |
| SCARB2_N206Q_mut_R | CATAGTCTCCATCATTAGTCCCCTGTTTCTCATAGAATAGGCC |
| SCARB2_N224Q_mut_F | CTGGAGAAGACAGTTACCTTCAGTTTACAAAAATTGTGGAATG |
| SCARB2_N224Q_mut_R | CATTCCACAATTTTTGTAAACTGAAGGTAACTGTCTTCTCCAG |
| SCARB2_N249Q_mut_F | CAGACAAGTGCAATATGATTCAGGGAACAGATGGAGATTCTTTTC |
| SCARB2_N249Q_mut_R | GAAAAGAATCTCCATCTGTTCCCTGAATCATATTGCACTTGTCTG |
| SCARB2_N304Q_mut_F | CTGCAGAAATATTAGCCCAGACGTCAGACAATGCCGGC |
| SCARB2_N304Q_mut_R | GCCGGCATTGTCTGACGTCTGGGCTAATATTTCTGCAG |
| SCARB2_N325Q_mut_F | CTGGGCTCAGGAGTTCTGCAGGTCAGCATCTGCAAGAAT |
| SCARB2_N325Q_mut_R | ATTCTTGCAGATGCTGACCTGCAGAACTCCTGAGCCCAG |
| SCARB2_N412Q_mut_F | GTTTTCCCAGTGATGTACCTCCAGGAGAGTGTTCACATTGATAAAG |
| SCARB2_N412Q_mut_R | CTTTATCAATGTGAACACTCTCCTGGAGGTACATCACTGGGAAAAC |
| GBA1_N58Q_mut_F | GCTCGGTGGTGTGTGTCTGCCAGGCCACATACTGTGACTCCTTTG |
| GBA1_N58Q_mut_R | CAAAGGAGTCACAGTATGTGGCCTGGCAGACACACACCACCGAGC |
| GBA1_N98Q_mut_F | GTATGGGGCCCATCCAGGCTCAGCACACGGGCACAGGCCTGC |
| GBA1_N98Q_mut_R | GCAGGCCTGTGCCCGTGTGCTGAGCCTGGATGGGCCCCATAC |
| GBA1_N185Q_mut_F | CTGATGATTTCCAGTTGCACCAGTTCAGCCTCCCAGAGGAAG |
| GBA1_N185Q_mut_R | CTTCCTCTGGGAGGCTGAACTGGTGCAACTGGAAATCATCAG |
| GBA1_N309Q_mut_F | CCTAGGTCCTACCCTCGCCCAGAGTACTCACCACAATGTCCG |
| GBA1_N309Q_mut_R | CGGACATTGTGGTGAGTACTCTGGGCGAGGGTAGGACCTAGG |
| qPCR_SCARB2_F | GCCAATACGTCAGACAATGCCG |
| qPCR_SCARB2_R | CTCATCTGCTTGGTAAAAGTGTGG |
| qPCR_GAPDH_F | GTCTCCTCTGACTTCAACAGCG |
| qPCR_GAPDH_R | ACCACCCTGTTGCTGTAGCCAA |
| qPCR_LAMP1_F | CGTGTCACGAAGGCGTTTTCAG |
| qPCR_LAMP1_R | CTGTTCTCGTCCAGCAGACACT |
| qPCR_ATP6V0D1_F | ACAAGACGCTGGAGGACCGATT |
| qPCR_ATP6V0D1_R | ACGATGTTGCGACACTCCTGCT |
| qPCR_APOE_F | GGGTCGCTTTTGGGATTACCTG |
| qPCR_APOE_R | CAACTCCTTCATGGTCTCGTCC |
| qPCR_TREM2_F | ATGATGCGGGTCTCTACCAGTG |
| qPCR_TREM2_R | GCATCCTCGAAGCTCTCAGACT |
| qPCR_ABCA7_F | CACTCTTCCGAGAGCTAGACAC |
| qPCR_ABCA7_R | CTCCATATCTGTGTCCGCAGCA |
| qPCR_NLRP3_F | GGACTGAAGCACCTGTTGTGCA |
| qPCR_NLRP3_R | TCCTGAGTCTCCCAAGGCATTC |
| qPCR_IL1B_F | CTGTCCTGCGTGTTGAAAGA |
| qPCR_IL1B_R | TTGGGTAATTTTTGGGATCTACA |
| qPCR_TNF_F | CAGCCTCTTCTCCTTCCTGAT |
| qPCR_TNF_R | GCCAGAGGGCTGATTAGAGA |
| qPCR_IL6_F | CCTTCCAAAGATGGCTGAAA |
| qPCR_IL6_R | TGGCTTGTTCCTCACTACT |
| qPCR_IL8_F | ATGACTTCCAAGCTGGCCGT |
| qPCR_IL8_R | TCCTTGGCAAAACTGCACCT |
| qPCR_CCL2_F | AGAATCACCAGCAGCAAGTGTCC |
| qPCR_CCL2_R | TCCTGAACCCACTTCTGCTTGG |
| qPCR_CXCL10_F | GGTGAGAAGAGATGTCTGAATCC |
| qPCR_CXCL10_R | GTCCATCCTTGGAAGCACTGCA |
| qPCR_GRP78_F | CTGTCCAGGCTGGTGTGCTCT |
| qPCR_GRP78_R | CTTGGTAGGCACCACTGTGTTC |
| qPCR_GRP94_F | GGAGAGTCGTGAAGCAGTTGAG |
| qPCR_GRP94_R | CCACCAAAGCACACGGAGATTC |
| qPCR_ATF6_F | CAGACAGTACCAACGCTTATGCC |
| qPCR_ATF6_R | GCAGAACTCCAGGTGCTTGAAG |
| qPCR_CHOP_F | GGTATGAGGACCTGCAAGAGGT |
| qPCR_CHOP_R | CTTGTGACCTCTGCTGGTTCTG |
